# traceCB: Trans-ancestry cell-type-specific eQTLs mapping by integrating scRNA-seq and bulk data

**DOI:** 10.64898/2026.06.20.733502

**Authors:** Wenxin Jiang, Jiashun Xiao, Mingxuan Cai

## Abstract

Mapping cell-type-specific expression quantitative trait loci (ct-eQTLs) is essential for interpreting disease-associated variants, yet studies in underrepresented populations are hindered by limited statistical power. Here, we present traceCB, a statistical framework that enhances ct-eQTL mapping in target ancestries by integrating summary statistics from single-cell and bulk-tissue eQTL studies across diverse populations. By explicitly modeling trans-ancestry genetic architecture and accounting for cellular heterogeneity in bulk tissues, traceCB optimizes information borrowing from well-powered European cohorts while robustly controlling for type I error. Simulation studies demonstrate that traceCB achieves superior statistical power compared to original ct-eQTL, particularly when leveraging tissue-level data. In an application to immune cells in East Asian and African cohorts, traceCB increased the effective sample size by up to 2.9-fold and identified approximately 40% more eGenes than single-ancestry analyses, with a replication rate exceeding 90% in independent datasets. Furthermore, traceCB improved the colocalization of regulatory variants with GWAS signals for blood and immune-related traits, revealing cell-type-specific mechanisms underlying complex diseases. These findings establish traceCB as a powerful and scalable tool for leveraging global genomic resources to improve regulatory variant discovery at the cellular level across diverse populations.

## Introduction

Genome-wide association studies (GWASs) have successfully identified associations between hundreds of thousands of genomic variants and complex traits or diseases [1]. Despite these fruitful discoveries, the majority (over 90%) of identified risk variants are located in non-coding regions, posing a significant challenge for deciphering their underlying biological functions [2, 3]. A critical insight is that GWAS signals are enriched in gene regulatory elements, particularly expression quantitative trait loci (eQTLs)—genetic variants associated with gene expression levels [4]. This enrichment underscores the hypothesis that many GWAS hits exert their effects through the regulation of gene expression, as trait-associated SNPs are three times more likely to be eQTLs [5]. While large-scale eQTL studies based on bulk tissues have enhanced our understanding of GWAS signals [6], they rely on average expression strength across all constituent cells in a tissue, which masks the cellular heterogeneity of the eQTL landscape. Growing evidence indicates that eQTL effects can be highly cell-type-specific, with many only detectable in the relevant cell type for a given trait [2, 7, 8], highlighting the critical need to map cell-type-specific eQTLs (ct-eQTLs) that improves our understanding of disease mechanisms at cellular level [9]. Recently, expanding resources of single-cell RNA sequencing (scRNA-seq) data have enabled extensive ct-eQTL studies in various human tissues. Compared with tissue-level bulk RNA-seq, scRNA-seq provides a higher resolution of gene expression profiles at the single-cell level, allowing the identification of ct-eQTLs [10].

Despite the promising insight offered by ct-eQTL mapping, its broad application has been restricted by the lack of generalizability in genetically diversified ancestries. So far, the sample composition of current eQTL studies is highly biased towards European ancestry (EUR) in both bulk and scRNA-seq eQTL data. In the tissue-level Genotype-Tissue Expression (GTEx) v10 dataset, 85% of the 981 donors are from European ancestry, while only 12% are from African descendants, which is the second largest ancestry in GTEx [11]. Similarly, across 11 single-cell eQTL studies from the eQTL Catalogue, 88.6% are of European ancestry, 8.8% are of African ancestry, and only 0.7% are of East Asian ancestry (EAS) [12]. Given the strongly unbalanced sample makeup, most of the eQTL studies have been based on the European subjects. These discoveries may not be extended to non-European populations, limiting their values in interpreting GWAS variants. For example, the heritability of schizophrenia in EAS was found less enriched in the brain eQTLs when they were identified from EUR samples [13]. Therefore, trans-ancestry ct-eQTL mapping, aiming to pinpoint ct-eQTL SNPs across global ancestries, especially for the underrepresented ones, is a critical step toward global health equity.

The challenges of trans-ancestry ct-eQTL mapping arise from two aspects. First, the genetic architecture of eQTLs can vary across ancestries. Recent studies have reported that allele frequency differences account for a substantial proportion of population differences in eQTLs [13]. The eQTL effects and linkage disequilibrium (LD) also display heterogeneous patterns in different ancestries. Second, the ct-eQTL mapping with scRNA-seq data has been restricted by technical constraints. The prohibitive cost of scRNA-seq has greatly restricted the scale of single-cell eQTL studies, limiting the power in ct-eQTL mapping. Furthermore, due to the lower sequencing depth and higher dropout rate, the gene expression levels measured with scRNA-seq are much noisier than the bulk RNA-seq, which can also lead to potential loss of power [14–16].

Unfortunately, existing ct-eQTL methods have only partially addressed the above challenges. To circumvent the noisy nature of scRNA-seq data, interaction-based methods take bulk RNA-seq data as input and detect the ct-eQTLs by modeling the interaction effects between the genetic variants and a proxy marker of cell type, which can be derived from established cell type deconvolution methods [17]. However, these methods can have limited power when working with rare cell types [2, 17]. To better leverage both data types, recent pioneering methods, such as IBSEP [16] and JOBS [18], integrate eQTL summary statistics from both bulk and scRNA-seq data. Although the integrative analysis can improve the statistical power for ct-eQTL prioritization, these methods were designed for single-ancestry analysis and can not account for the heterogeneous genetic architectures across diverse populations. To integrate genetic data from the heterogeneous resources, a number of meta-analysis methods have been developed to combine trans-ancestry genetic studies. The classical fixed-effect (FE) meta analysis model assumes the effect sizes are homogeneous across studies, limiting its applicability in trans-ancestry setting. The random-effects model (RE) [19] and its extension, RE2 [20], relaxes this assumption by allowing the effect sizes to vary. While these methods have been applied to meta analyze trans-ancestry studies, they have not adequately accounted for critical heterogeneity across ancestries, such as the LD and allele frequencies, leading to inflated type-I errors.

In this paper, we propose traceCB, a novel statistical framework designed to enhance the statistical power for ct-eQTL mapping in underrepresented ancestries. Our method achieves this by integrating eQTL summary statistics from scRNA-seq and bulk RNA-seq data across diverse ancestries, thereby leveraging information from a well-powered auxiliary population (Fig. 1). Our framework offers three key advantages over existing methods. First, it explicitly models the local genetic architectures of *cis*-SNPs across ancestries, optimally integrating scRNA-seq data from different populations. Second, it can further utilize high-quality bulk RNA-seq data from European populations to substantially improve the statistical power of ct-eQTL mapping in the target population. Third, the efficient moment-based estimation framework of traceCB operates with summary-level eQTL data, enabling genome-scale analysis. Under this framework, we also introduce a special version, called traceC, that can integrate trans-ancestry ct-eQTL data when bulk data are not available. Through extensive simulations, we demonstrated that traceCB substantially boosts statistical power by borrowing information from trans-ancestry single-cell and bulk eQTL data, while maintaining well-calibrated type I error rates. As a real-world application, we applied traceCB to improve ct-eQTL mapping of 5 major blood cell types in East Asian individuals by integrating BioBank Japan (BBJ) [21] ct-eQTL summary data with 10 European scRNA-seq studies from eQTL Catalogue [12] and whole blood eQTL data from eQTLGen [6]. For genes with significant trans-ancestry co-heritability, traceC increased the effective sample size by 1.82-fold (from 103 to 187.84), whereas traceCB achieved a 2.32-fold increase (from 103 to 239.24, Supplementary Table 8). Averaged across the 10 auxiliary studies (Supplementary Table 9), traceC identified 28% more eGenes across cell types (from 1,929 to 2,463), and traceCB further increased this gain to 38% (from 1,929 to 2,669) as compared to the original BBJ study. We replicated these discoveries in both internal and external validation datasets [22, 23]. High external replication rates of 93% for traceC and 92% for traceCB support the robustness and reliability of our findings (Supplementary Table 9). With colocalization analysis, we showed that ct-eQTLs prioritized by traceCB colocalized with GWAS signals, revealing cell-type-specific gene-regulatory mechanisms underlying blood-related traits and immune diseases. Furthermore, we demonstrated the generalizability of our framework to other ancestries by applying traceCB to integrate African scRNA-seq eQTL data with European studies. Collectively, these results indicate that traceCB substantially enhances the statistical power of trans-ancestry ct-eQTL mapping, helping to bridge the gap between GWAS findings and the underlying cell-type-specific regulatory mechanisms in underrepresented populations.

**Figure 1:**
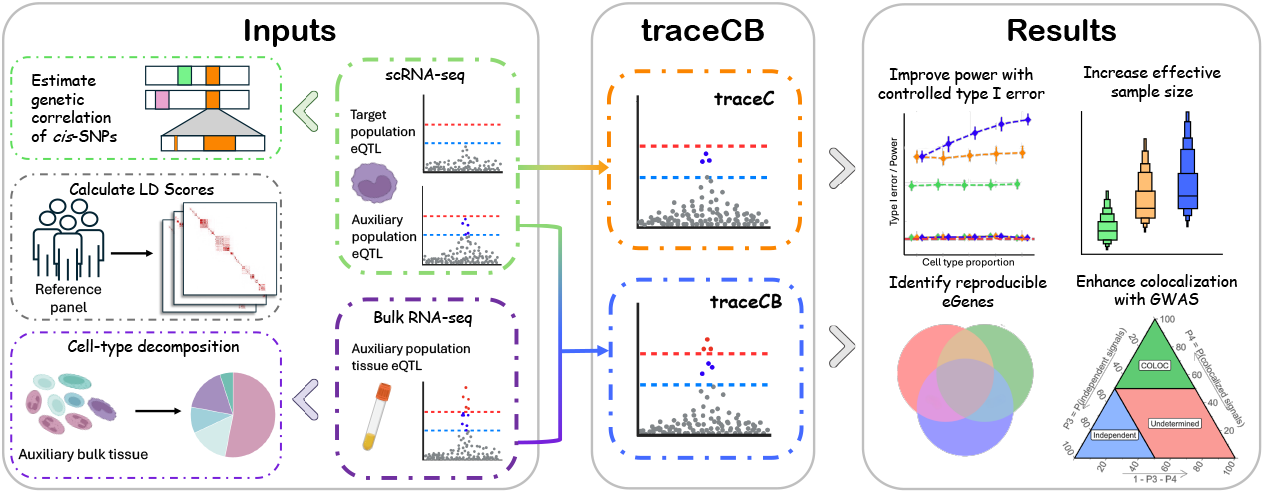
Method overview. traceCB takes summary-level scRNA-seq eQTLs and bulk RNA-seq eQTLs as input. LD scores from a reference panel (e.g. 1000 Genomes Project [26]) and cell-type proportions are required to account for the effects of LD and cell-type composition. It first estimates cell-type-specific heritability and co-heritability across ancestries, and then obtains the improved estimates of eQTL effects. In the absence of bulk tissue data, traceC can be used to prioritize eQTLs without cell-type proportions. Both traceCB and traceC can improve the power of eQTL mapping with controlled type I error rates and empower downstream analyses. Images of blood cells and test tubes come from BioRender.com.

## Results

### Method overview

We present traceCB, a statistical method to prioritize ct-eQTLs in an underrepresented target ancestry by leveraging bulk RNA-seq and scRNA-seq eQTLs from a larger auxiliary ancestry. With the summary statistics from trans-ancestry scRNA-seq eQTLs, traceCB leverages the cell-type-specific *cis*-heritabilities and co-heritabilities of gene expression levels to borrow information across ancestries. Built upon the cell type deconvolution model, traceCB can further leverage the cell-type proportions in the large bulk tissue samples to boost its power. traceCB performs eQTL analysis in a generalized method of moments (GMM) framework, making it adaptive to complex genetic architectures and computationally efficient. The resulting traceCB estimator increases the effective sample size of ct-eQTLs and identifies more eGenes in the target ancestry, facilitating downstream analyses such as colocalization with GWAS or eGene discovery. For scenarios where bulk tissue data is unavailable, we additionally introduce traceC, a simplified version of traceCB that prioritizes eQTLs using only scRNA-seq data. Both methods are computationally efficient and scalable to genome-wide analysis. For instance, in our real data analysis, the processing of chromosome 1 took approximately 432 seconds, corresponding to an average of only 0.3 seconds per gene for both traceC and traceCB. The code of software and analysis process can be found at https://github.com/LucaJiang/traceCB, and the documentation is available at https://lucajiang.github.io/traceCB/.

### Simulation studies: traceCB controls type I error and enhances power in the trans-ancestry setting

We conducted comprehensive simulation studies to evaluate the performance of traceCB under a wide range of genetic architectures and study designs, comparing it against the original summary statistics of eQTL study and the widely-used random effects meta-analysis method (RE2) [20]. We also included traceC, which only leverages trans-ancestry scRNA-seq eQTLs, as a baseline to assess the benefits of incorporating bulk tissue data in traceCB. To mimic real-world scenarios, we utilized genotype data from EAS and EUR ancestries obtained from the UK BioBank (UKBB) cohort [24] and a Chinese study [25]. The LD scores for both ancestries were constructed using the genotypes from the 1000 Genomes Project [26]. We selected *M* = 2, 000 SNPs from Chr 22:44,804,977–46,207,955 bp (GRCh38), treating them as *cis*-SNPs mapped to the target gene. We then randomly sampled *N*_1_ and *N*_2_ individuals from EAS and EUR populations, respectively, to serve as genotypes for scRNA-seq studies, and additionally sampled *N*_*t*_ individuals from EUR as genotypes for bulk eQTL data. The standardized genotype matrices for scRNA-seq in EAS and EUR are denoted as **X**_1_ and **X**_2_, respectively, while the standardized genotype matrix for bulk eQTLs is denoted as **X**_*t*_. To mimic the realistic genetic architectures across ancestries, we considered a sparse setting for the EUR and EAS effect sizes in the target cell type. Specifically, we randomly selected a proportion, denoted as *p*_causal_, of the *cis*-SNPs as causal variants, and simulated their effect sizes from a bivariate normal distribution:

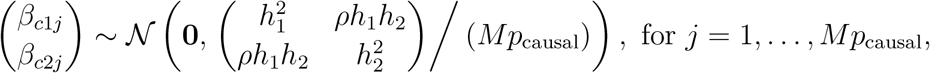

where 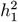 and 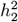 are the heritabilities of ct-eQTL effects in EAS and EUR populations, respectively, and *ρ* is their trans-ancestry genetic correlation. Non-causal variants have zero effect size. To generate the eQTL data of bulk tissue, we considered a simplified scenario incorporating two cell types within the tissue. For the non-target cell type, we similarly selected *p*_causal_ among the *M* SNPs as causal variants and sampled their effect sizes from 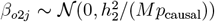. We generated the proportion of the target cell type for *i*-th individual in the bulk tissue samples with 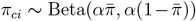, a Beta distribution with shape parameter *α* = 5 to control the variability across samples and 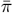 represents the mean proportion of the target cell type. The non-target cell-type proportions are given by *π*_*oi*_ = 1 − *π*_*ci*_. The gene expression levels are generated according to the following linear models:

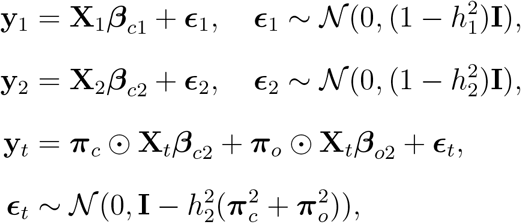

where ***ϵ***_1_, ***ϵ***_2_, and ***ϵ***_*t*_ are independent error terms with variance set to mimic normalized gene expression levels. The eQTL summary statistics were computed by performing marginal regression of the simulated gene expression levels on the genotypes of each SNP. For each setting, we conducted 100 independent simulation replicates to obtain a robust evaluation of model performance. In traceC and traceCB, we estimated the heritability and genetic correlation parameters from the generated summary statistics using LD score regression [27], mimicking the parameter estimation process in real-world applications where individual-level data are unavailable (details provided in Methods). In these simulations, the mean cell-type proportion was set to the true value in order to isolate the statistical behavior of traceCB under controlled tissue-composition settings. This should not be interpreted as assuming that cell-type proportions are observed without error in real applications. In practice, traceCB treats the mean cell-type proportion as an external input, which may be estimated from bulk RNA-seq data using deconvolution methods [18, 28] or obtained from external cell-composition references. For comparison, we implemented two baseline methods: RE2(sc), which performs a random effects meta-analysis of the two scRNA-seq eQTL summary statistics, and RE2(sc+tissue), which extends this approach to additionally incorporate bulk tissue eQTL summary statistics.

To evaluate type I error control under the null hypothesis, we set the heritability of EAS ct-eQTL effects 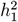 and the trans-ancestry genetic correlation *ρ* to zero, while varying the heritability of EUR ct-eQTL effects 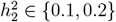 and proportion of causal variants *p*_causal_ ∈ {0.005, 0.01, 0.02}. We also varied the mean cell-type proportion 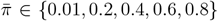 to assess its impact on type I error rates when incorporating bulk tissue data. As shown in Fig. 2 a, both traceCB and traceC consistently maintained type I error rates at the nominal level of 0.05 across all evaluated scenarios, demonstrating robust type I error control. In contrast, the RE2-based methods exhibited severely inflated type I error rates under the null hypothesis. The inflation was exacerbated when the EUR heritability 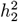 increased (indicating stronger effects in the EUR population) or the proportion of causal SNPs *p*_causal_ was higher (representing denser causal variants). Furthermore, the type I error rate of RE2(sc+tissue) was severely inflated as the proportion of target cell type 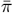 increased. For instance, when 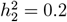 and *p*_causal_ = 0.02, the type I error rate of RE2(sc) reached 0.13. When a tissue-level study with a mean cell-type proportion of 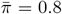 was included, the type I error rate of RE2(sc+tissue) reached 0.4, substantially exceeding the nominal significance level of 0.05. This is because the RE2-based methods rely on the homogeneous-effects assumption that ignores the different genetic architectures across populations, thereby incorrectly borrow the information from the EUR studies.

**Figure 2:**
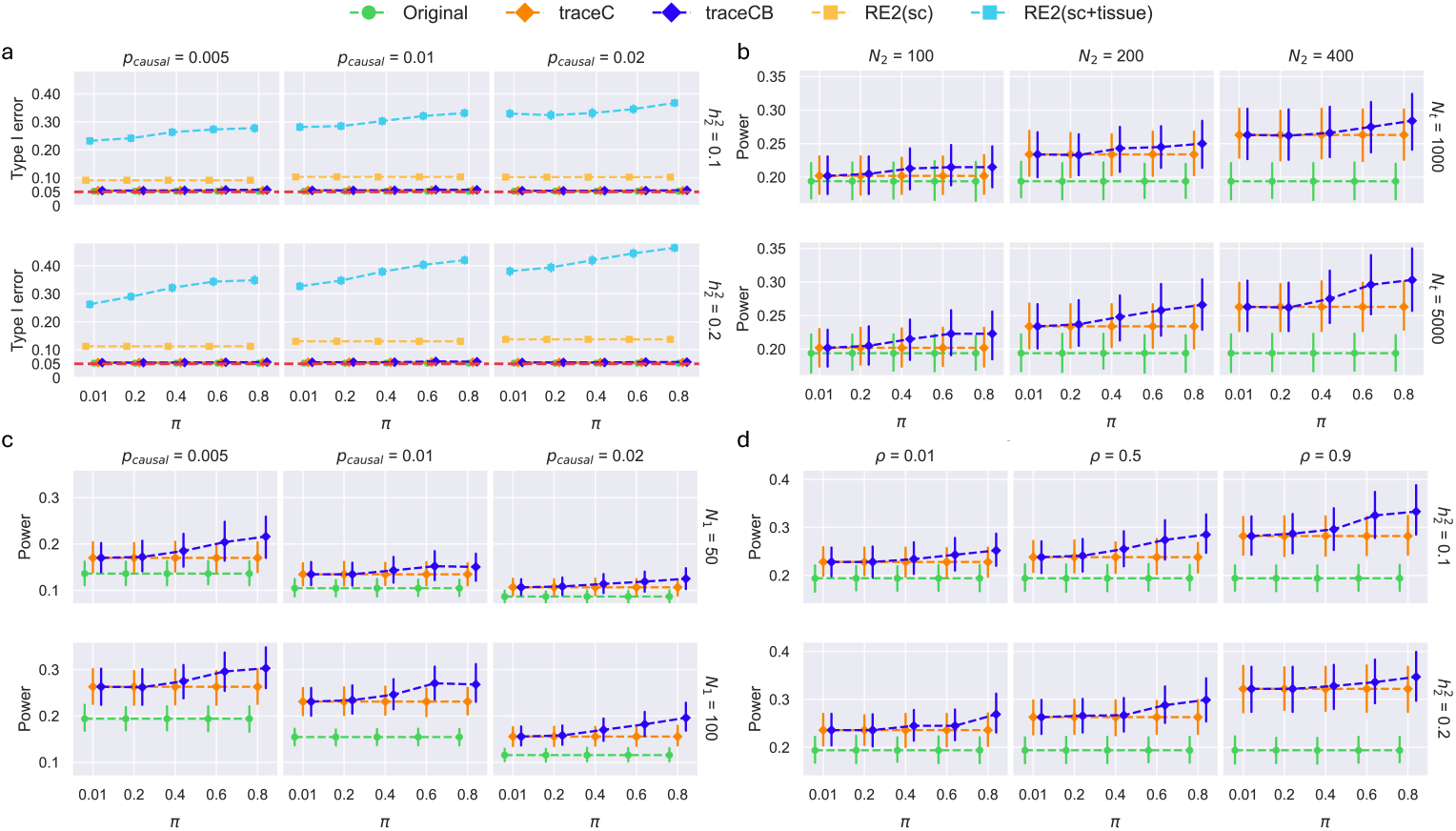
Evaluation of type I error rate and statistical power under simulated scenarios. We compared four methods: “Original”, which uses only the target population’s scRNA-seq summary statistics; “traceC”, a simplified version of traceCB without bulk tissue data; “RE2(sc)”, a random effects meta-analysis of trans-ancestry ct-eQTLs; and “RE2(sc+tissue)”, which extends RE2(sc) to include bulk eQTLs. **a**.Comparison of type I error rate under different *cis*-heritability of the other cell types, fractions of causal SNPs, and mean target cell-type proportion in tissue samples. **b-d**. Comparison of statistical power under different auxiliary sample sizes, proportion of causal SNPs, genetic correlations between the two cell types, *cis*-heritability of the other cell types, and mean target cell-type proportion in tissue samples. All results are averaged over 100 replicates, and error bars represent 95% confidence intervals.

To test whether deconvolution error could induce false-positive discoveries, we repeated the null simulations after perturbing the mean target cell-type proportion supplied to traceCB. TraceCB remained close to the nominal type I error rate across deterministic underestimation/overestimation and random perturbation settings, whereas RE2(sc+tissue) became increasingly inflated as the target cell-type proportion increased (Supplementary Fig. 4). This analysis addresses calibration rather than power or effect-size robustness. The conservative behavior of traceCB reflects its significance-screening step: auxiliary covariance is used only when the estimated trans-ancestry co-heritability is significant; otherwise, the covariance term is set to zero and the estimator effectively reverts to the target-population statistic.

We next considered whether a same-ancestry scRNA-seq-plus-bulk integration strategy could be applied ad hoc in the trans-ancestry setting. Across simulation settings with varying causal-SNP proportions, trans-ancestry genetic correlations and target cell-type proportions, traceC and traceCB maintained type I error control near the nominal level of 0.05. In contrast, the ad hoc same-ancestry strategies showed substantial type I error inflation, with stronger inflation at higher target cell-type proportions and causal-SNP proportions (Supplementary Fig. 11). These results show that trans-ancestry integration requires explicit modeling of ancestry-specific LD and trans-ancestry co-heritability, rather than direct reuse of same-ancestry bulk-plus-single-cell models.

To assess performance when causal variants are not fully shared across ancestries, we partitioned the same 2,000-SNP cis window into a target-null auxiliary-specific region and a shared-effect region. In the auxiliary-specific region, where no target-population causal effect was present, traceC and traceCB maintained well-calibrated target-population type I error, whereas RE2-based strategies were more prone to inflation, particularly when auxiliary or bulk-tissue signals were stronger (Supplementary Fig. 5, Supplementary Fig. 6). These results indicate that covariance-based borrowing in traceC/traceCB does not spuriously convert auxiliary-specific SNP-level signals into target-population discoveries.

Next, we evaluated the statistical power under a wide spectrum of genetic architectures. We set 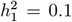 to generate eQTL effects in EAS and varied 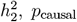 and 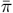 in the same way as we did in the null simulation. Besides, we considered a range of *ρ* ∈ {0.01, 0.5, 0.9} to produce varying levels of genetic correlation and varied *N*_*t*_ ∈ {1000, 5000} and *N*_1_ ∈ {50, 100} to investigate the impact of sample size. For easier demonstration, we varied a subset of the above parameters at a time while fixing the others at default values. Unless otherwise specified, we used the default parameters: 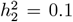, *ρ* = 0.7, *N*_1_ = 100, *N*_2_ = 400, *N*_*t*_ = 5000, and *p*_causal_ = 0.005. Because RE2-based methods showed severe type I error inflation, we omitted them from the main-text power panels to preserve visual clarity; the full comparison is reported in Supplementary Fig. 2. As shown in Fig. 2 b-d, both traceC and traceCB outperformed the original summary statistics across all settings, demonstrating the substantial benefits of leveraging auxiliary information. They achieved greater power gains when integrating a larger scRNA-seq data from EUR and when the trans-ancestry genetic correlation was higher (Fig. 2 b and d). Besides, traceCB consistently achieved higher power than traceC, particularly when the target cell-type proportion in the bulk tissue was non-negligible. This performance gain is attributable to traceCB’s ability to integrate information from large-sample bulk tissue eQTL studies. The power gained by traceCB became more pronounced in scenarios with higher target cell-type proportions or larger tissue sample sizes, where bulk eQTLs contain more relevant signal for the target cell type. For example, when integrating an EUR scRNA-seq with a sample size of *N*_2_ = 200 and a bulk tissue with a sample size of *N*_*t*_ = 5000 with a cell-type proportion of 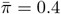, traceCB could achieve a power of 0.26 (Fig. 2 b). To achieve a similar performance, traceC would require a two-fold larger scRNA-seq sample size (*N*_2_ = 400). These results highlight traceCB’s effectiveness in leveraging the shared genetic signals from both scRNA-seq and bulk tissue data, thereby boosting the power of trans-ancestry ct-eQTL mapping.

We further decomposed the oracle power gain into the improvement from the original target-population statistic to traceC and the additional improvement from traceC to traceCB under true covariance parameters. The traceC gain increased with trans-ancestry genetic correlation, whereas the incremental traceCB gain increased with the mean target cell-type proportion and auxiliary bulk eQTL sample size (Supplementary Fig. 3). This pattern is consistent with traceCB drawing complementary information from shared cell-type-specific effects and bulk-tissue eQTL signals.

In summary, our simulations demonstrate that traceC effectively harnesses trans-ancestry information from the scRNA-seq data and traceCB achieves superior statistical power by additionally leveraging bulk tissue eQTLs. Although the RE2-based methods showed a power trend similar to traceCB (Supplementary Fig. 2), their practical utility is undermined by severe type I error inflation, especially under conditions with strong auxiliary signals. These findings suggest that traceCB is a more robust and reliable method for trans-ancestry ct-eQTL mapping compared to conventional meta-analysis approaches like RE2.

### traceCB identifies novel immune ct-eQTLs in East Asian populations

We performed trans-ancestry ct-eQTL mapping using BBJ cohort as the target population (sample size: 103, Supplementary Table 2), with a focus on immune cell types, including monocytes, CD4^+^ T cells, CD8^+^ T cells, NK cells and B cells [21, 29]. Our analysis incorporated 10 European ancestry single-cell eQTL studies from eQTL Catalogue with matched cell types as auxiliary ct-eQTL data sources (sample sizes ranging from 167 to 420, Supplementary Table 3) [12], complemented by eQTLGen blood eQTLs (sample size: 31,684, Supplementary Table 4) [6] serving as auxiliary tissue-level data. We analyzed the 10 auxiliary ct-eQTL studies separately: each study was first used by traceC to improve the baseline BBJ ct-eQTL map and was then combined with eQTLGen bulk eQTLs in traceCB. We observed that the significant cell-type-level genetic correlation of *cis*-eQTLs had a median value of 0.68–0.87 across all cell types (Supplementary Fig. 41), and was generally above 0.5, indicating the shared genetic basis of ct-eQTLs across populations. Leveraging this observation, we applied our methods to integrate scRNA-seq and bulk tissue samples to enhance ct-eQTL mapping in EAS, restricting our analysis to genes exhibiting significant trans-ancestry co-heritability (see Methods for details). Across the 10 BBJ auxiliary-study analyses, 2,054–4,184 genes passed the trans-ancestry co-heritability filter, with a mean of 2,769.8 genes per analysis (Supplementary Table 7).

We evaluated the effective sample size for the outputs of our methods, which represents the sample size that would be needed in the original study to attain an equivalent mean *χ*^2^ statistic obtained with traceCB (see Methods for details). As illustrated in Fig. 3a, the distribution of effective sample size demonstrates substantial improvements for both traceC (orange box) and traceCB (blue box) relative to the original BBJ study size (red dashed line). On average, traceC achieved a 1.82-fold (from 103 to 187.84, Supplementary Table 8) increase in effective sample size. traceCB further enhanced this gain to 2.32-fold (from 103 to 239.24, Supplementary Table 8) by incorporating eQTLGen study, highlighting the information gain from bulk tissue. The magnitude of effective sample size improvement exhibited a positive scaling relationship (*R*^2^ = 0.8377) with the sample size of auxiliary ct-eQTL studies, with the exception of the two BLUEPRINT studies that employed different sequencing methodologies (Fig. 3b, Supplementary Table 3, 8). This pattern indicates that our methods can leverage the power from trans-ancestry ct-eQTLs with larger sample sizes by effectively utilizing the shared genetic basis of eQTLs at the cell type level. Furthermore, we showed that the effective sample size of traceCB’s enhancement had a strong positive correlation (*R*^2^ = 0.8657) with the proportion of the target cell type within the bulk tissue (Fig. 3c, Supplementary Table 5, 8). Using the Fairfax_2014 (420) dataset as an example (Fig. 3d), we also investigated the impact of genetic correlation on effective sample size. As anticipated, genes with higher absolute genetic correlations showed progressively greater improvements in effective sample size for both traceC and traceCB (Fig. 3d, upper left panel). Notably, traceCB consistently outperformed traceC across all correlation strata, underscoring the additional value of integrating bulk tissue data. Gene-level effective sample size scatter plots (Fig. 3d, lower panel) further validate our approach, demonstrating substantial improvements exclusively for genes with significant trans-ancestry correlation (red points), whereas uncorrelated genes (blue points) showed no such enhancement. Similar patterns were observed in analyses using other auxiliary ct-eQTL studies (Supplementary Fig. 44–52). These findings align with our simulation results and theoretical expectations, suggesting that the power gain is attributed to shared genetic basis.

**Figure 3:**
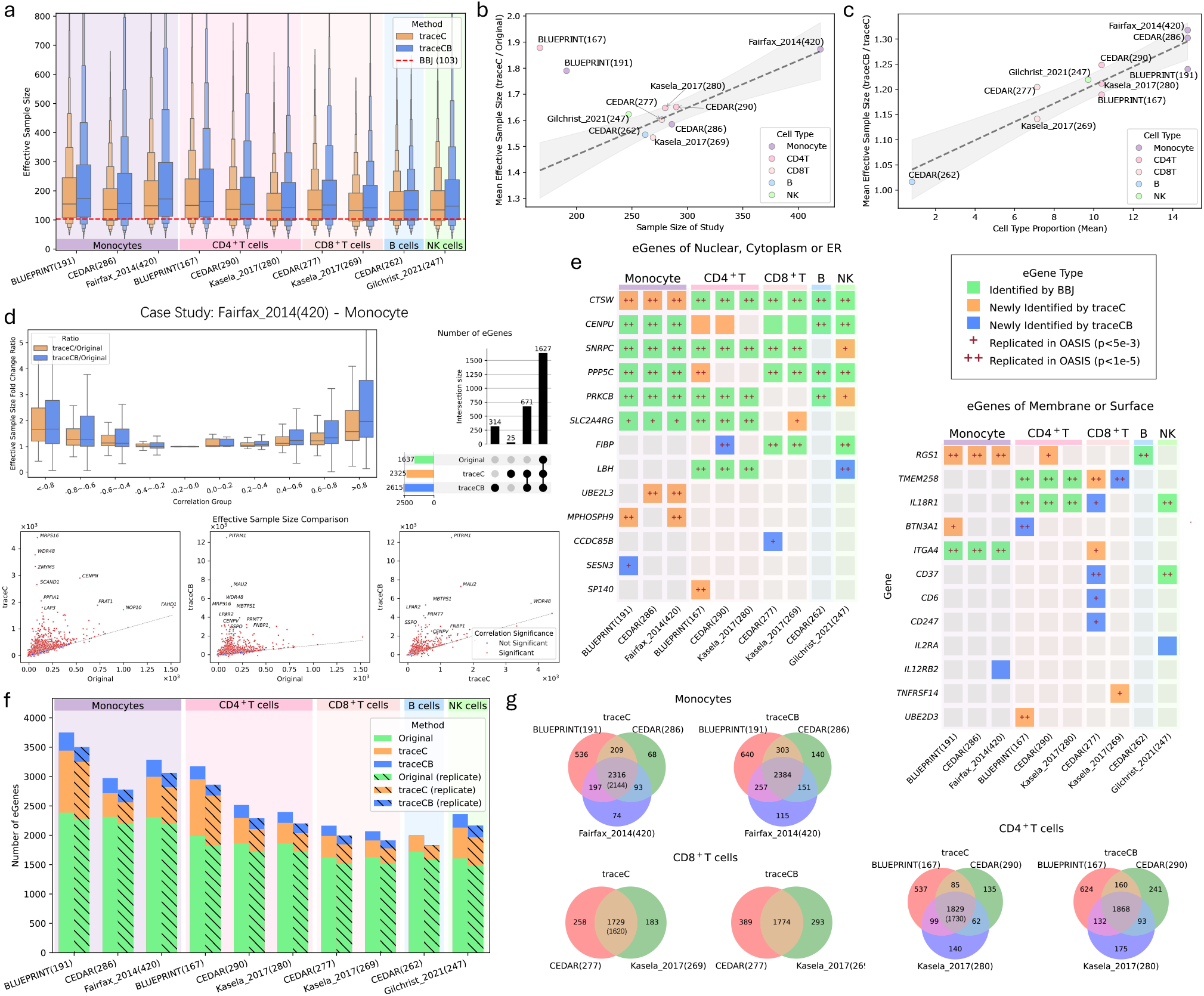
Effective sample size enhancement and eGene identification in prioritizing BBJ immune cell types using traceCB. 10 EUR single-cell eQTL studies from eQTL Catalogue served as auxiliary ct-eQTL data sources, while eQTLGen blood eQTLs provided auxiliary tissue-level information. **a.** Distribution of effective sample size for ct-eQTLs prioritized by traceC (orange) and traceCB (blue) relative to the original BBJ eQTL study sample size (red dashed line, sample size = 103). **b**,**c**. Relationship between mean effective sample size incremental fold and auxiliary data characteristics. Regression lines with 95% confidence intervals are shown in gray: **b**. traceC performance versus auxiliary ct-eQTL in terms of mean effective sample size per gene (two BLUEPRINT studies excluded as outliers in regression line due to different sequencing method); **c**. Additional effective sample size incremental fold of traceCB compared to traceC versus the target cell type’s proportion in bulk tissue. **d**. Case study of BBJ monocyte eQTL prioritization using the Fairfax_2014 (420) dataset. Upper left: Effective sample size incremental fold comparison across genetic correlation quartiles; Upper right: Number of eGenes identified of each method. Lower: Gene-wise effective sample size scatter plots distinguishing genes with (red) and without (blue) significant trans-ancestry correlation. **e**. Novel cell-type-specific eGenes identified in two functional categories: proteins localized to the nucleus, cytoplasm, or ER (left), and cell surface or membrane-associated proteins (right). Genes replicated in the OASIS dataset are marked with “+” (p < 5e-3) or “++” (p < 1e-5). **f**. Number of eGenes identified. Shaded bars indicate the proportion replicated in Wang et al.’s independent East Asian bulk eQTL study. **g**. Consistency of eGene discovery for monocytes, CD4^+^ T cells and CD8^+^ T cells across different auxiliary studies. Numbers in parentheses are the number of eGenes identified by original BBJ eQTL study.

With the improved effective sample size, our methods successfully identified more eGenes across all the cell types, defined as the genes harboring at least one significant eSNP (*p*-value < 1 × 10^−5^). On average, traceC identified 28% (from 1,929 to 2,463) more eGenes than the original BBJ study, while traceCB achieved a more substantial 38% (from 1,929 to 2,669) increase (Fig. 3f, Supplementary Table 9). With Fairfax_2014 (420) as the auxiliary study, traceC and traceCB identified 2,996 and 3,286 eGenes, respectively, representing 29.8% and 42.4% increases over the original BBJ count of 2,308 eGenes (Supplementary Table 9). The number of identified eGenes is a thresholded discovery outcome and is influenced by factors beyond the mean abundance of the target cell type in bulk tissue, including auxiliary-study sample size and quality, baseline BBJ power, the number of tested genes and SNPs, and gene-level cis-heritability and trans-ancestry co-heritability. Thus, the absolute traceCB eGene count is not expected to increase monotonically with cell-type proportion alone. To separate these contributions, we first examined the traceC/original eGene growth ratio and found that the trans-ancestry ct-eQTL gain was related to the sample size of the matched European ct-eQTL study (Supplementary Fig. 43). We then isolated the bulk-eQTL contribution by comparing traceCB with traceC. This normalized comparison showed that the additional traceCB discovery gain generally increased with the mean target cell-type proportion in whole blood (Supplementary Fig. 42), consistent with the simulation and effective-sample-size analyses. We examined the consistency of eGenes identified for monocytes, CD8^+^ T cells, and CD4^+^ T cells using different European auxiliary studies, focusing on the overlapping subset of genes across all studies. The results showed consistency, with the majority of newly identified eGenes being common across analyses (Fig. 3g). For example, in monocytes, traceC and traceCB consistently identified 2,316 and 2,384 common eGenes, respectively, representing a marked improvement over the 2,144 found in the original BBJ eQTL study. Similar results were observed for CD8^+^ T cells and CD4^+^ T cells. To rigorously validate these novel discoveries, we performed a replication analysis using an independent, large-scale bulk eQTL study of whole blood samples (*N* = 1, 019) from Japanese individuals by Wang et al. [23]. Replication was assessed at the gene level. In each discovery analysis, an eGene identified by the original BBJ analysis, traceC, or traceCB was considered replicated if the same gene was detected as an eGene in the independent dataset at the same significance threshold. This analysis did not require the same lead SNP or SNP-level allele-direction concordance across datasets. We used a gene-level criterion because Fig. 3f evaluates eGene discovery, whereas SNP-level replication across bulk and cell-type-specific eQTL datasets can be affected by differences in LD, variant coverage, allele harmonization, lead-SNP selection and cell-type composition. Under this criterion, 92% of traceC and traceCB eGenes were supported as eGenes in the independent Japanese bulk eQTL dataset, comparable to the 93% support rate observed for original BBJ eGenes (Fig. 3f, Supplementary Table 9).

The identification of cell-type-level eGenes serves as a critical step to uncover the cellular-level regulatory mechanisms of complex diseases. To evaluate the functional significance of our novel eGenes, we focused on two key pathways: protein coding genes of nucleus, cytoplasm, or endoplasmic reticulum, and genes encoding cell surface or membrane, a group particularly relevant to autoimmune diseases [8]. As shown in Fig. 3e, traceC/traceCB identified 13 eGenes associated with nucleus, cytoplasm, or endoplasmic reticulum, 5 of which were novel. For the pathway of cell surface or membrane, traceC/traceCB identified 12 novel eGenes. We sought to validate these findings by assessing their replication in the OASIS dataset, an independent East Asian immune cell eQTL study [22] with more than 1.5 million peripheral blood mononuclear cells from 235 Japanese. Remarkably, all but three of the identified eGenes in these categories replicated at a significance level of *p* < 5 × 10^−3^; the exceptions, *CENPU, IL2RA* and *IL2RB2*, were not tested in the relevant cell types in the OASIS study. Furthermore, a substantial fraction of these findings met a more stringent replication threshold of *p* < 1 × 10^−5^. To further assess whether traceC/traceCB-prioritized eGenes were supported by single-cell eQTL effects in Chinese individuals, we performed an additional replication analysis using the Chinese Immune Multi-Omics Atlas (CIMA), a large-scale immune single-cell multi-omics study of Chinese adults [30]. We used CIMA as an independent external replication resource rather than as a new target cohort for traceCB, because CIMA is already a well-powered East Asian single-cell reference and uses a much finer immune-cell taxonomy than the five broad immune-cell categories analyzed here. Among the functional eGene categories highlighted above, many traceC/traceCB-prioritized gene-cell-type pairs were replicated in CIMA at *p* < 5 × 10^−3^, and a substantial fraction reached *p* < 1 × 10^−5^ (Supplementary Fig. 55, Supplementary Fig. 56).

Taken together, these consistent replications across independent datasets provide strong support for the biological authenticity of our discoveries.

We also investigated the influence of auxiliary bulk tissue sample size on the performance of traceCB. We substituted the eQTLGen with GTEx [31] whole blood eQTLs (sample size: 670, Supplementary Table 4) as the auxiliary tissue-level data source to assess the impact of bulk tissue sample size. The results (Supplementary Fig. 59–61, Supplementary Table 8) were consistent with our previous findings, showing that traceCB outperformed traceC across all auxiliary ct-eQTL studies. However, the magnitude of improvement was more modest due to the smaller sample size of GTEx compared to eQTLGen. Specifically, the mean effective sample size for traceCB decreased to 194.14, representing a 1.9-fold improvement over the original BBJ study (Supplementary Fig. 60, Supplementary Table 8). This translated to a 29.5% (from 1,900 to 2,462) increase in the number of identified eGenes compared to the original study, with a higher replication rate of 94% in the independent dataset (Supplementary Table 10).

### Prioritized ct-eQTLs reveal cell-type-specific regulatory mechanisms for immune-mediated diseases

The increased statistical power from our framework provides an opportunity to connect cell-type-specific regulatory signals with complex immune-related traits. We therefore performed colocalization analyses between prioritized ct-eQTLs and GWAS signals for 14 blood cell traits from the Blood Cell Consortium (BCX) [32] and three immune-related traits from BBJ, including asthma [33], rheumatoid arthritis (RA) [33], and atopic dermatitis (Atopy) [34]. For the BCX analysis, we used East Asian GWAS summary statistics to match the ancestry of the BBJ eQTL data as closely as possible, thereby reducing potential bias from ancestry differences in LD structure and allele frequencies. Colocalization was performed using the R package COLOC [35].

We classified loci as colocalized when the posterior probability for a shared causal variant was P4 > 70% [36, 37], as having independent signals when P3 > 70%, and as undetermined otherwise (Fig. 4a; see Methods). Unless otherwise specified, the counts reported below summarize trait-specific locus classifications aggregated across GWAS traits and eQTL-study analyses, rather than unique genomic loci collapsed across all traits. Thus, a locus or gene colocalized with multiple traits, or supported by multiple auxiliary ct-eQTL studies, can contribute multiple times to these aggregated counts.

**Figure 4:**
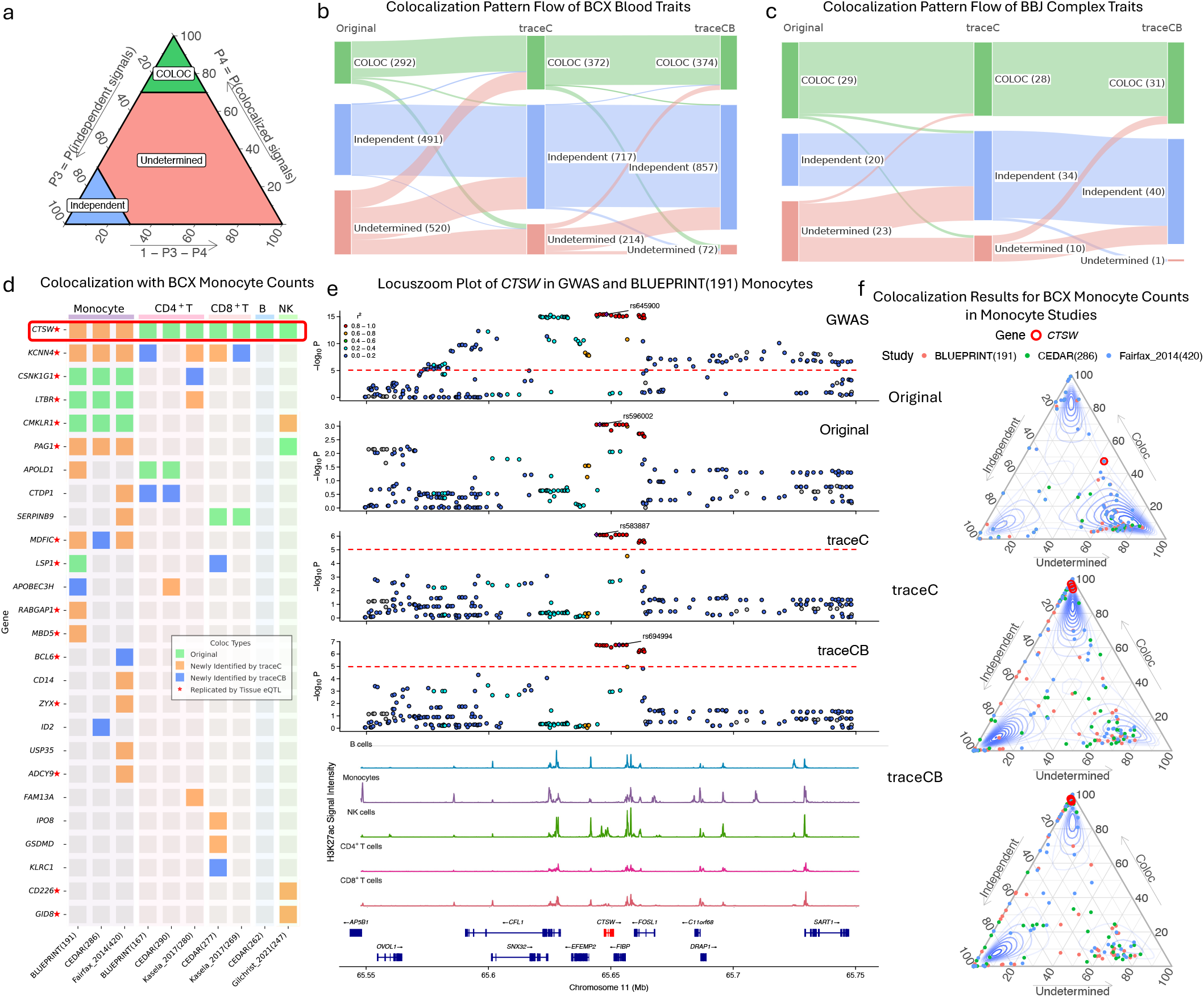
Colocalization analysis between prioritized ct-eQTLs and immune-related phenotypes. **a**. Schematic of the ternary plot interpreting colocalization posterior probabilities. The space is partitioned into regions indicating high confidence for a shared causal variant (P4 > 70%), high confidence for distinct independent signals (P3 > 70%), otherwise undetermined status. **b, c**. Comparison of colocalization patterns obtained with the original BBJ analysis, traceC, and traceCB for East Asian BCX blood cell traits (**b**) and 3 BBJ immune-related traits (**c**). Counts represent aggregated trait-specific locus classifications across the corresponding GWAS analyses, rather than unique loci collapsed across all traits; therefore, a locus associated with multiple traits can contribute once per trait-specific analysis. **d**. Heatmap of newly identified colocalized genes between monocyte ct-eQTLs and BCX monocyte count. Each entry corresponds to a gene–auxiliary-study analysis, so the same gene can appear multiple times when it is supported by multiple auxiliary ct-eQTL studies. Red asterisks next to gene names indicate genes that were also colocalized in an independent Japanese tissue-level eQTL analysis by Wang et al. **e**. LocusZoom visualization of the *CTSW* locus. Tracks display, from top to bottom: GWAS summary statistics for BCX monocyte count, original BBJ monocyte eQTLs, traceC and traceCB prioritized eQTLs (using BLUEPRINT (191) as the auxiliary study), monocyte H3K27ac ChIP-seq signals, and gene annotations. **f**. Colocalization probabilities for monocyte ct-eQTLs and BCX monocyte count across auxiliary ct-eQTL studies, showing the resolution of signal ambiguity for the *CTSW* gene (highlighted in the red circle).

Due to the limited sample size of the original BBJ ct-eQTL study, many loci remained in the undetermined region when colocalization was performed using the original ct-eQTL statistics. Prioritization by traceC and traceCB substantially reduced this ambiguity. Ternary plots showed that the posterior support shifted toward either high-confidence colocalization or high-confidence independent signals across the analyzed traits (Supplementary Fig. 14, 15). For BCX blood cell traits, traceC classified 1,089 trait-specific locus analyses as either colocalized or independent, and traceCB increased this number to 1,231, compared with 783 using the original BBJ ct-eQTL statistics (Fig. 4b, Supplementary Fig. 15). Similar trends were observed for BBJ immune-related traits, where traceC and traceCB classified 62 and 71 trait-specific locus analyses, respectively, compared with 49 from the original analysis (Fig. 4c, Supplementary Fig. 14). These results indicate that traceC and traceCB improve the resolution of downstream colocalization analyses by strengthening target-population ct-eQTL signals, thereby increasing the ability to distinguish shared regulatory signals from independent trait and eQTL associations. We further summarized study-level sharing of colocalized genes using an UpSet analysis, in which colocalized results were first collapsed to unique genes within each eQTL study before overlaps across studies were evaluated (Supplementary Fig. 16). This analysis complements the trait-specific counts in Fig. 4b–c and shows that many colocalized genes were supported across multiple auxiliary eQTL studies.

Among the analyzed traits, monocyte count provided a clear example of the improved interpretability of traceC/traceCB-prioritized ct-eQTLs. Monocyte count is an important indicator of immune activity and chronic inflammation, and identifying its regulatory determinants can provide insight into immune-mediated disease biology. Fig. 4d summarizes newly identified colocalizations between monocyte ct-eQTLs and BCX monocyte count across auxiliary ct-eQTL studies. traceC identified 36 new gene–study colocalization events, and traceCB further increased this number to 41 by incorporating bulk tissue eQTL information. Several of these newly prioritized genes were also supported in the independent East Asian whole-blood eQTL study by Wang et al. [23], providing gene-level external support for the regulatory signals. Notably, the newly colocalized genes included both signals shared across multiple immune cell types and signals restricted to a subset of cell types. For example, *CTSW*, a known cytotoxic gene with high transcriptional activity in monocytes [38], was identified as an eGene in all the immune cell types except for monocytes when using the original BBJ eQTL study (Fig. 4d, Supplementary Fig. 17–25). However, the H3K27ac ChIP-seq tracks reveal distinct regulatory peaks at the *CTSW* locus across all 5 cell types, indicating accessible chromatin and active regulation (Fig. 4e). Our traceC and traceCB successfully identified this functionally relevant eQTL in monocytes, leading to a colocalization with monocyte count. As shown in Fig. 4e, Supplementary Fig. 17 and 18, while the original analysis could not identify a significant association, both traceC and traceCB revealed highly significant eQTL signals that closely align with both the GWAS signal of monocyte count and the H3K27ac enrichment peaks at the *CTSW* locus. Specifically, when using BLUEPRINT (191) as the auxiliary scRNA-seq study, traceC elevated the leading SNP’s significance level to *p* ≈ 1 × 10^−6^, and traceCB further amplified it to *p* ≈ 1 × 10^−7^. These enhancements, which were consistently observed across different auxiliary studies, dramatically increased the colocalization probability P4 of *CTSW* from less than 50% to nearly 100% (Fig. 4f).

As a second example, the analysis of *KCNN4* illustrates the benefit of incorporating tissue-level data. *KCNN4* is a calcium-activated potassium channel implicated in immune cell function [39, 40], and recent study has highlighted its role in modulating immune responses during COVID-19 infection [41]. Although this gene’s eQTL was replicated in the East Asian bulk study by Wang et al. [23], the original BBJ analysis showed no colocalization evidences. However, epigenetic data (H3K27ac) indicates potential regulatory activity across all 5 immune cell types (Supplementary Fig. 33). By leveraging trans-ancestry scRNA-seq data with traceC, we detected 5 new colocalizations with BCX monocyte count GWAS signals, while traceCB further increased this to 7 new colocalizations by integrating bulk tissue information (Fig. 4d, Supplementary Fig. 26–32). The absence of results from three auxiliary studies is due to data unavailability for *KCNN4* in those datasets.

Finally, the colocalization of *MDFIC* highlights our framework’s ability in detecting cell-type-specific signals. *MDFIC* is associated with lymphatic function [42]. In our analysis, we observed strong eQTL signals exclusively in monocytes when prioritized by traceC and traceCB (Supplementary Fig. 35–40), leading to significant colocalization with monocyte count GWAS signals (Fig. 4d). Consistent with this, we identified strong H3K27ac, GWAS, and eQTL signals in the region from chr7: 114.65Mb to 114.75Mb (GRCh37) specifically in monocytes (Supplementary Fig. 34–40).

In summary, these colocalization analyses demonstrate that traceC and traceCB strengthen target-population ct-eQTL signals in a way that improves downstream trait-linking analyses. Rather than providing definitive causal fine-mapping, these results prioritize regulatory loci with increased posterior support for either shared or independent signals, resolve a subset of previously ambiguous loci, and highlight functionally relevant gene–trait associations that were not detected using the original BBJ ct-eQTL statistics alone.

### Application of traceCB to identify ct-eQTLs in African population

To rigorously test the broad applicability and robustness of our framework, we extended our analysis to an African ancestry cohort (sample size: 80, Supplementary Table 4) [43]. This population presents a more challenging test case due to its smaller sample size than the previous BBJ eQTL study (sample size: 103, Supplementary Table 2) and greater genetic divergence from the European auxiliary populations used as a reference. Aligned with our previous analysis, we utilized 10 European single-cell eQTL studies with matched cell types as auxiliary ct-eQTL data sources (sample sizes ranging from 167 to 420, Supplementary Table 3) [12], along with eQTLGen blood eQTLs (sample size: 31,684, Supplementary Table 4) [6] as auxiliary tissue-level data. In the African ancestry analysis, 156–452 genes passed the trans-ancestry co-heritability filter across the 10 auxiliary studies, with a mean of 276.6 genes per analysis (Supplementary Table 11).

Our framework demonstrated robust performance, mirroring the results from the East Asian cohort. We observed a substantial increase in the effective sample size for both traceC and traceCB relative to the original African study (Fig. 5a). On average, traceC increased the effective sample size by 1.50-fold (from 80 to 119.8) and traceCB by 1.70-fold (from 80 to 135.9, Supplementary Table 12, Supplementary Fig. 62). This relative gain, which was greater than that observed in the BBJ cohort, was likely due to the lack of power of the smaller African study. Therefore, integrating trans-ancestry eQTLs yielded a greater improvement in power. Notably, the magnitude of this enhancement scaled with the sample size of the auxiliary data, the genetic correlation of each gene and proportion of target cell types (Supplementary Fig. 62– 72). For instance, integration of the larger Fairfax_2014 (420) dataset with traceC yielded greater improvements than the smaller CEDAR (286) dataset (Fig. 5, Supplementary Table 12, Supplementary Fig. 62). The performance gains in the African cohort were remarkably consistent than in the BBJ cohort. Nearly all genes with significant positive trans-ancestry genetic correlation exhibited an improvement (Fig. 5b, Supplementary Fig. 66–72). This consistent pattern of leveraging shared genetic architecture across diverse ancestries and datasets further underscores the robustness of our approach. As expected, both traceC and traceCB identified a considerable number of novel eGenes missed by the single-ancestry analysis (Supplementary Fig. 64). The magnitude of this improvement also scaled with the power of the auxiliary data, with the larger Fairfax_2014 (420) dataset yielding more discoveries. Overall, traceC identified 57 (from 502 to 559) more eGenes than the original African eQTL study, while traceCB achieved a 67 (from 502 to 569) increase (Supplementary Table 12). These findings confirm that our framework can robustly enhance ct-eQTL discovery even in small, genetically distinct populations, yielding biologically meaningful results.

**Figure 5:**
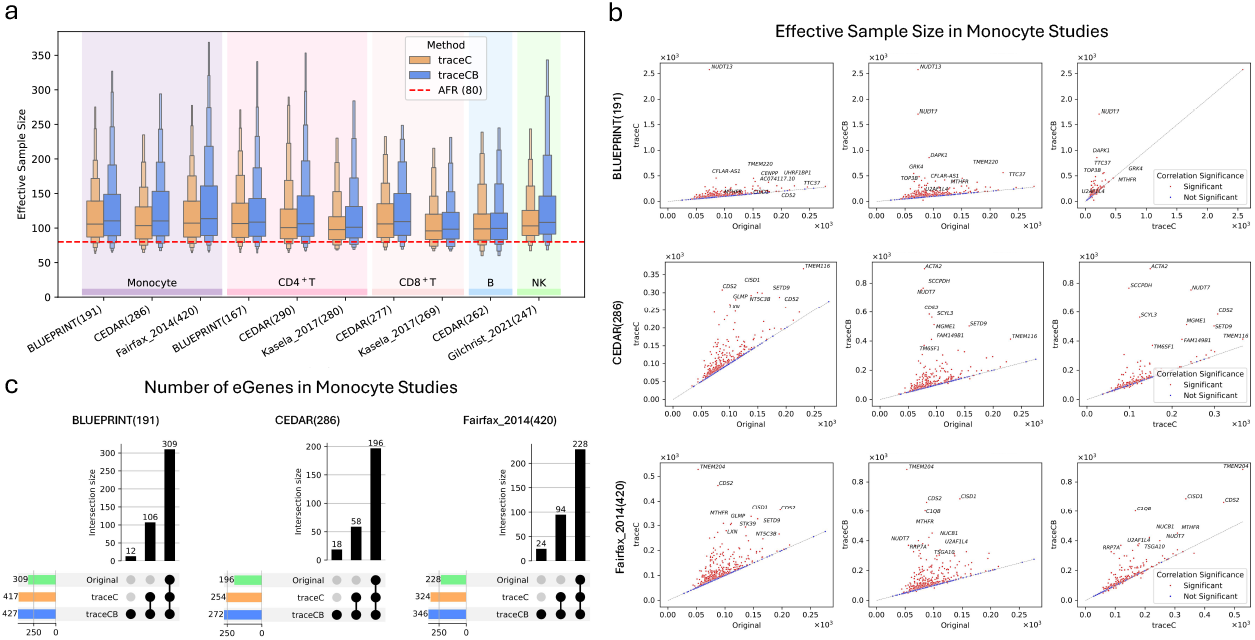
Application of traceC and traceCB to enhance ct-eQTL mapping in an African ancestry cohort. The analysis used European auxiliary data from eQTL Catalogue and eQTLGen blood eQTLs to prioritize ct-eQTLs in an African cohort. **a.** Distribution of effective sample size for ct-eQTLs prioritized by traceC and traceCB, compared to the original study’s sample size (red dashed line: sample size = 80). **b**,**c**. Case studies of eGene identification and effective sample size enhancement in monocytes, using three different European auxiliary datasets: BLUEPRINT (191), CEDAR (286) and Fairfax_2014 (420). **b**. Comparison of effective sample size between the original analysis with traceC/traceCB. Red points denote genes with significant trans-ancestry genetic correlation, which are targeted for enhancement, while blue points represent non-targeted genes. **c**. Upset plots illustrate the overlap and novel discovery of eGenes by traceC and traceCB compared to the original analysis.

## Discussion

In this paper, we present traceCB, a statistical framework that addresses the challenge of underpowered ct-eQTL mapping in underrepresented ancestries. By integrating summary statistics from single-cell and bulk-tissue eQTL studies across diverse populations, traceCB leverages the statistical power of large European ancestry cohorts to enhance discovery in target populations such as East Asian and African. In contrast to conventional meta-analysis approaches (for example, RE2) that often assume homogeneous effects across studies, traceCB explicitly models both cell-type-specific heritability and trans-ancestry co-heritability. This design allows the framework to adaptively borrow information, increasing power for genes with shared genetic architecture while avoiding spurious associations at genetically distinct loci. Our analyses show that traceCB increases effective sample size, improves the detection of novel and replicable eGenes, and strengthens the functional interpretation of GWAS signals through improved colocalization with related GWAS loci.

Our traceCB framework needs further investigation in the following directions. First, our analyses have been mainly focused on the blood cell types. Although the single-cell eQTL data have been collected in other human tissues, such as brain [44, 45] and liver [46] for the European ancestry, they remain unavailable in the under-represented populations. This data unavailability restricted the application of traceCB to a more comprehensive analyses of disease-related cell types. With the accumulating single-cell eQTL data resources, we expect that traceCB will be important for translating these resources into cellular-level biological insight and more equitable applications in human genetics.

Second, the performance of our method depends on the availability and quality of auxiliary data. The eQTLGen study includes a fraction of non-European samples; for example, African ancestry samples account for 8.8% of the overall samples [6]. Despite this, we found that traceCB is robust to this contamination in our applications, as evidenced by the high replication rate of traceCB’s output. Besides, its effectiveness naturally scales with the sample size, technical quality and methodological consistency of the contributing studies. Continued generation of large, well-characterized single-cell and bulk eQTL resources from ancestrally homogeneous and admixed cohorts will be important for realizing the full potential of this type of integrative approach.

Third, traceCB requires accurate estimation of gene-level heritability and co-heritability parameters, which can be difficult in settings with limited sample sizes and few SNPs at individual loci. We also conducted additional simulations where the true heritabilities and genetic correlation parameters were known (Supplementary Fig. 2). With the true parameters, traceC achieved a statistical power nearly identical to RE2 while maintaining strict type I error control. Notably, the power advantage of traceCB over traceC was even more pronounced when the true parameters were used, underscoring the potential for further performance improvements through more accurate parameter estimation techniques in future work. Developing more robust and efficient estimators by pooling information across functionally relevant genes [50] is therefore an important direction for future work.

Fourth, deconvolution of bulk-tissue data in traceCB relies on mean cell-type proportions inferred by external tools, which are treated as fixed inputs. Although the sensitivity analysis suggests that traceCB remains conservative for false-positive control under misspecified mean cell-type proportions, the current implementation does not propagate deconvolution uncertainty into downstream inference. Errors in cell-type proportion estimates may still affect power and the magnitude of bulk-derived information borrowing, especially for rare cell types or tissues with complex cellular composition. A future extension could incorporate uncertainty-aware deconvolution estimates or jointly model the distribution of cell-type proportions within the traceCB variance framework. Moreover, our current formulation models cell types and cell states as discrete categories, whereas more refined representations, such as continuous or hierarchical cell-state models, may capture regulatory heterogeneity more accurately [51–56].

Finally, the current implementation focuses on integrating eQTL data from two populations. Extending the framework to accommodate additional populations and multiple molecular layers (for example, splicing QTLs, protein QTLs, methylation QTLs, and chromatin accessibility QTLs) would increase its scope in the context of multi-omics and global genomics.

## Methods

### The traceCB model

To begin with, we introduce the traceCB model with the individual-level scRNA-seq and bulk RNA-seq data. For an interested cell type’s gene expression in a target underrepresented population, we consider its scRNA-seq eQTL data {**y**_1_, **X**_1_}, where 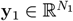 is the normalized gene expression vector and 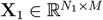 is the standardized genotype matrix, *M* is the number of *cis*-SNPs around the target gene, *N*_1_ is the sample size of target scRNA-seq data. Typically, *N*_1_ is quite small in the non-EUR populations, e.g., *N*_1_ ≈ 100. We assume that the covariates such as genotype principal components and PEER factors have been properly adjusted, with details given in our previous studies [25, 60]. To relate the gene expression to genotype, we consider the following linear model:

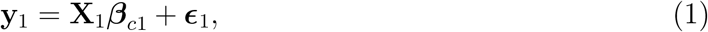

where ***β***_*c*1_ = [*β*_*c*11_, …, *β*_*c*1*M*_]^⊤^ ∈ ℝ^*M*^ collects the effect sizes of *cis*-SNPs for the interested cell type, 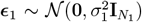 is the independent noise term. The small sample size of scRNA-seq data limits the power to identify the *cis*-SNPs associated with the target gene.

In the main traceCB formulation, we assume that the auxiliary scRNA-seq ct-eQTL data and the auxiliary bulk tissue eQTL data are collected from the same auxiliary ancestry, typically European ancestry. This assumption matches the primary applications in this study, where the target ct-eQTL datasets are from East Asian or African ancestry cohorts and the largest available auxiliary scRNA-seq and bulk eQTL resources are predominantly European. Extensions to target-ancestry or multi-ancestry bulk eQTL resources are discussed in Supplementary Note 3.4.

Suppose that we have additionally collected the scRNA-seq and bulk tissue eQTL data of an auxiliary population, typically from the European ancestry, denoted as {**y**_2_, **X**_2_} and {**y**_*t*_, **X**_*t*_}, where 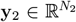 is the gene expression vector of the interested cell type, 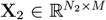 is the genotype matrix of *cis*-SNPs of the individuals from scRNA-seq samples, 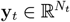 collects the gene expression levels in the bulk tissue, 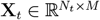 is the genotype matrix of bulk samples, and *N*_2_ and *N*_*t*_ are the sample sizes of the two auxiliary studies with *N*_2_ ∈ [200, 500] and *N*_*t*_ ∈ [500, 30000]. We also assume that the gene expression vectors and columns of genotype matrices have been normalized, and that relevant covariates have been properly adjusted. The relationship between gene expression levels and genotypes can be characterized with the following linear models:

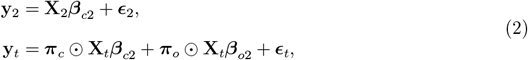

where ***β***_*c*2_ ∈ ℝ^*M*^ and ***β***_*o*2_ ∈ ℝ^*M*^ are the *cis*-SNP effect sizes in the target cell type and a surrogate for the other cell types in tissue, respectively, 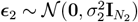, and 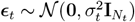 are independent error terms, 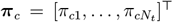 and 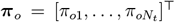 represent the proportions of the cell types within tissue samples, and ⊙ denotes the element-wise product. The second equation in Equation (2) is granted by the fact that the bulk gene expression levels can be decomposed into a weighted sum of genetically controlled expressions in cell types (**X**_*t*_***β***_*c*2_ and **X**_*t*_***β***_*o*2_) with the weights being cell type proportions *π*_*c*_ and *π*_*o*_.

Here, ***β***_*o*2_ should be interpreted as an aggregate nuisance effect representing the combined genetic contribution from non-target cell types in the bulk tissue, rather than the effect of a single explicitly observed cell type. In the two-component working model in Equation 2, ***π***_*c*_ denotes the target cell-type proportion and ***π***_*o*_ = **1** − ***π***_*c*_ denotes the aggregate proportion of all non-target cell types. More generally, if the bulk tissue contains *K* cell types, the tissue expression can be written as

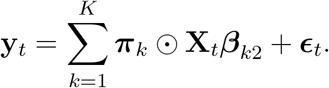

For inference on a target cell type *c*, the term ***π***_*c*_ ⊙ **X**_*t*_***β***_*c*2_ is the component that is directly informative for the target cell-type-specific eQTL effect. The remaining contribution, ∑_*k*≠*c*_ ***π***_*k*_ ⊙ **X**_*t*_***β***_*k*2_, is summarized in traceCB by the aggregate nuisance component ***π***_*o*_ ⊙ **X**_*t*_***β***_*o*2_. This aggregation does not assume that non-target cell types have no genetic effects; rather, it reflects the fact that, with only tissue-level summary statistics, the individual non-target cell-type effects {***β***_*k*2_ : *k≠ c}* and their proportions are not separately identifiable without additional assumptions or external cell-type-specific reference data. Therefore, traceCB focuses on estimating the target cell-type effect while treating the combined non-target contribution as a background component.

In the summary-level GMM formulation, the bulk-tissue association statistic is informative for the target ct-eQTL only through the target-cell component, scaled by the mean target cell-type proportion and linked to the target-population effect through the trans-ancestry covariance between ***β***_*c*1_ and ***β***_*c*2_. The aggregate non-target component contributes to the variability of the tissue-level statistic but is not separately identified or used as direct evidence for the target cell-type effect. This modeling choice keeps the summary-level estimator identifiable and scalable, while acknowledging that unresolved non-target cell-type effects may reduce the amount of useful information that can be borrowed from bulk tissue.

Due to the unbalanced sample composition and technical constraints, typically, we have *N*_1_ < *N*_2_ ≪ *N*_*t*_. Considering the substantial sharing of genetic basis across populations, we aim to leverage the information of the larger scRNA-seq and bulk tissue eQTL studies from the auxiliary cohorts to enhance the ct-eQTL mapping in the target population.

To characterize the shared genetic basis of cell-type-level eQTL effects across populations, we introduce the following probabilistic model for the effect sizes ***β***_*c*1_ and ***β***_*c*2_:

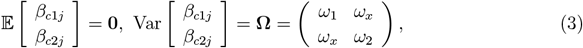

where **Ω** captures the covariance of cell type-specific *cis*-effects between the two populations, *ω*_1_ and *ω*_2_ are the per-SNP *cis*-heritability for target and auxiliary population, respectively, and *ω*_*x*_ is the covariance (per SNP *cis*-co-heritability) between the two populations. Notably, the per-SNP *cis*-eritability *ω* is related to the total *cis*-heritability *h*^2^ by the number of SNPs *M* in the *cis*-window, such that *ω* ≈ *h*^2^*/M*. The probabilistic model (3) exhibits four distinct features. First, its flexible design allows the effect sizes of *cis*-SNPs to vary across populations. This characteristic distinguishes traceCB from traditional meta-analysis methods that rely on the assumption of homogeneous effects. Second, it characterizes the shared genetic information in specific cell types via the covariance *ω*_*x*_, which quantifies the average covariance of *cis*-SNPs within a local region. A non-zero *ω*_*x*_ indicates correlated effect sizes between the two populations, which can be either positive or negative. When ***β***_*c*1_ and ***β***_*c*2_ diverge significantly, *ω*_*x*_ approaches zero. By empirically estimating *ω*_*x*_ from the data (details in Supplementary Section 3.5), traceCB can adaptively borrow cell type-specific information from auxiliary scRNA-seq data while mitigating the risk of introducing spurious signals. Third, the framework enables the integration of large-scale auxiliary bulk samples. When the interested cell type exhibits both significant non-zero genetic covariance (*ω*_*x*_*≠* 0) and constitutes a substantial proportion (i.e., ***π***_*c*_ > 0) in the auxiliary bulk tissue, well-powered tissue-level eQTL studies can be leveraged to enhance ct-eQTL mapping (details in Supplementary Section 3.6). Fourth, the model relies solely on moment conditions, making no assumptions regarding the distribution of eQTL effects. With these moment conditions, traceCB is capable of scaling effectively to accommodate arbitrarily complex genetic architectures.

### The traceCB model with summary-level eQTL data

In the real applications, individual-level eQTL data may not be easily accessible. Instead, we usually have access to the ct-eQTL summary statistics of two populations, 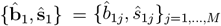 and 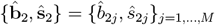, and the tissue-level eQTL summary statistics 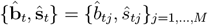, which can be obtained from marginal regressions:

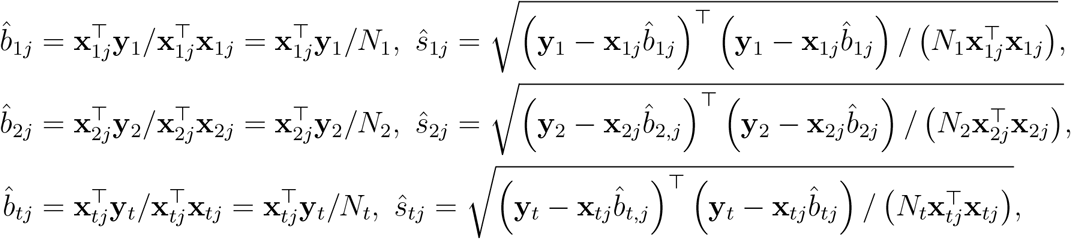

where **x**_1*j*_, **x**_2*j*_ and **x**_*tj*_ are the *j*-th columns of the genotype matrices **X**_1_, **X**_2_ and **X**_*t*_, respectively. In the following, we aim to derive a traceCB estimator that improves 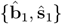 by using the information from 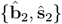 and 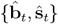.

To work with the summary statistics defined above, we consider the random design of genotype matrices that assumes the rows of **X**_1_, **X**_2_, and **X**_*t*_ are drawn from independent and identical distributions from the two populations, receptively. Then, we can denote the correlation between SNPs *j* and *k* in the two populations as 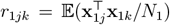 and 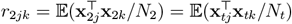, respectively. As such, we can define the underlying true marginal effect sizes at cell type level as:

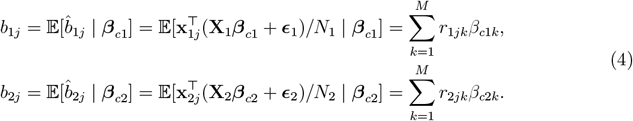

Clearly, the true marginal effect sizes at the cell-type level, *b*_1*j*_ and *b*_2*j*_, are a weighted sum of the true effect sizes of local SNPs in LD with SNP *j*, where the weights are the LD correlations in the corresponding population. Similarly, we can obtain the true marginal effects at the bulk tissue level:

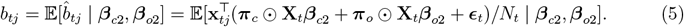

A close observation on the above expression reveals that ***π***_*c*_ and ***π***_*o*_ varies across samples, making it difficult to relate the tissue level marginal effects *b*_*tj*_ to the cell type level effects. To overcome this difficulty, we prove that with reasonably large sample size of the bulk data, the individual-level cell type composition vectors ***π***_*c*_ and ***π***_*o*_ can be approximated with the mean proportions across bulk samples, denoted as 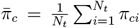 and 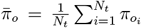, respectively [16]. The detailed proof is given in Supplementary Note 3.1.

Therefore, Equation (5) can be updated as:

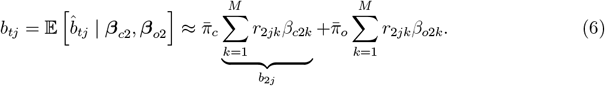

As we can see, this approximation decomposes *b*_*tj*_ into the sum of the true marginal effects in the target and the other cell types weighted by the mean cell type proportions. As such, the relationship between *b*_*tj*_ and *b*_2*j*_ becomes clear. We have conducted extensive simulation studies to show that this approximation is highly accurate when *N*_*t*_ > 500, which is satisfied in most of the bulk eQTL data [16].

With the above specifications, the following summary-level model can be established:

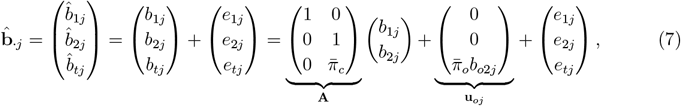

where **A** represents the transformation matrix linking the observed marginal effects to the true cell type level effects, **u**_*oj*_ captures the aggregated non-target cell-type contribution to the bulk marginal effect, with 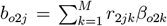. This nuisance component is not treated as a target-cell mean component and contributes to the estimator only through the covariance term **Σ**_*oj*_, and *e*_1*j*_, *e*_2*j*_, and *e*_*tj*_ denote the independent estimation errors for the *j*-th *cis*-SNP in the respective populations, with variances 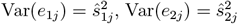, and 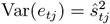.

Given the above summary-level model, we can establish the relationship between the summary data and variance component parameter **Ω**. By combing models (4) and (6) with the moment conditions (3), we have:

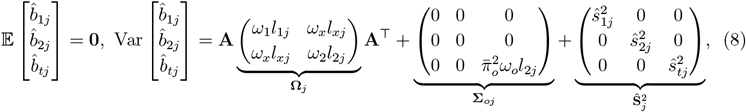

where 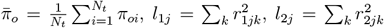 are the LD scores in two populations, respectively, and 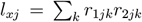 is the trans-ancestry LD score for the *j*-th *cis*-SNP. *ω*_*o*_ is the per-SNP heritability term for the aggregated non-target cell-type component (Details in Supplementary Note 3.6). This covariance decomposition uses the working assumption that the target and aggregated non-target cell-type genetic components are uncorrelated after conditioning on ancestry-specific LD; cross-cell-type co-heritability terms can be added to **Σ**_*oj*_ when such components are explicitly modeled. Under the genetic drift assumption of LDSC model, we can generalize models (1) and (2) to account for confounding bias, leading to a modified covariance of marginal effects in Equation (8):

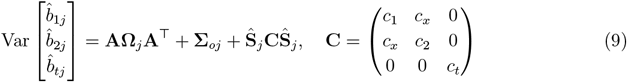

where *c*_1_, *c*_2_, *c*_*x*_, and *c*_*t*_ are inflation constants reflecting the hidden confounding bias such as sample overlap and residual population structures. The inflation constants can be set to their theoretical values or adaptively estimated from the data, depending on the presence of population structures and sample overlap [27].

### The traceCB estimator

With the summary-level model, we will introduce the traceCB estimator of the target population’s ct-eQTL effect based on GMM. Based on Equations (7) and (9), we first obtain the conditional mean and covariance of the observed marginal effects (details in Supplementary note 3.2):

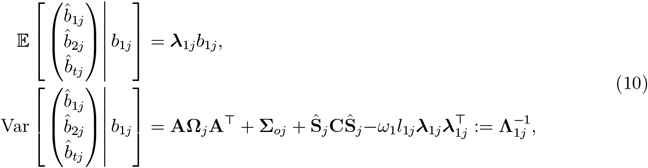

where 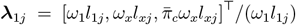 is the SNP-specific projection coefficient. This notation keeps the per-SNP variance-component parameters *ω* while explicitly incorporating the LD scores used in **Ω**_*j*_. Hence, we can define the moment equation

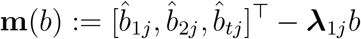

and the first-order moment condition:

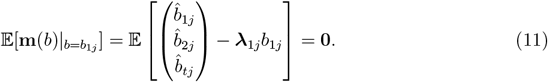

To obtain the traceCB estimator of 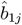, we apply GMM to solve for the moment condition (11):

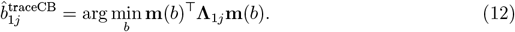

It turns out this optimization problem has closed form solution:

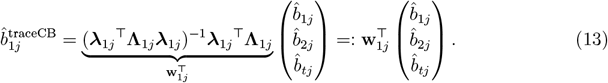

The variance of the above traceCB estimator is given as

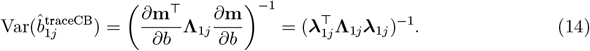

Notably, traceCB estimator given in Equation (13) is determined by three major components. First, it incorporates the genetic covariance **Ω**, which enables traceCB to borrow information from auxiliary scRNA-seq data for improving ct-eQTL mapping when *ω*_*x*_ ≠ 0. When *ω*_*x*_ = 0, traceCB estimator is equivalent to the original ct-eQTL statistics, which prevents falsely transferring unshared signals from the auxiliary population. Second, **Ω** is further weighted by the trans-ancestry LD scores *l*_1*j*_, *l*_2*j*_, and *l*_*xj*_, effectively accounting for heterogeneous LD between populations. Third, the information from bulk eQTL study is also incorporated via the mean cell type proportion 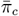 in the link matrix **A**, as defined in Equation (8). A non-zero 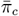 allows traceCB to utilize the information from the large-scale bulk eQTL study.

### Parameter estimation

The traceCB estimator requires estimates of the genetic covariance matrix **Ω**, the inflation constant matrix **C**, and the mean cell type proportion 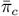. To estimate the mean cell type proportions 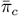, we first apply cell type deconvolution methods to the bulk RNA-seq data using CIBERSORTx [28], and then average the estimated proportions across samples to obtain the mean proportions.

We estimate **Ω** and **C** using a trans-ancestry LDSC framework [27, 61]. Note that the covariance structure from Equation (9) can be rearranged into the following equations related to Z scores:

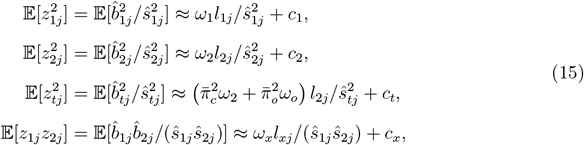

The unknown parameters can be obtained by regressing the realization of the products of 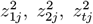, and *z*_1*j*_*z*_2*j*_ to the corresponding LD scores, *l*_1*j*_, *l*_2*j*_, and *l*_*xj*_, respectively. The slopes of these regressions provide estimates for the per-SNP heritabilities (*ω*_1_, *ω*_2_, *ω*_*t*_) and co-heritability (*ω*_*x*_), while the intercepts are the estimates of inflation parameters (*c*_1_, *c*_2_, *c*_*x*_, *c*_*t*_). Following LDSC [60, 61], we use a weighted least squares estimator to account for dependence among Z scores. See Supplementary Note 3.5 and 3.6 for more details on the regression equations and the estimation procedure.

Once we have estimated the heritabilities and inflation constants, we can plug the final estimates 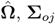 and **Ĉ** into Equations (13) and (14) to obtain traceCB estimators. To ensure the robustness of our method, we adopt a conservative strategy that only utilizes the genetic correlation that exhibits strong significance across populations. Specifically, we conducted a Wald test to assess the statistical significance of each covariance component and only retained the corresponding covariance estimate with p-values smaller than a stringent threshold (i.e., *p* < 0.10 in this study); otherwise, we set the covariance term to zero. If the estimated variance of the traceCB estimator is negative, which typically arises when 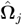 is not positive definite, we revert to reporting the results from traceC. For the bulk nuisance component, when 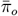 is close to zero or the LDSC slope for the tissue-level variance yields a negative residual estimate after subtracting the target-cell contribution, we truncate the corresponding nuisance variance at zero and use the reduced positive-definite covariance matrix; if the resulting GMM variance remains non-positive, we revert to the corresponding estimator that excludes the unstable bulk component.

With the estimation of 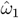 and 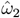, we can evaluate the effective sample size for the outputs of our methods using the standard LD score regression relationship between the mean association *χ*^2^ statistic, sample size, per-SNP heritability, and LD score. For a SNP *j* in the cis region, the marginal genetic effect can be written as *b*_*j*_ =∑_*k*_ *r*_*jk*_*β*_*k*_, where *r*_*jk*_ denotes the LD correlation between SNPs *j* and *k*. Under the local random-effects model with per-SNP cis-heritability *ω*, we have

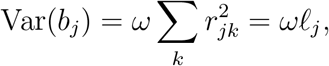

where *ℓ*_*j*_ is the LD score of SNP *j*. After standardization, the association statistic satisfies 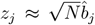, and the null sampling variance contributes 1 to the expected squared *z*-score. Therefore, following LD score regression,

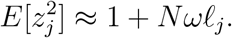

Averaging over SNPs within the cis region gives

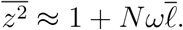

Solving this equation for *N* yields the effective sample size estimator

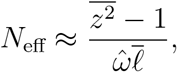

where 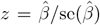 denotes the marginal association *z*-score, 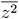 is the mean squared *z*-score across SNPs within the *cis* region, 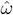 represents the estimated per-SNP *cis*-heritability, and 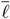 is the average LD score for SNPs within the same *cis* region [27].

In our analysis, *z* is computed from the output of the original, traceC, or traceCB statistic, and 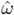 is the corresponding target-population per-SNP cis-heritability estimate. Thus, *N*_eff_ should be interpreted as the sample size required in a primary target-population ct-eQTL study to attain an equivalent mean *χ*^2^ statistic to that achieved by traceC or traceCB under the same local heritability and LD structure.

### Chromosome-wide mixture-architecture simulations

To better evaluate traceC and traceCB under more realistic gene-level architectures, we performed an additional chromosome-wide simulation using genes on chromosome 22. For each gene, cis-SNPs were defined using the gene body *±*500 kb window. Genes were assigned to one of four architecture classes: shared-effect genes, population-1-specific genes, population-2-specific genes, and fully null genes, with proportions 30%, 20%, 20%, and 30%, respectively. Shared-effect genes had causal effects in both populations with genetic correlation *r*_*g*_ = 0.7; population-1-specific genes had causal effects only in the target population; population-2-specific genes had causal effects only in the auxiliary population and tissue component; and null genes had no causal eQTL effects in either population or tissue component. We evaluated both gene-level power and gene-level type I error using the same eGene-level significance criterion as in the real-data analysis. Bars show gene-level means across genes within each architecture class, and error bars represent 95% confidence intervals across genes.

The chromosome-wide mixture simulation confirmed the behavior observed in the single-region simulations while evaluating gene-level outcomes under heterogeneous architectures. For shared-effect genes, traceC and traceCB improved target-population eQTL discovery power over the original statistic, with traceCB showing additional gain when the target cell-type proportion in bulk tissue was higher (Supplementary Fig. 10). For population-1-specific genes, where auxiliary information was not informative, traceC and traceCB remained close to the original target-population statistic, indicating that the methods did not force borrowing when trans-ancestry sharing was absent (Supplementary Fig. 9). For fully null genes, all methods maintained gene-level type I error near the nominal level (Supplementary Fig. 7). In the more challenging population-2-specific setting, where auxiliary and tissue signals were present but the target population was null, traceC and traceCB maintained calibrated target-population type I error, whereas RE2(sc+tissue) showed substantial inflation, especially as the target cell-type proportion in bulk tissue increased (Supplementary Fig. 8). These results provide a gene-level validation that traceC/traceCB gain power primarily for shared genetic architectures while avoiding spurious transfer of auxiliary-population-specific signals.

### Preparation of eQTL data

We obtained cell type-specific eQTL summary statistics for the East Asian population from the BBJ project [29], publicly available at https://humandbs.dbcls.jp/en/hum0099-v1. The dataset encompasses 5 major immune cell types, defined by the following surface markers: CD4^+^ T cells (CD3^+^CD4^+^CD8^−^CD19^−^), CD8^+^ T cells (CD3^+^CD4^−^CD8^+^CD19^−^), B cells (CD3^−^CD19^+^), NK cells (CD3^−^CD14^−^CD19^−^CD56^+^), and monocytes (CD3^−^CD14^+^CD19^−^). Only single nucleotide variants (SNVs) with an assigned dbSNP Reference SNP cluster ID (RSID) and non-palindromic alleles were retained. The processed summary statistics for each cell type included approximately 20,000 genes, with an average of 3,700 SNPs per gene (Supplementary Table 2).

For auxiliary ct-eQTL data, we curated 10 European-ancestry studies from the eQTL Catalogue [12] matched to the 5 immune cell types in BBJ: CD4^+^ T cells (BLUEPRINT [62], Kasela_2017 [63], CEDAR [64]), CD8^+^ T cells (CEDAR [64], Kasela_2017 [63]), B cells (CEDAR [64]), NK cells (Gilchrist_2021 [65]), and monocytes (BLUEPRINT [62], Fairfax_2014 [66], CEDAR [64]). After filtering for RSIDs, these datasets contained approximately 19,000 genes and 5,800 SNPs per gene for each pair (Supplementary Table 3). For the African ancestry target cohort, we generated ct-eQTL summary statistics using data from the POPCELL project [43], which comprises genotype and single-cell RNA-seq data for 80 healthy individuals. Following the official POPCELL pipeline available at https://github.com/h-e-g/popCell_SARS-CoV-2, we performed *cis*-eQTL mapping for SNPs with a minor allele frequency (MAF) ≥ 0.05 within a 1-Mb window of each gene using MatrixEQTL [67]. This yielded summary statistics for 12,114 genes per cell type, with an average of 1,279 SNPs per gene (Supplementary Table 4).

For auxiliary tissue-level data, we utilized whole blood eQTL summary statistics from GTEx v8 [31] and the eQTLGen consortium [6]. After filtering for SNPs with RSIDs, the GTEx dataset retained 19,696 genes, and the eQTLGen dataset retained 19,250 genes (Supplementary Table 4). A comprehensive list of all data sources is provided in Supplementary Table 1.

Data harmonization involved combining summary statistics from the target study (BBJ or POPCELL), corresponding auxiliary ct-eQTL and tissue-level studies, and the 1000 Genomes Project Phase 3 reference panel [26] to ensure consistent effect alleles. We then computed population-specific and trans-ancestry LD scores for each gene, restricting manually to SNPs with MAF ≥ 0.05, using S-LDXR [68]. The final analysis datasets comprised approximately 12k genes for the East Asian studies and 10k genes for the African study (Supplementary Table 6). All data processing scripts are publicly available at https://github.com/LucaJiang/traceCB.

### Cell type deconvolution

To estimate cell type proportions in bulk tissue, we applied CIBERSORTx [28] to GTEx v8 whole blood RNA-seq data (TPM normalized), sourced from the GTEx Portal https://storage.googleapis.com/adult-gtex/bulk-gex/v8/rna-seq/tpms-by-tissue/gene_tpm_2017-06-05_v8_whole_blood.gct.gz. To optimize computational efficiency and the signal-to-noise ratio, we employed a curated subset of the LM22 signature matrix derived from the complete 11,845-gene reference [69]. This refined matrix was constructed by selecting the top 30 most differentially expressed genes (ranked by ANOVA F-statistics) for each of the 22 immune cell subtypes, resulting in a signature set of 620 unique genes.

Deconvolution was performed with batch correction mode and quantile normalization disabled using CIBERSORTx webpage https://cibersortx.stanford.edu/. The resulting individual-level cell-type fractions are provided in the Supplementary Data; mean proportions across the cohort are summarized in Supplementary Table 5 and visualized in Supplementary Fig. 1.

### Colocalization analysis

To identify shared causal variants between eQTL and GWAS signals, we performed colocalization analysis following the methodology established by Bryois et al. [45] We first identified leading GWAS SNPs as those with the most significant association signal (smallest *P*-value) exceeding 1 × 10^−6^ within each genomic region. Genomic regions were defined according to the CytoBand (hg19) annotation to ensure comprehensive coverage while avoiding overlap between adjacent regions. For each identified leading SNP, we mapped the nearest protein-coding gene using the Ensembl database annotation.

We subsequently employed the COLOC R package [35] to assess the posterior probability of shared causal variants within each locus via a Bayesian framework. This method evaluates five mutually exclusive hypotheses: H0 (no association with either trait), H1 (eQTL association only), H2 (GWAS association only), H3 (both traits associated but driven by distinct causal variants), and H4 (both traits associated and sharing a single causal variant). The corresponding posterior probabilities are denoted as P0 through P4, where 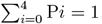. Loci were considered colocalized if P4 > 0.7, suggesting strong evidence for a common causal mechanism. Conversely, loci with P3 > 0.7 were classified as having independent signals, while all other loci were categorized as undetermined.

To investigate the functional context of colocalized signals, we assessed their overlap with active regulatory elements using cell-type-matched H3K27ac ChIP-seq data from the ENCODE project [70]. Specifically, we analyzed data for CD4^+^ T cells (ENCFF357NOB), CD8^+^ T cells (ENCFF455UVQ), B cells (ENCFF701BIL), NK cells (ENCFF473CXT), and monocytes (ENCFF840HBF), sourced from the ENCODE Portal https://www.encodeproject.org/.

### Runtime Analysis

We evaluated the computational efficiency of traceCB across 10 studies and all 22 autosomes with BBJ, eQTL Catalogue and eQTLGen data as described in the previous sections (Supplementary Fig. 58). The analysis was performed on a server equipped with double AMD EPYC 9754 128-Core Processors. On average, processing all 22 autosomes for a single study required 3,201.7 seconds of aggregated CPU time. However, traceCB supports parallel execution at the chromosome level, substantially reducing CPU time. For instance, in the BLUEPRINT (167) study, the maximum runtime for a single chromosome was 432 seconds (chromosome 1), corresponding to an average of 0.3 seconds per gene (432 seconds / 1,338 genes). Assuming sufficient computational resources are available to process all 22 chromosomes in parallel, the effective CPU time would be limited by the longest chromosome, reducing the total analysis time to approximately 7.2 minutes per study. This demonstrates traceCB’s high scalability and computational feasibility for large-scale trans-ancestry ct-eQTL mapping.

## Data Availability

The individual-level cell type proportions estimated in this study are provided in the Supplementary Data. Publicly available datasets analyzed in this study are listed below:

- **ct-eQTL summary statistics**:
  - BBJ eQTL (EAS; CD4^+^T cells, CD8^+^ T cells, B cells, NK cells, Monocytes): https://humandbs.dbcls.jp/en/hum0099-v1
  - OASIS (EAS): https://humandbs.biosciencedbc.jp/en/hum0197-latest
  - CIMA (EAS): https://db.cngb.org/trueblood/cima/
  - Onek1k (EUR): https://onek1k.org/
  - eQTL Catalogue (EUR; Studies QTD000021, QTD000031, QTD000066, QTD000067, QTD000069, QTD000073, QTD000081, QTD000115, QTD000371, QTD000372): https://www.ebi.ac.uk/eqtl/
  - Popcell (80 Africans, non-stimulated condition): https://doi.org/10.1038/s41586-023-06422-9

- **Tissue eQTL summary statistics**:
  - GTEx eQTL (EUR; Whole Blood): https://www.gtexportal.org
  - eQTLGen (EUR; Full *cis*-eQTL summary statistics): https://eqtlgen.org/cis-eqtls.html
  - Wang et al. (EAS; hum0343.v3.eqtl.v1): https://humandbs.dbcls.jp/en/hum0343-v4

- **GWAS summary statistics**:
  - BBJ GWAS (EAS; Asthma, Atopic dermatitis, Rheumatoid arthritis): http://jenger.riken.jp/result
  - Blood Cell Consortium (EAS; Red blood cells, White blood cells, Platelets): http://www.mhi-humangenetics.org/en/resources/

- **Other data**:
  - 1000 Genomes Project Phase 3 genotype data (EAS, EUR, AFR): https://www.cog-genomics.org/plink/1.9/resources
  - GTEx RNA-Seq individual-level bulk gene expression (EUR; Whole Blood): https://www.gtexportal.org
  - CIBERSORTx Reference (GEPIA2021 LM22): https://github.com/zwj-tina/GEPIA2021/blob/main/reference/LM22.txt
  - ENCODE H3K27ac ChIP-seq data including CD4^+^ T cells (ENCFF357NOB), CD8^+^ T cells (ENCFF455UVQ), B cells (ENCFF701BIL), NK cells (ENCFF473CXT), and monocytes (ENCFF840HBF): ENCODE Portal https://www.encodeproject.org/.

## Code Availability

traceCB is an open-source Python package available at https://github.com/LucaJiang/traceCB. Analysis scripts for reproducing the results are available at the same repository.

## Acknowledgments

M.C. was supported in part by National Natural Science Foundation of China [12501402], Hong Kong Research Grant Council [21305525], Guangdong Basic and Applied Basic Research Foundation via Shenzhen Research Institute at City University of Hong Kong [2026A1515010402, 2026A1515060010], and City University of Hong Kong [21300423]. J.X. was supported by National Natural Science Foundation of China [12401384], Guangdong Natural Science Foundation General Project [2025A1515011603], and Sun Yat-sen University Startup Grant [2026_51000_B26833]. This research was conducted using the UK Biobank resource under application number 96744. We thank Xinyi Yu for providing resources regarding IBSEP. During the preparation of this work, the authors used Gemini 3.0 Pro for language editing and to refine the manuscript’s clarity. Additionally, initial drafts of specific computational code components were generated with the assistance of Gemini 3.0 Pro, and subsequently subjected to rigorous manual verification, debugging, and validation by the authors to ensure technical accuracy. The authors reviewed and edited the final content as needed and take full responsibility for the results and integrity of the work.

