## Supplementary figures and note for "traceCB: Trans-ancestry cell-type-specific eQTLs mapping by integrating scRNA-seq and bulk data"

**2 Supplementary Tables** **50**

**3 Supplementary Notes** **56**
3.1 Approximation of tissue-level marginal effects using mean cell-type proportions . 56

### 1 Supplementary Figures

#### 1.1 Estimated cell-type proportions

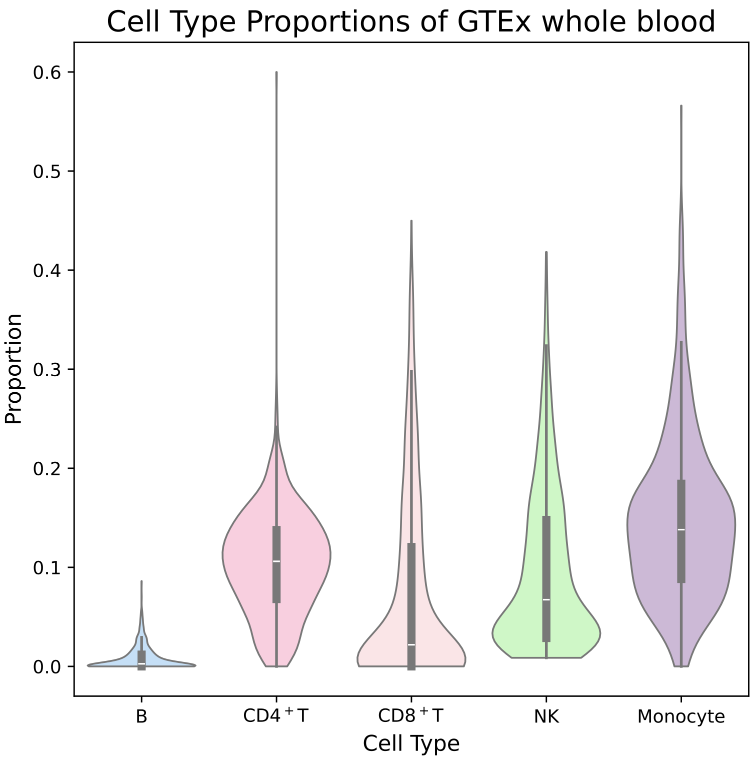

**Supplementary Figure 1:** Estimated cell-type proportions in GTEx whole blood samples using CIBERSORTx.

#### 1.2 Additional simulation results

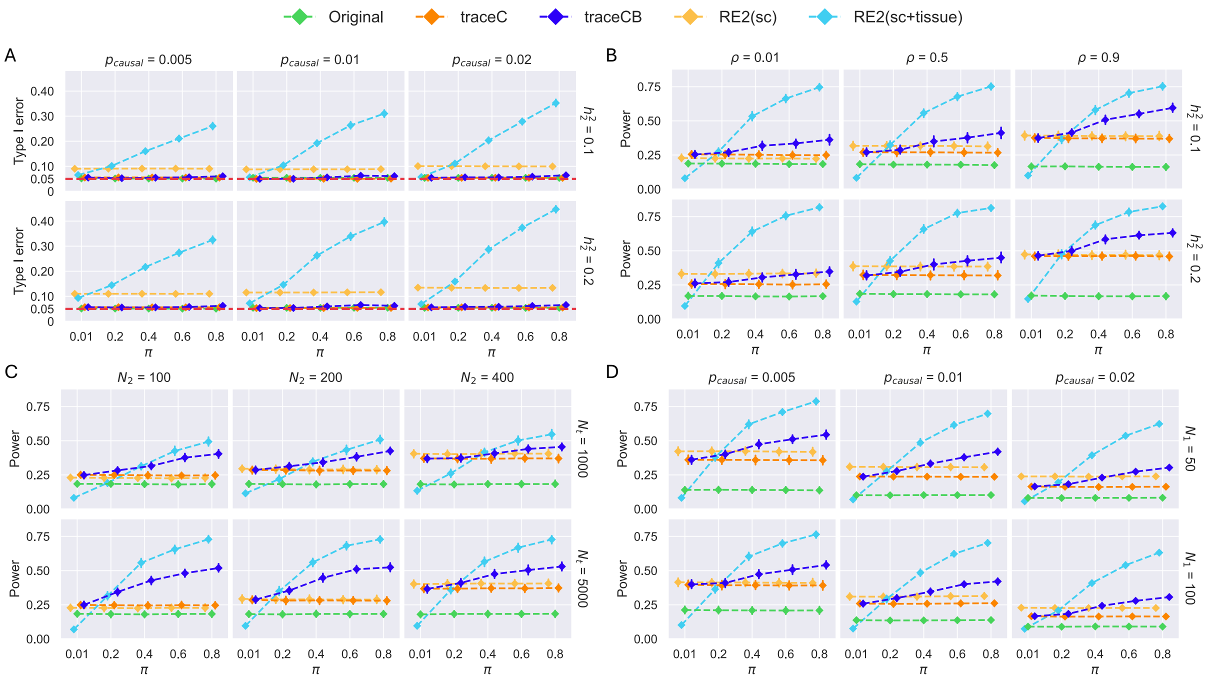

**Supplementary Figure 2:** Evaluation of type I error rate and statistical power under simulated scenarios where the ground truth of heritabilities and co-heritabilities are known. We compared four methods: "Original", which uses only the target population's scRNA-seq summary statistics; "traceC", a simplified version of traceCB without bulk tissue data; "RE2(sc)", a random effects meta-analysis of trans-ancestry ct-eQTLs; and "RE2(sc+tissue)", which extends RE2(sc) to include bulk eQTLs. Default settings are the same with those in Fig. 2 except that the true parameters are used in traceC and traceCB. All results are averaged over 100 replicates, and error bars represent 95% confidence intervals.

#### 1.3 Colocalization Results

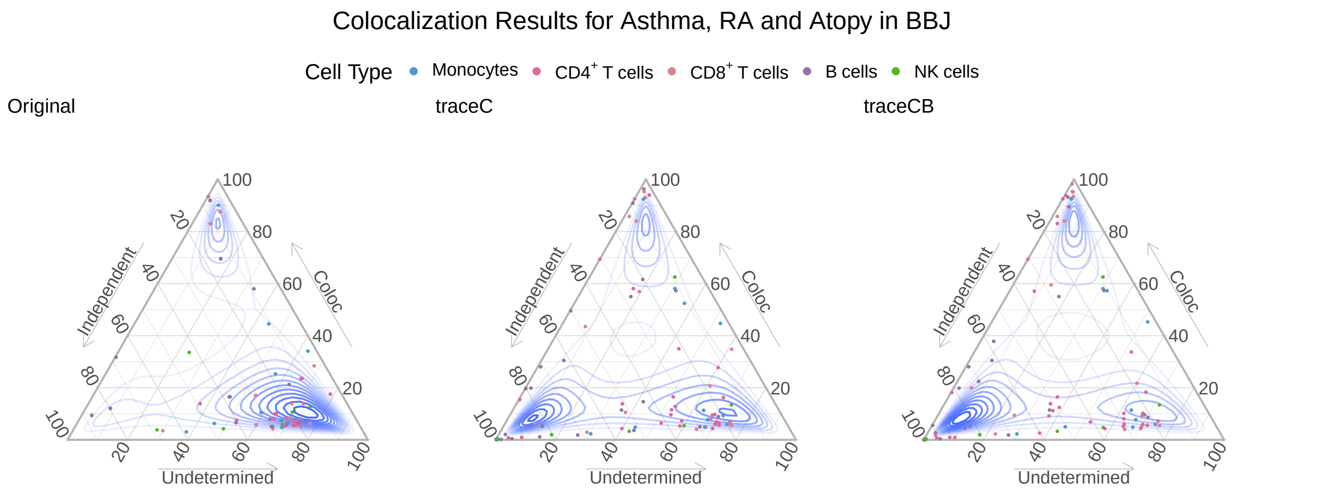

**Supplementary Figure 3:** Colocalization analysis with GWAS of 3 immune-related traits from BBJ (Atopy: atopic dermatitis, RA: Rheumatoid Arthritis). Ternary plots illustrate colocalization probabilities for the original, traceC, and traceCB analyses using BBJ, eQTL Catalogue, and eQTLGen eQTLs. Background contours represent point density.

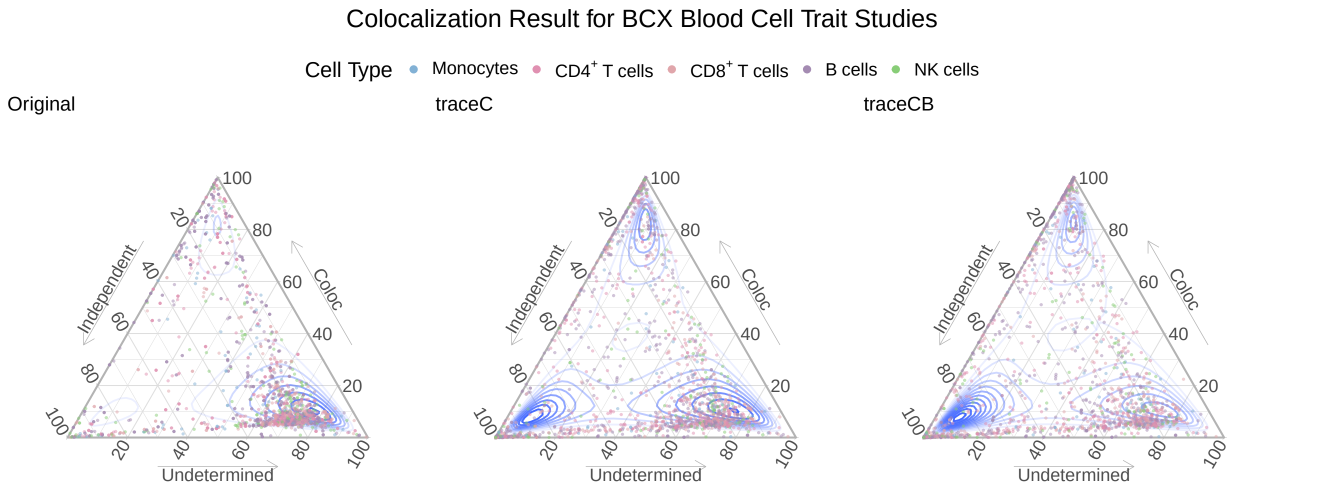

**Supplementary Figure 4:** Colocalization analysis with GWAS of 14 blood-related traits from BCX (red blood cell count, hemoglobin concentration, hematocrit, mean corpuscular hemoglobin, mean corpuscular volume, mean corpuscular hemoglobin concentration, white blood cell count, neutrophil count, lymphocyte count, monocyte count, basophil count, eosinophil count, platelet count, and mean platelet volume). Ternary plots illustrate colocalization probabilities for the original, traceC, and traceCB analyses using BBJ, eQTL Catalogue, and eQTLGen eQTLs. Background contours represent point density.

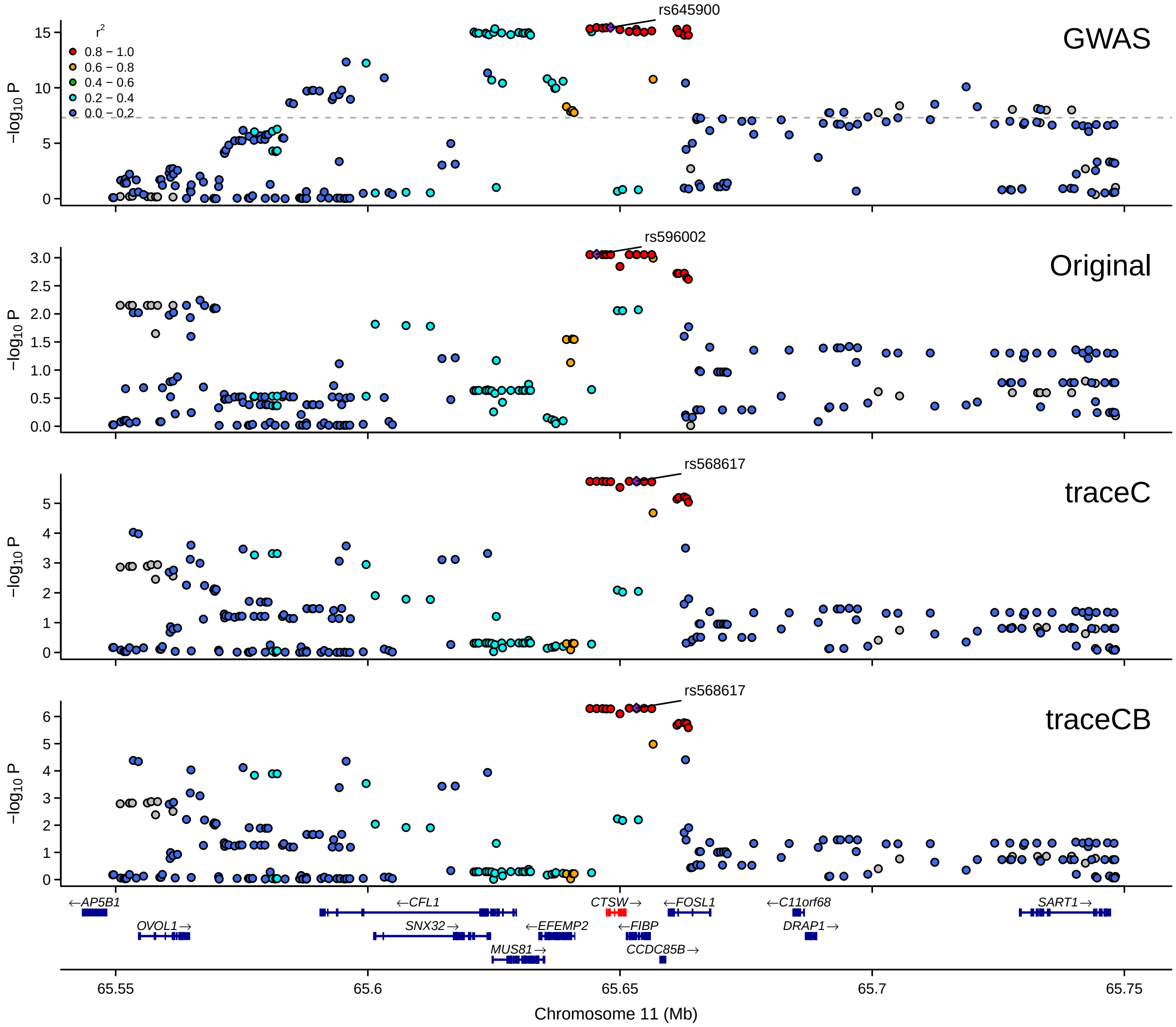

**Supplementary Figure 5:** LocusZoom plot for *CTSW* in prioritizing BBJ monocyte eQTL with CEDAR (286) and eQTLGen. GWAS summary statistics are from BCX monocyte count.

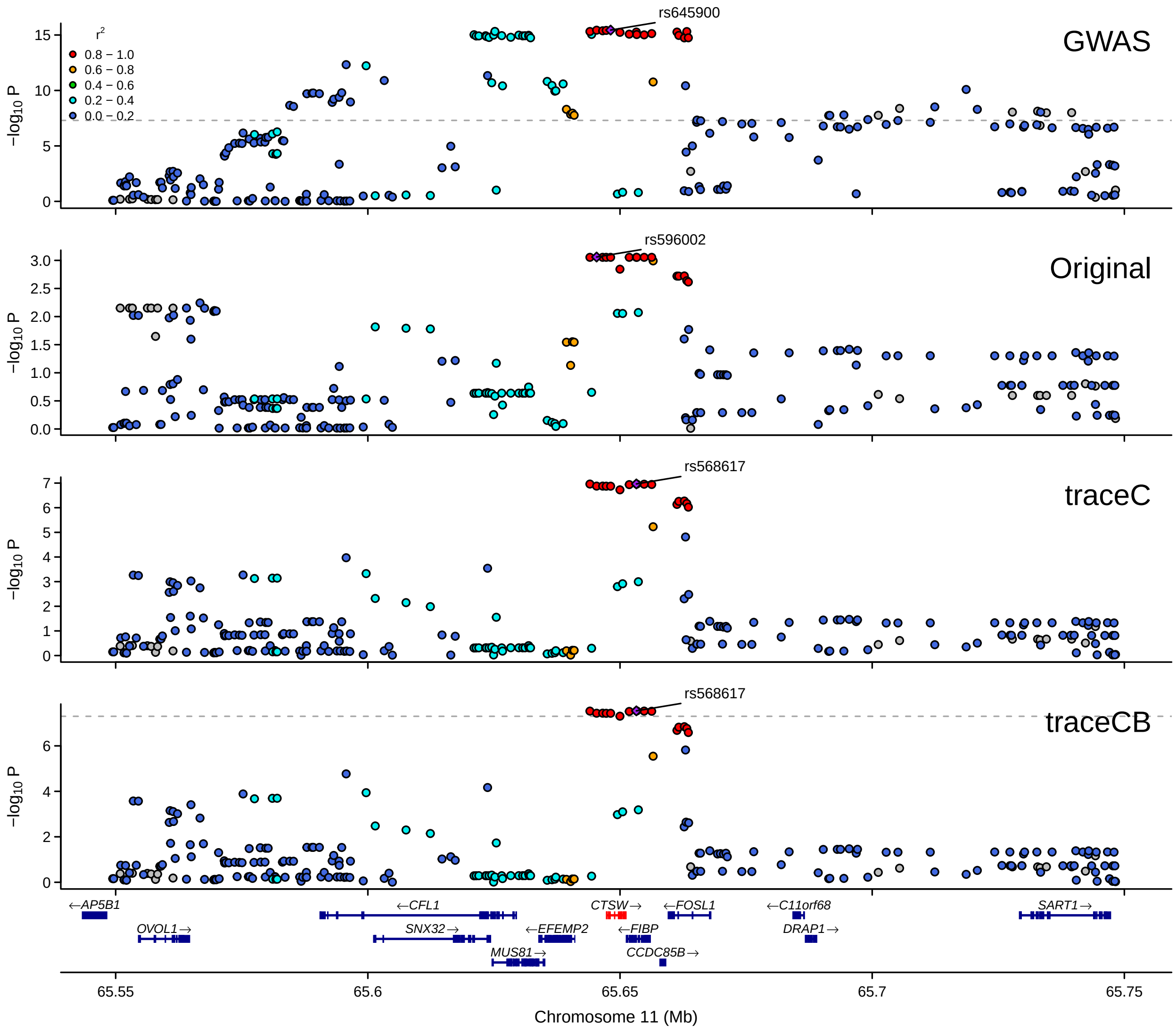

**Supplementary Figure 6:** LocusZoom plot for *CTSW* in prioritizing BBJ monocyte eQTL with Fairfax_2014 (420) and eQTLGen. GWAS summary statistics are from BCX monocyte count.

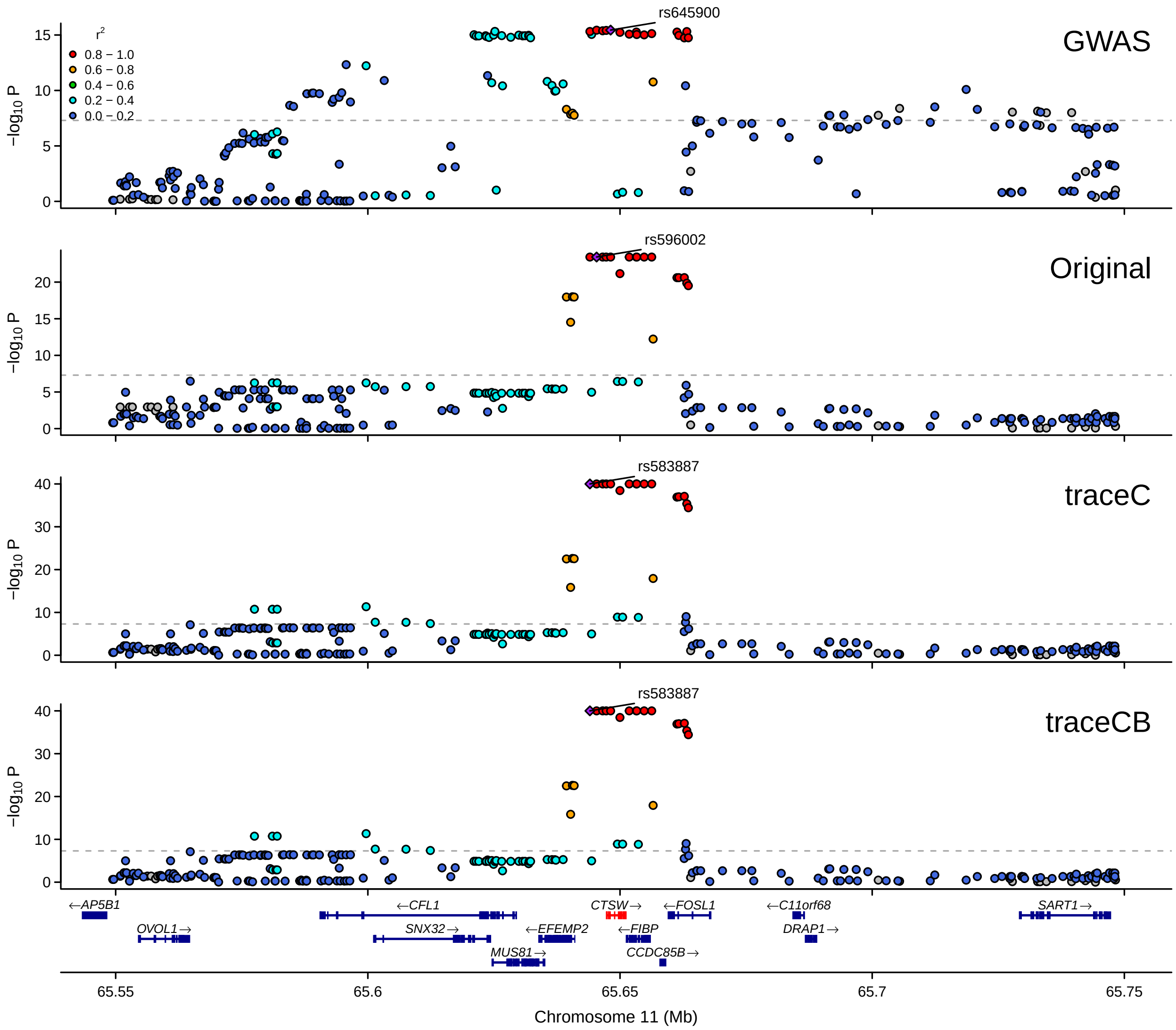

**Supplementary Figure 7:** LocusZoom plot for *CTSW* in prioritizing BBJ $CD4^+$ T cell eQTL with BLUEPRINT (167) and eQTLGen. GWAS summary statistics are from BCX monocyte count. Minimum $P$-value is set to $10^{-40}$.

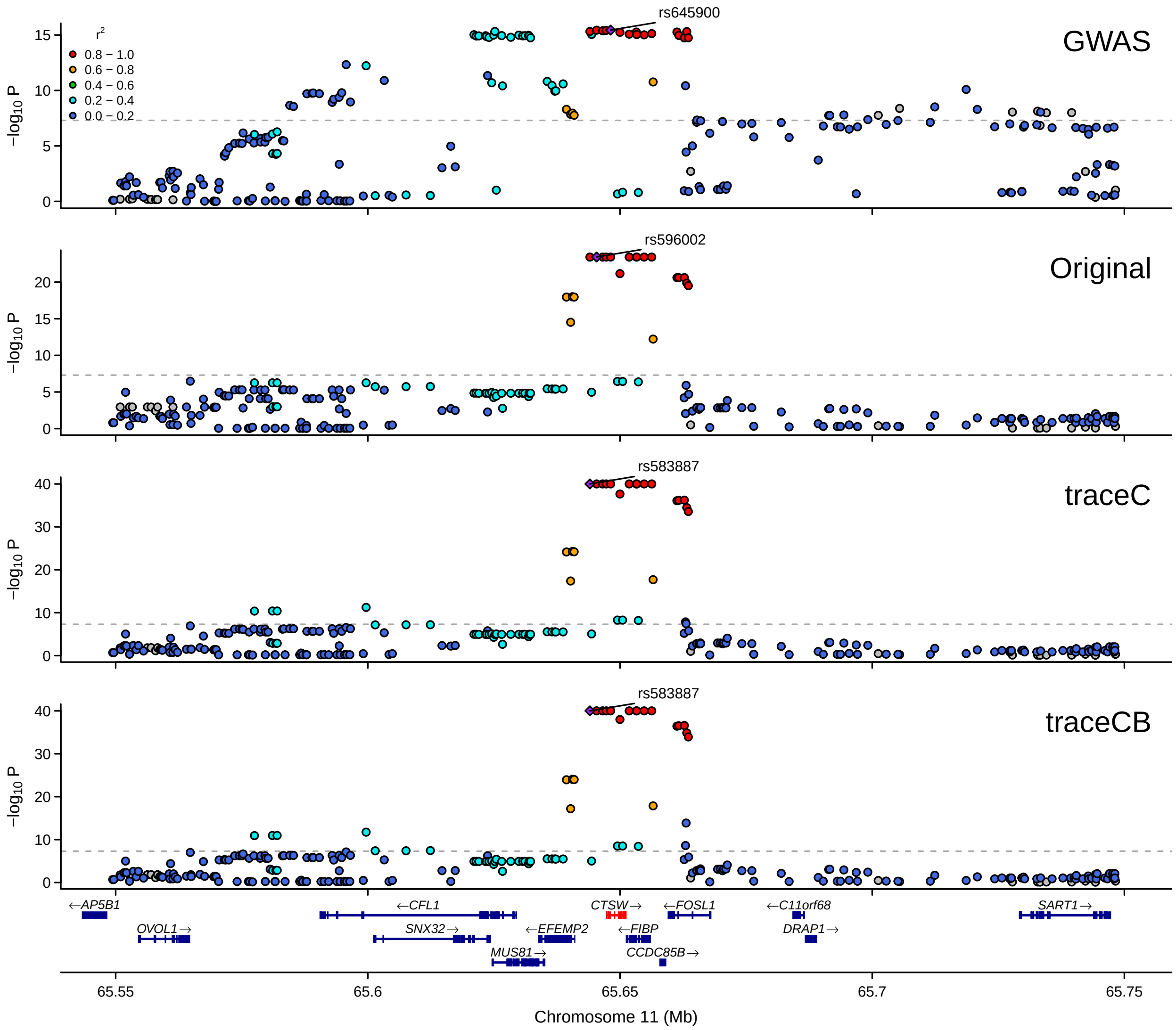

**Supplementary Figure 8:** LocusZoom plot for *CTSW* in prioritizing BBJ CD4$^{+}$ T cell eQTL with CEDAR (290) and eQTLGen. GWAS summary statistics are from BCX monocyte count. Minimum $P$-value is set to $10^{-40}$.

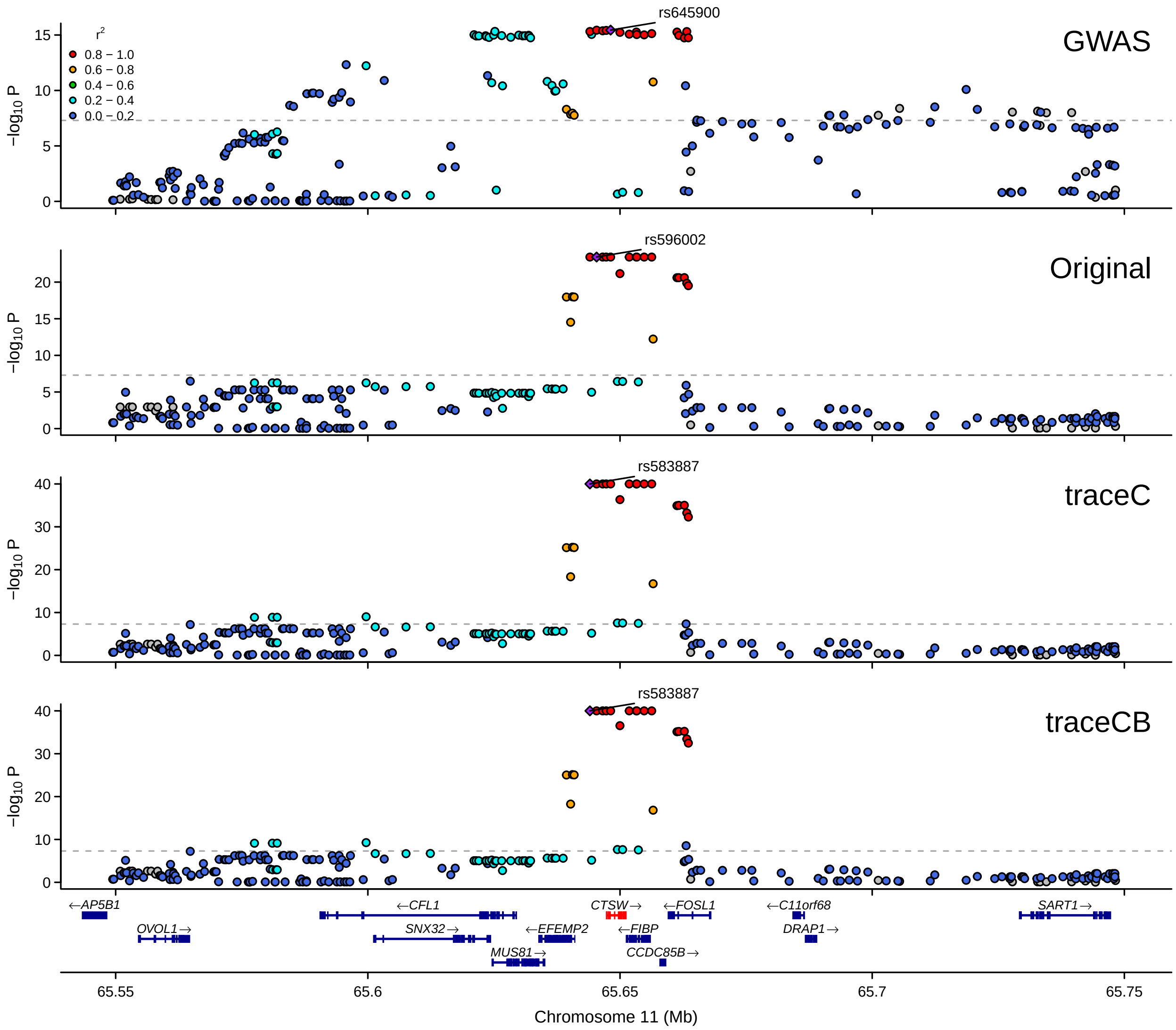

**Supplementary Figure 9:** LocusZoom plot for *CTSW* in prioritizing BBJ CD4$^+$ T cell eQTL with Kasela_2017(280) and eQTLGen. GWAS summary statistics are from BCX monocyte count. Minimum $P$-value is set to $10^{-40}$.

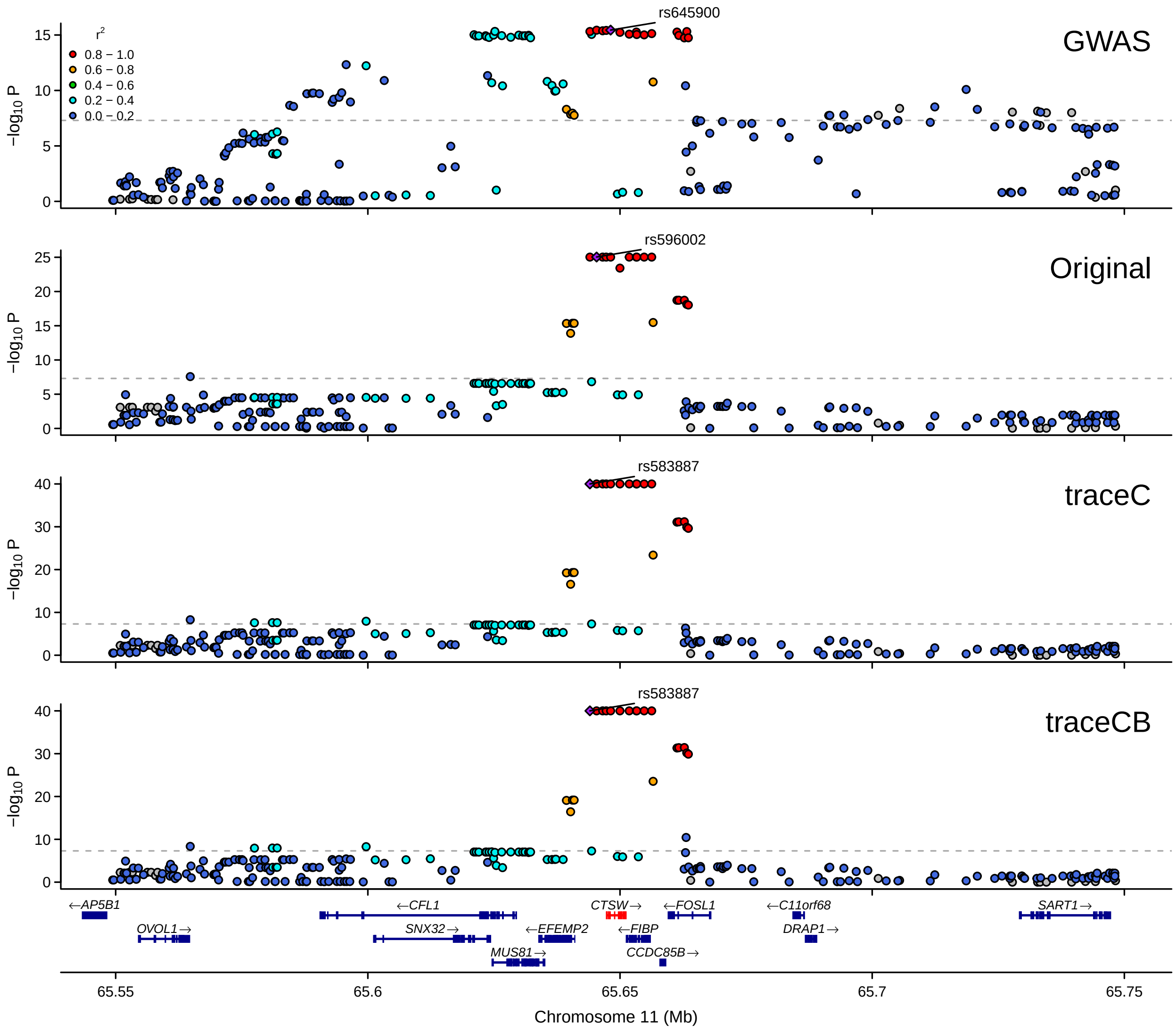

**Supplementary Figure 10:** LocusZoom plot for *CTSW* in prioritizing BBJ CD8$^+$ T cell eQTL with CEDAR (277) and eQTLGen. GWAS summary statistics are from BCX monocyte count. Minimum $P$-value is set to $10^{-40}$.

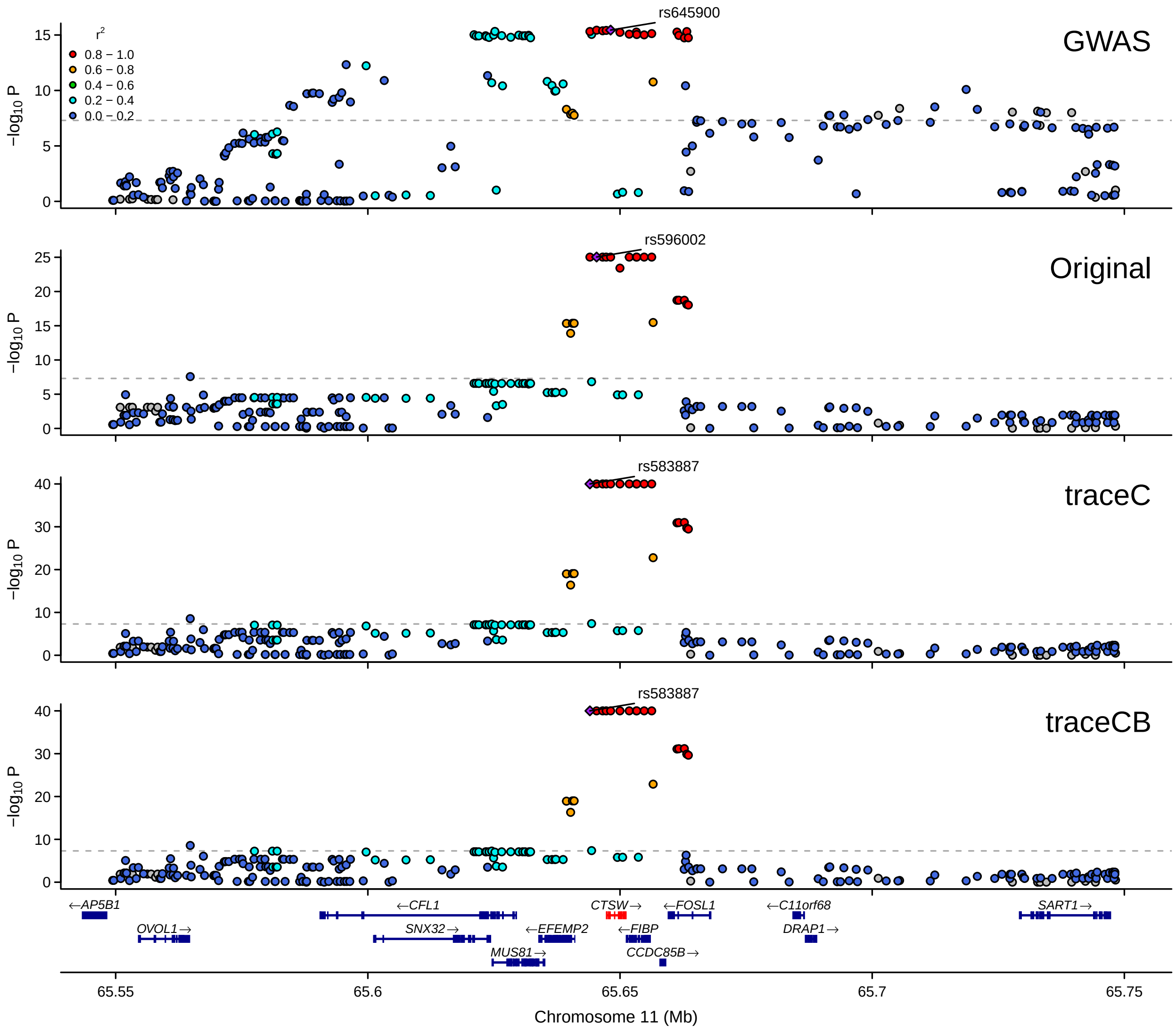

**Supplementary Figure 11:** LocusZoom plot for *CTSW* in prioritizing BBJ CD8$^{+}$ T cell eQTL with Kasela_2017(269) and eQTLGen. GWAS summary statistics are from BCX monocyte count. Minimum $P$-value is set to $10^{-40}$.

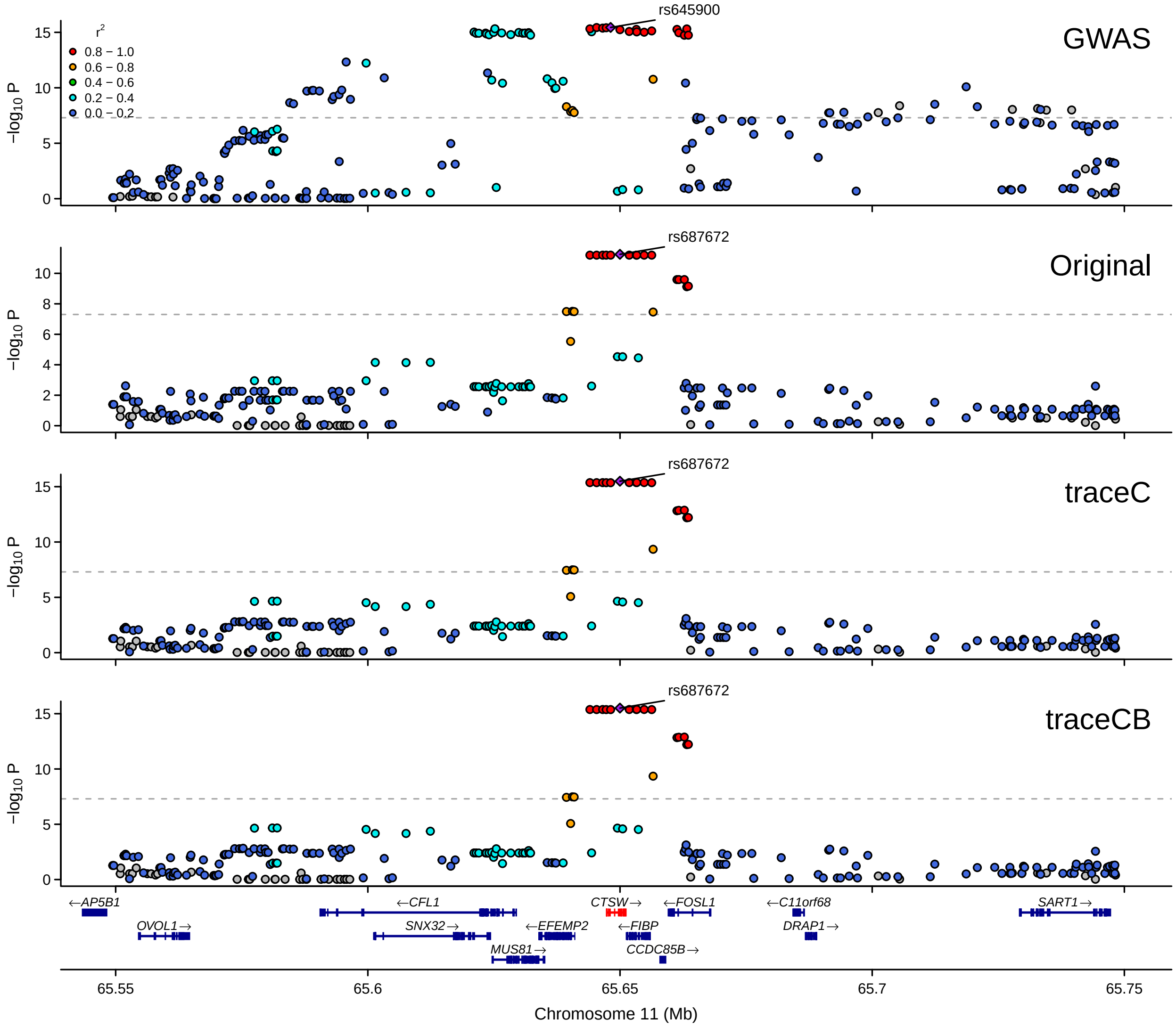

**Supplementary Figure 12:** LocusZoom plot for *CTSW* in prioritizing BBJ B cell eQTL with CEDAR (262) and eQTLGen. GWAS summary statistics are from BCX monocyte count.

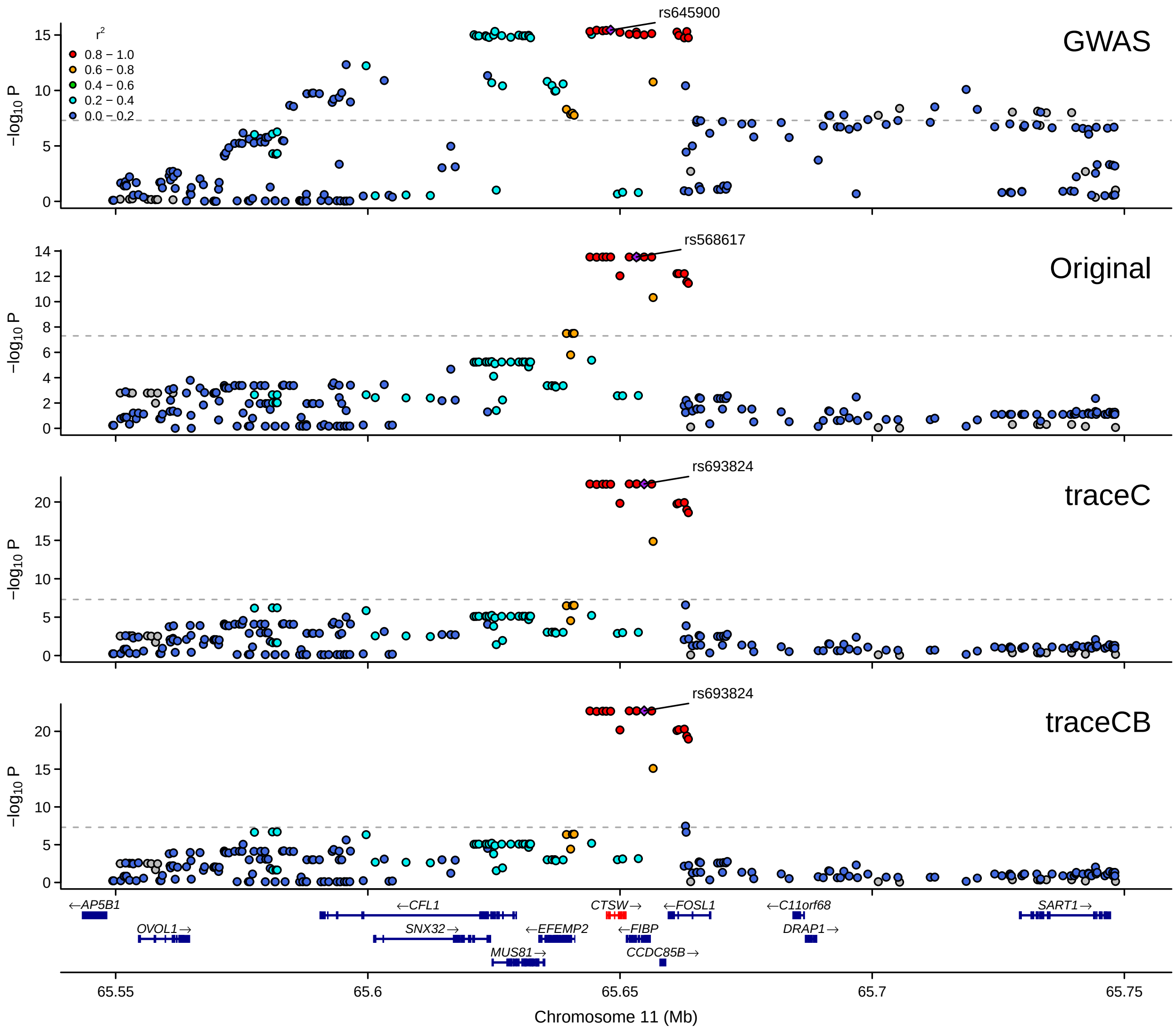

**Supplementary Figure 13:** LocusZoom plot for *CTSW* in prioritizing BBJ NK cell eQTL with Gilchrist_2021 (247) and eQTLGen. GWAS summary statistics are from BCX monocyte count.

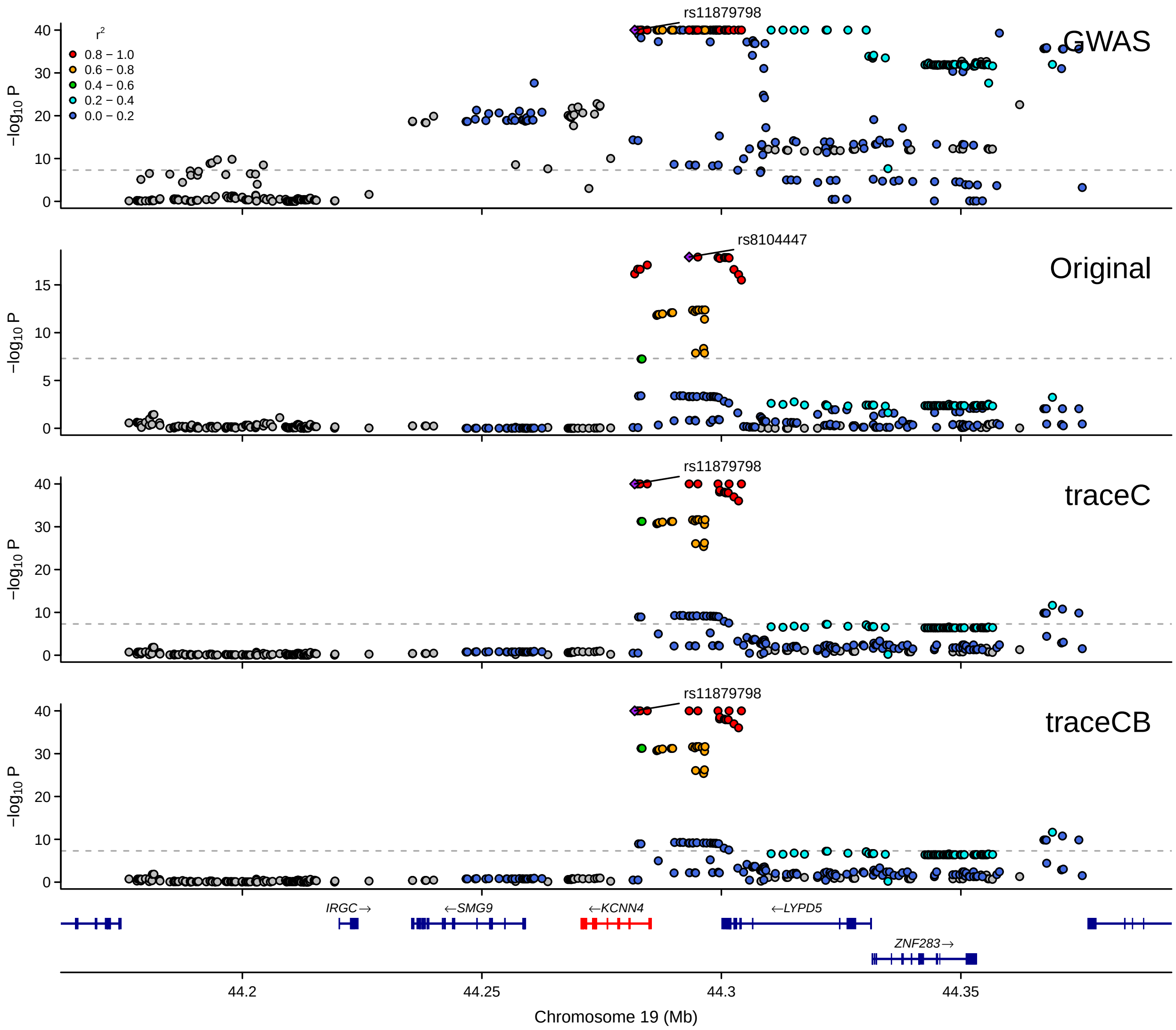

**Supplementary Figure 14:** LocusZoom plot for *KCNN4* in prioritizing BBJ monocyte eQTL with BLUEPRINT (191) and eQTLGen. GWAS summary statistics are from BCX monocyte count. Minimum $P$-value is set to $10^{-40}$.

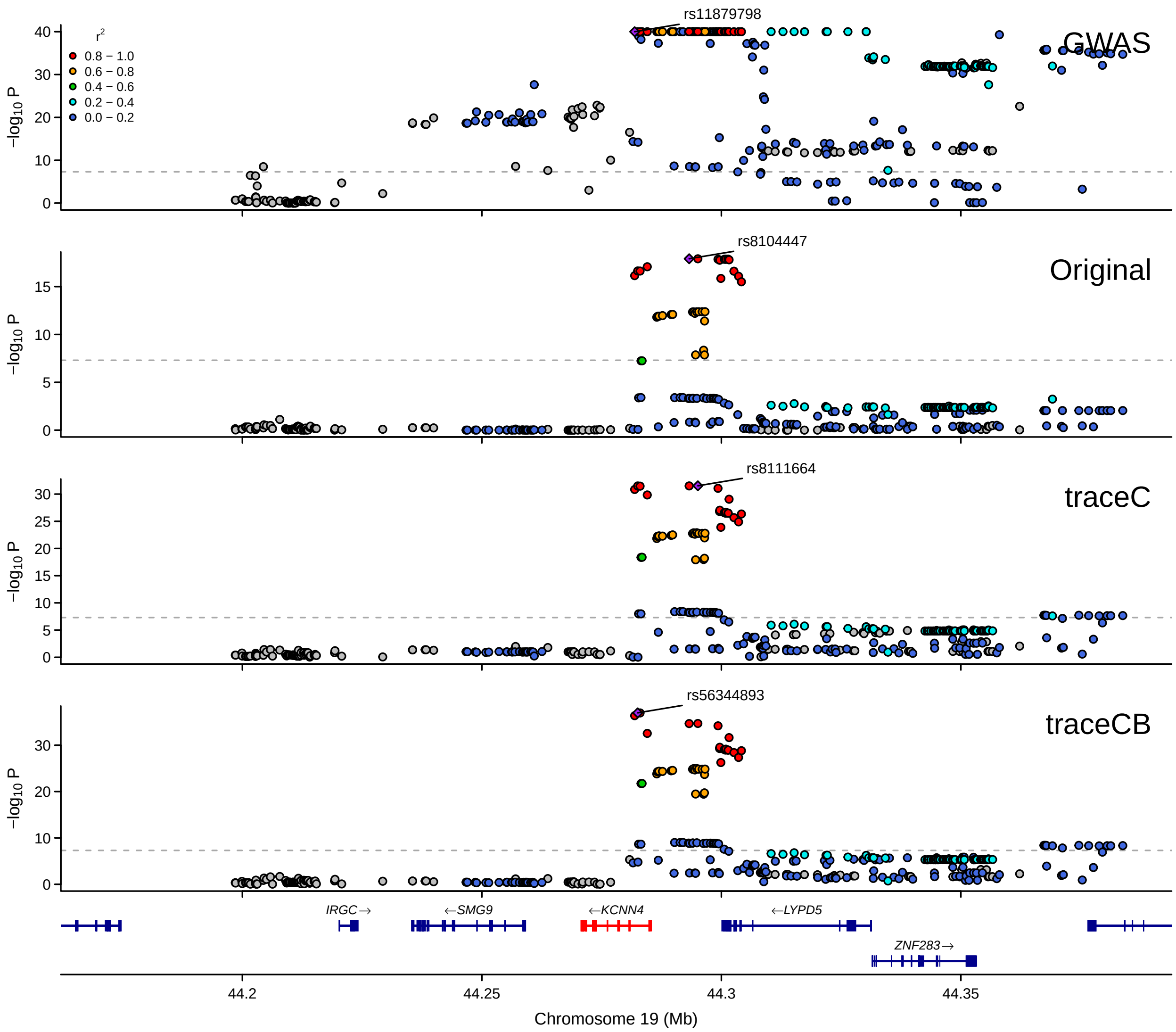

**Supplementary Figure 15:** LocusZoom plot for *KCNN4* in prioritizing BBJ monocyte eQTL with CEDAR (286) and eQTLGen. GWAS summary statistics are from BCX monocyte count.

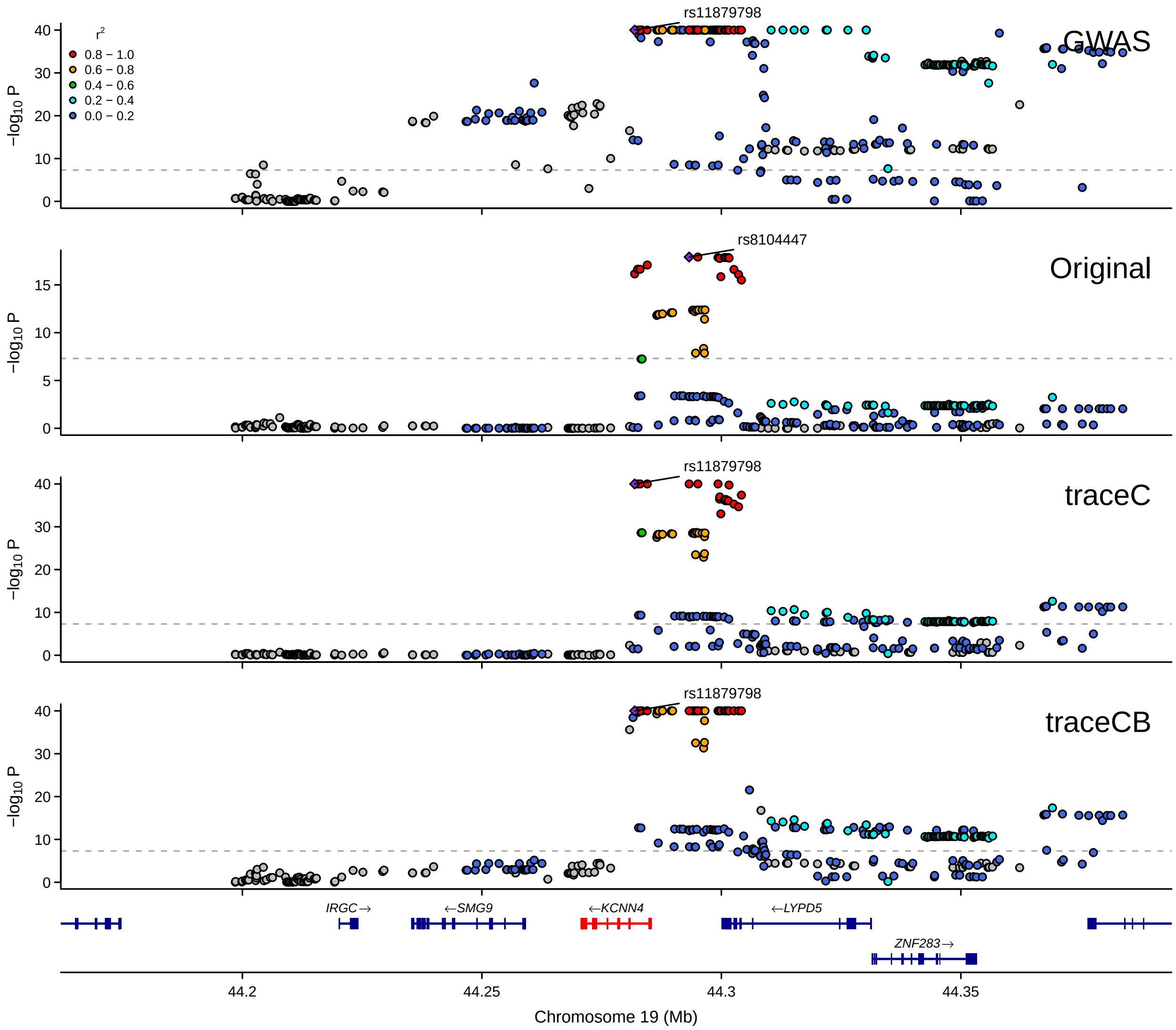

**Supplementary Figure 16:** LocusZoom plot for *KCNN4* in prioritizing BBJ monocyte eQTL with Fairfax_2014 (420) and eQTLGen. GWAS summary statistics are from BCX monocyte count. Minimum $P$-value is set to $10^{-40}$.

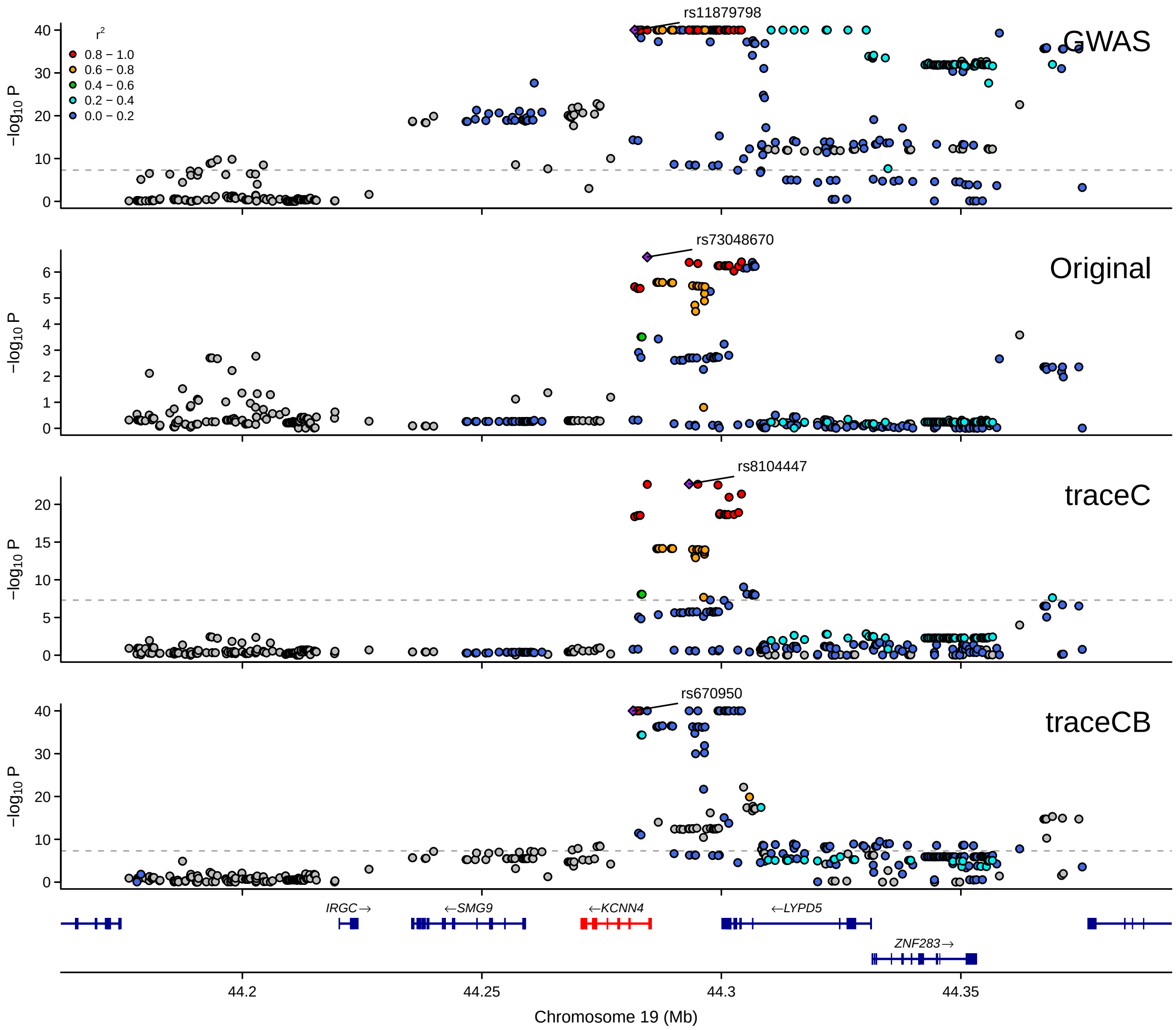

**Supplementary Figure 17:** LocusZoom plot for *KCNN4* in prioritizing BBJ CD4$^{+}$ T cell eQTL with BLUEPRINT (167) and eQTLGen. GWAS summary statistics are from BCX monocyte count. Minimum $P$-value is set to $10^{-40}$.

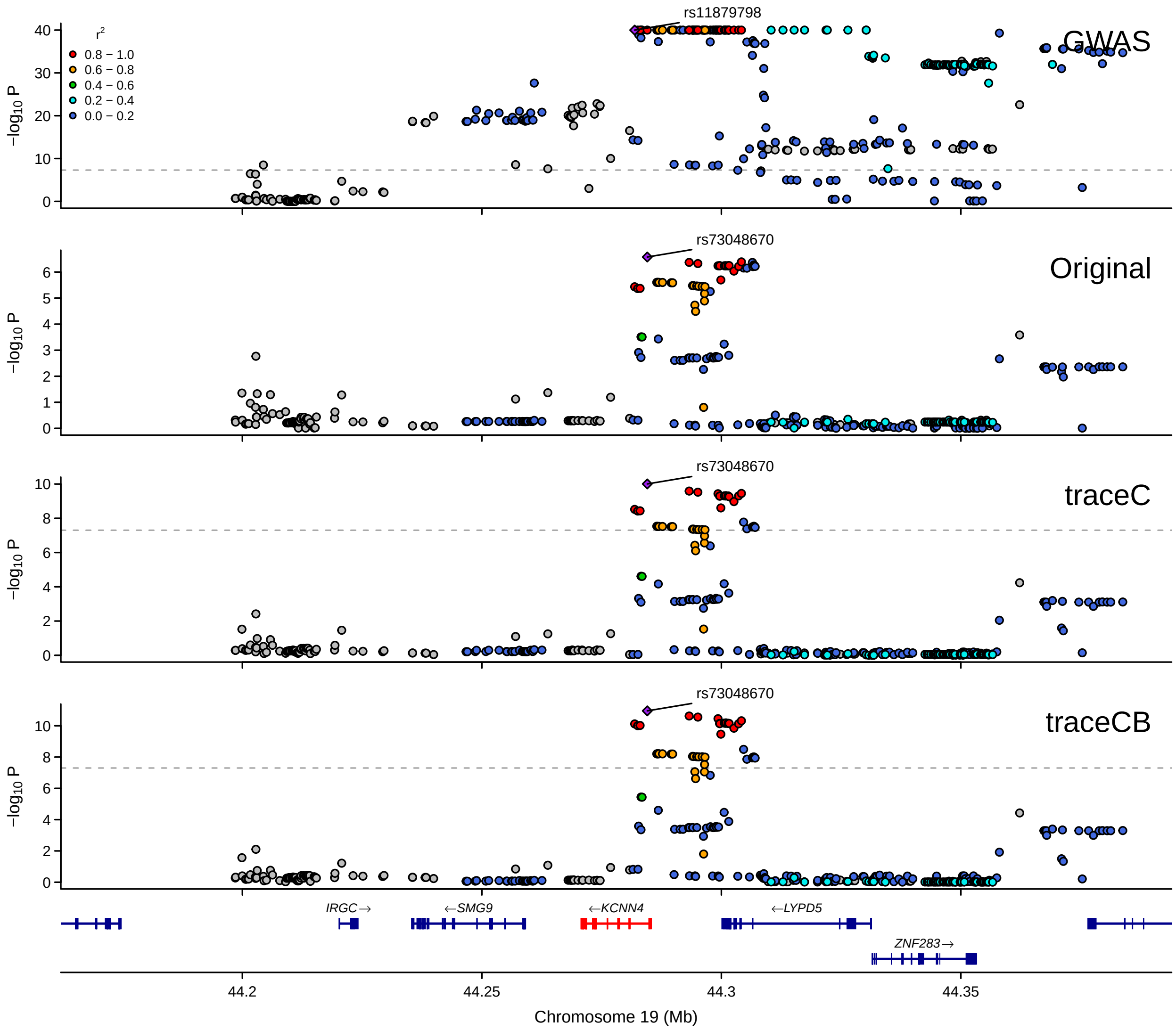

**Supplementary Figure 18:** LocusZoom plot for *KCNN4* in prioritizing BBJ CD4$^+$ T cell eQTL with Kasela_2017(280) and eQTLGen. GWAS summary statistics are from BCX monocyte count.

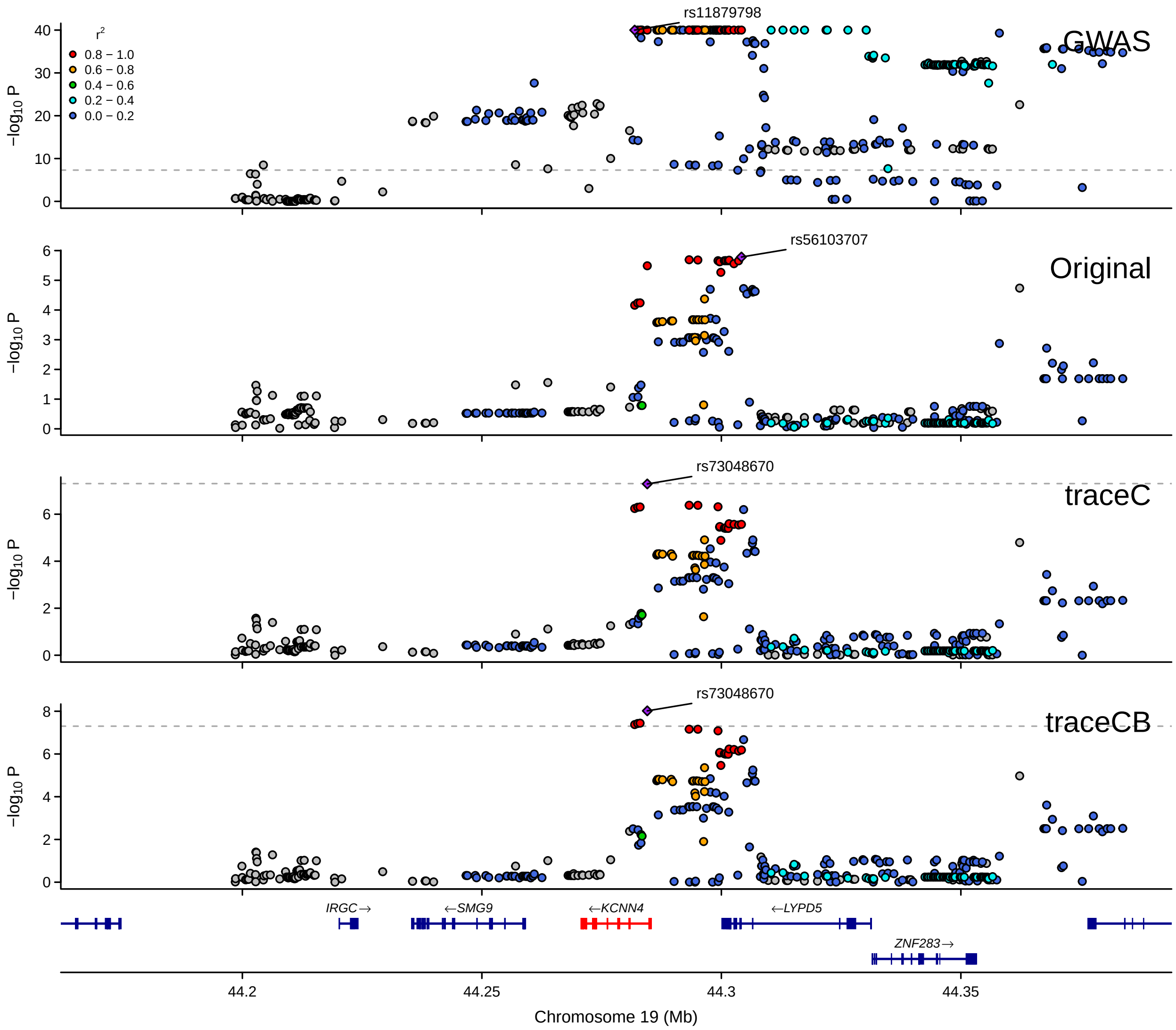

**Supplementary Figure 19:** LocusZoom plot for *KCNN4* in prioritizing BBJ CD8$^{+}$ T cell eQTL with CEDAR (277) and eQTLGen. GWAS summary statistics are from BCX monocyte count.

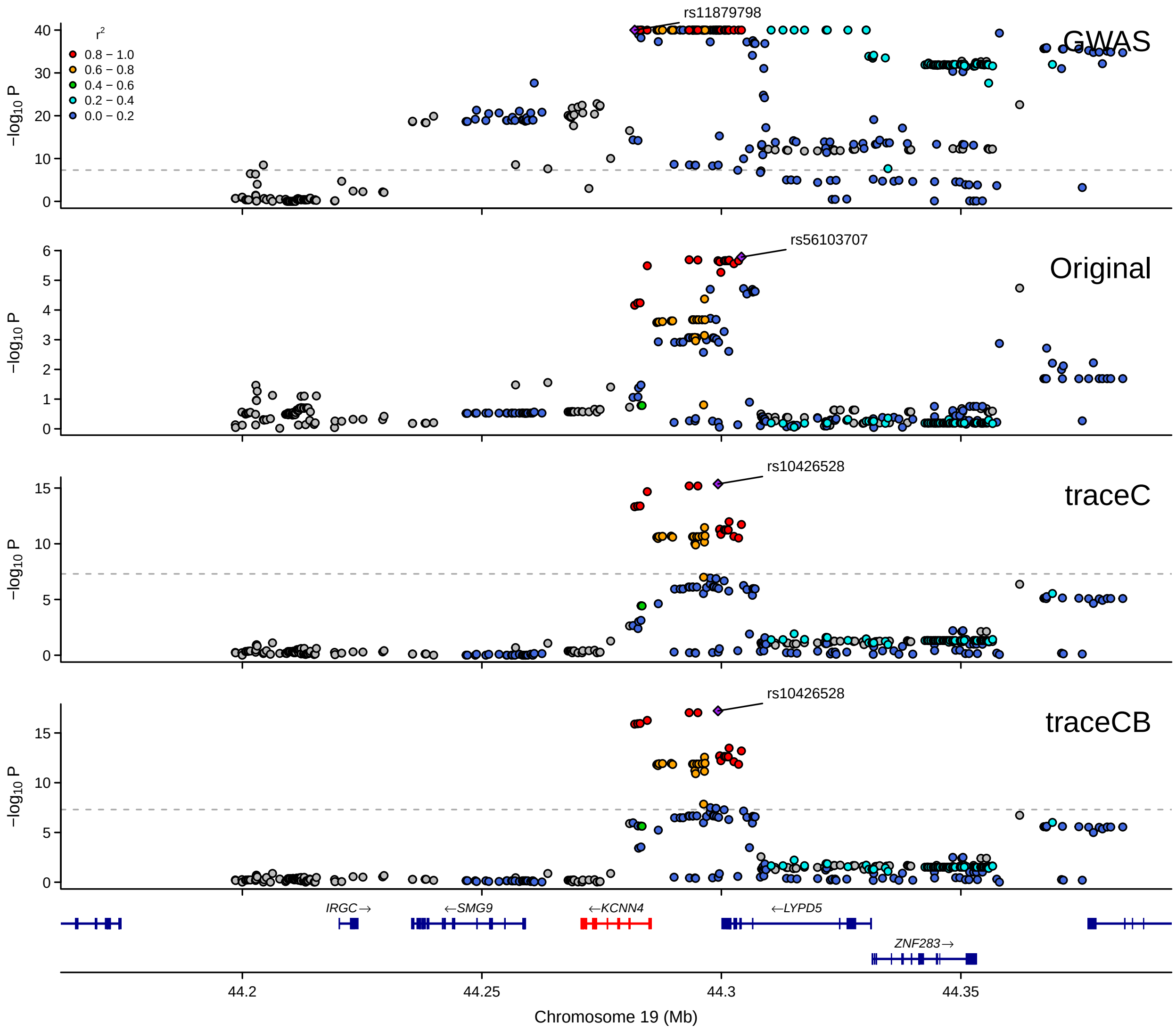

**Supplementary Figure 20:** LocusZoom plot for *KCNN4* in prioritizing BBJ CD8$^{+}$ T cell eQTL with Kasela_2017(269) and eQTLGen. GWAS summary statistics are from BCX monocyte count.

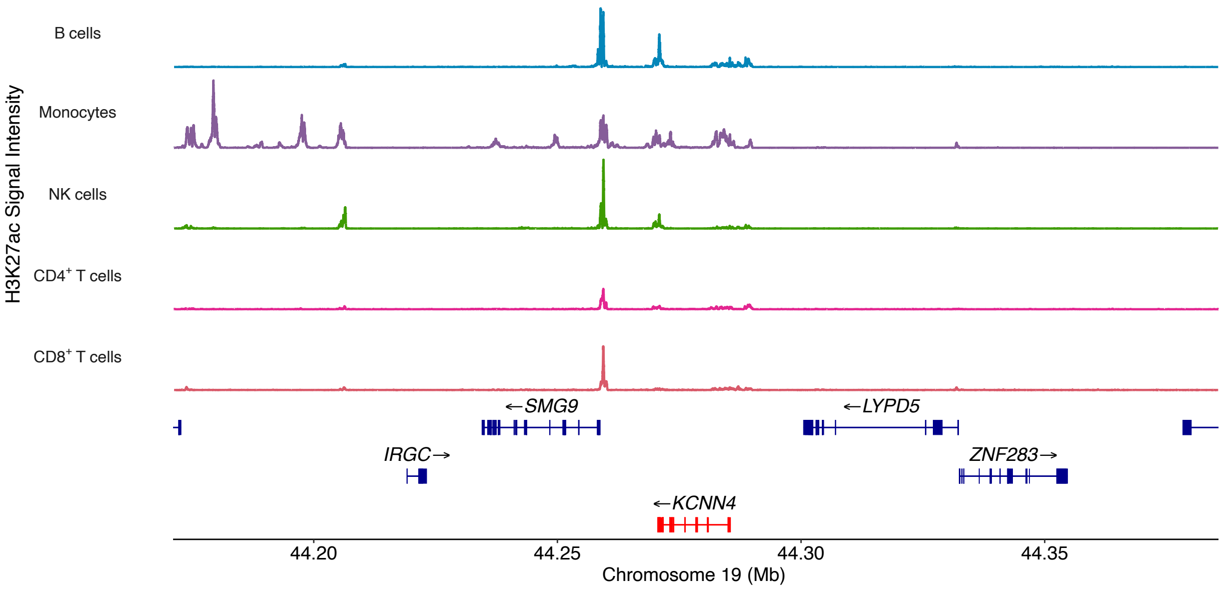

**Supplementary Figure 21:** H3K27ac signal for *KCNN4* in 5 immune cell types.

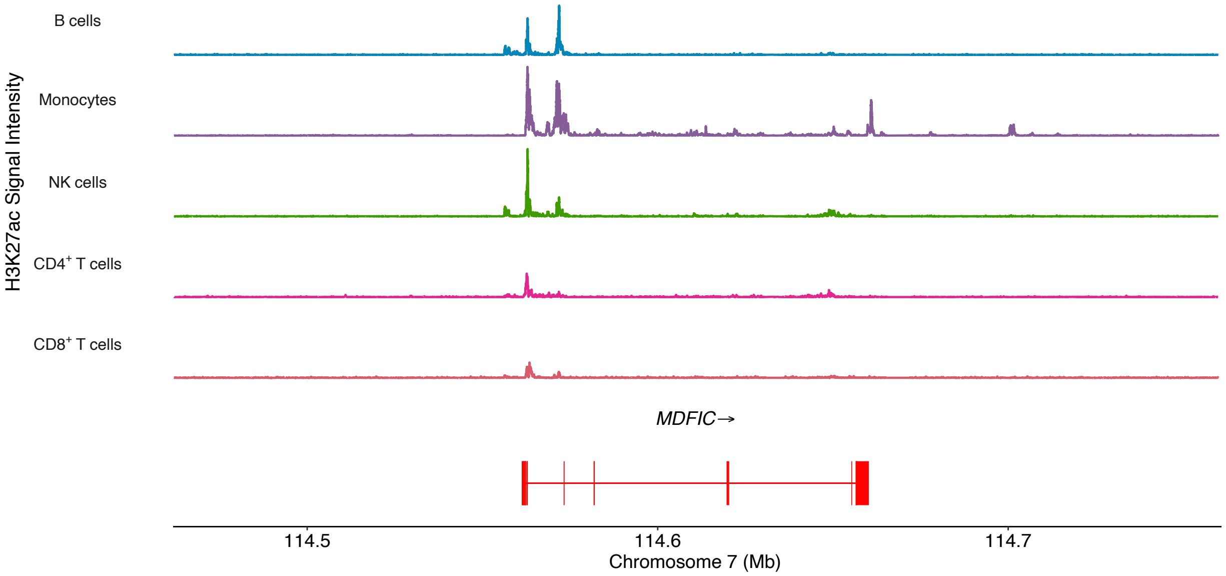

**Supplementary Figure 22:** H3K27ac signal for *MDFIC* in 5 immune cell types.

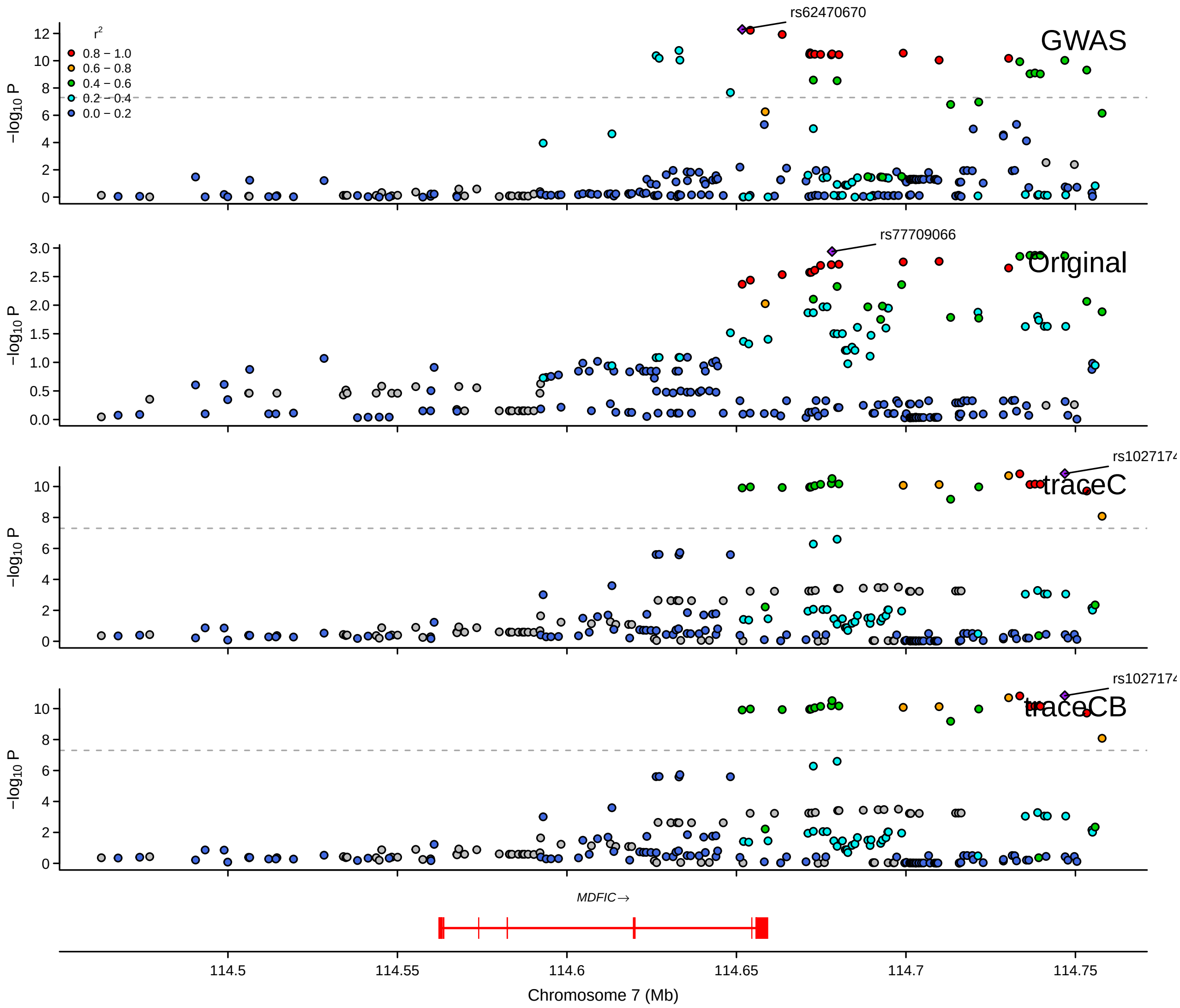

**Supplementary Figure 23:** LocusZoom plot for *MDFIC* in prioritizing BBJ monocyte eQTL with BLUEPRINT (191) and eQTLGen. GWAS summary statistics are from BCX monocyte count.

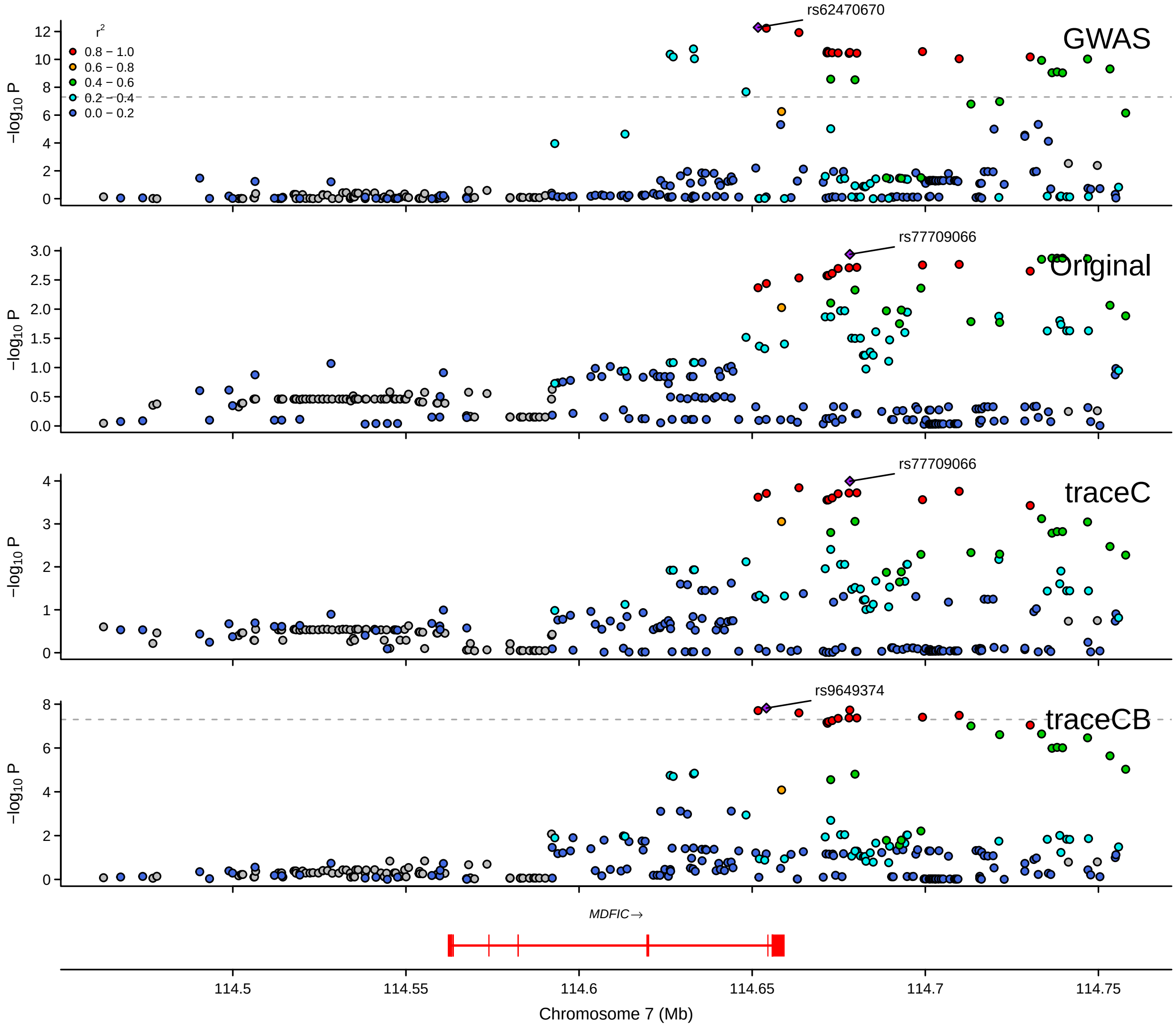

**Supplementary Figure 24:** LocusZoom plot for *MDFIC* in prioritizing BBJ monocyte eQTL with CEDAR (286) and eQTLGen. GWAS summary statistics are from BCX monocyte count.

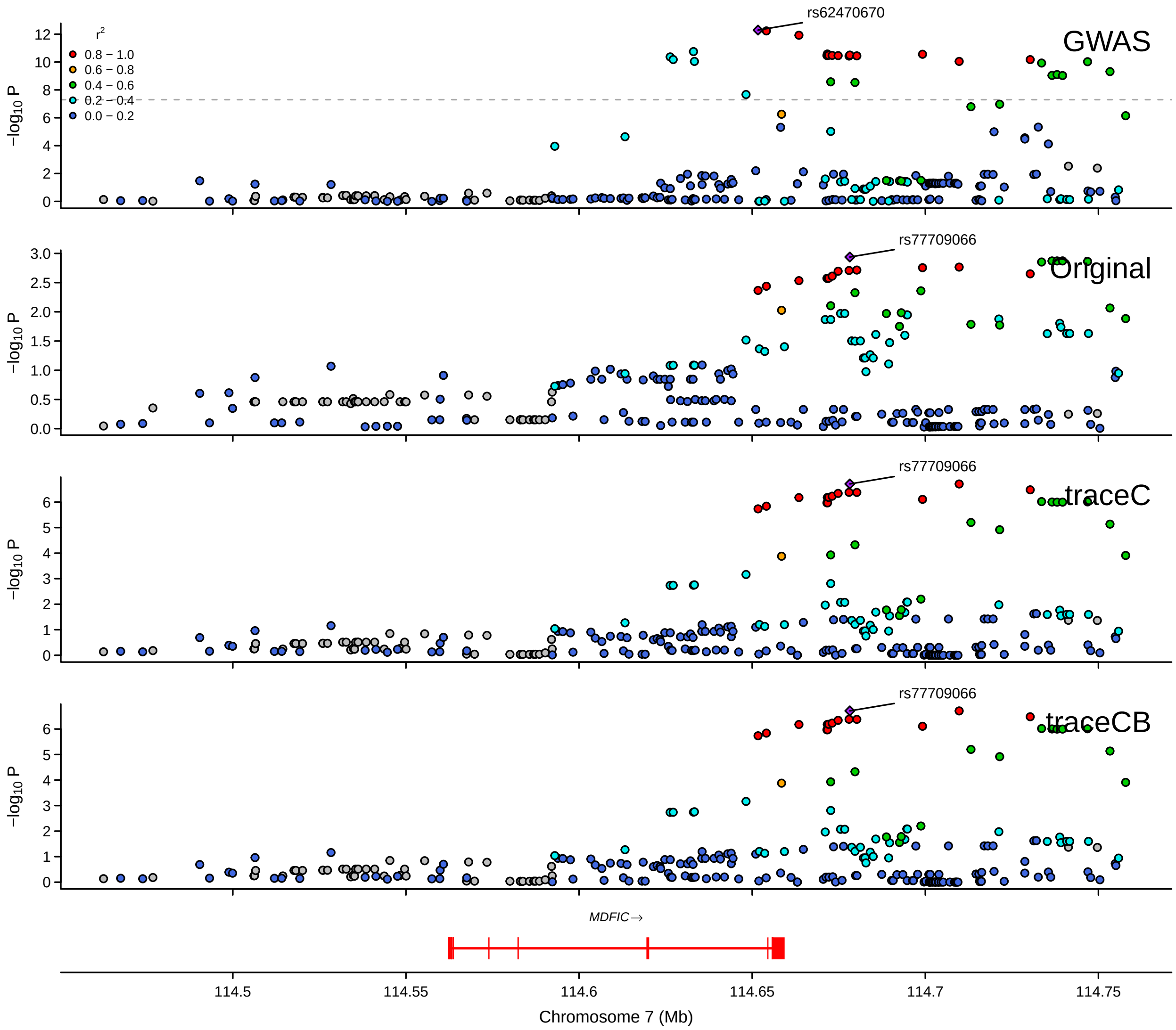

**Supplementary Figure 25:** LocusZoom plot for *MDFIC* in prioritizing BBJ monocyte eQTL with Fairfax_2014(420) and eQTLGen. GWAS summary statistics are from BCX monocyte count.

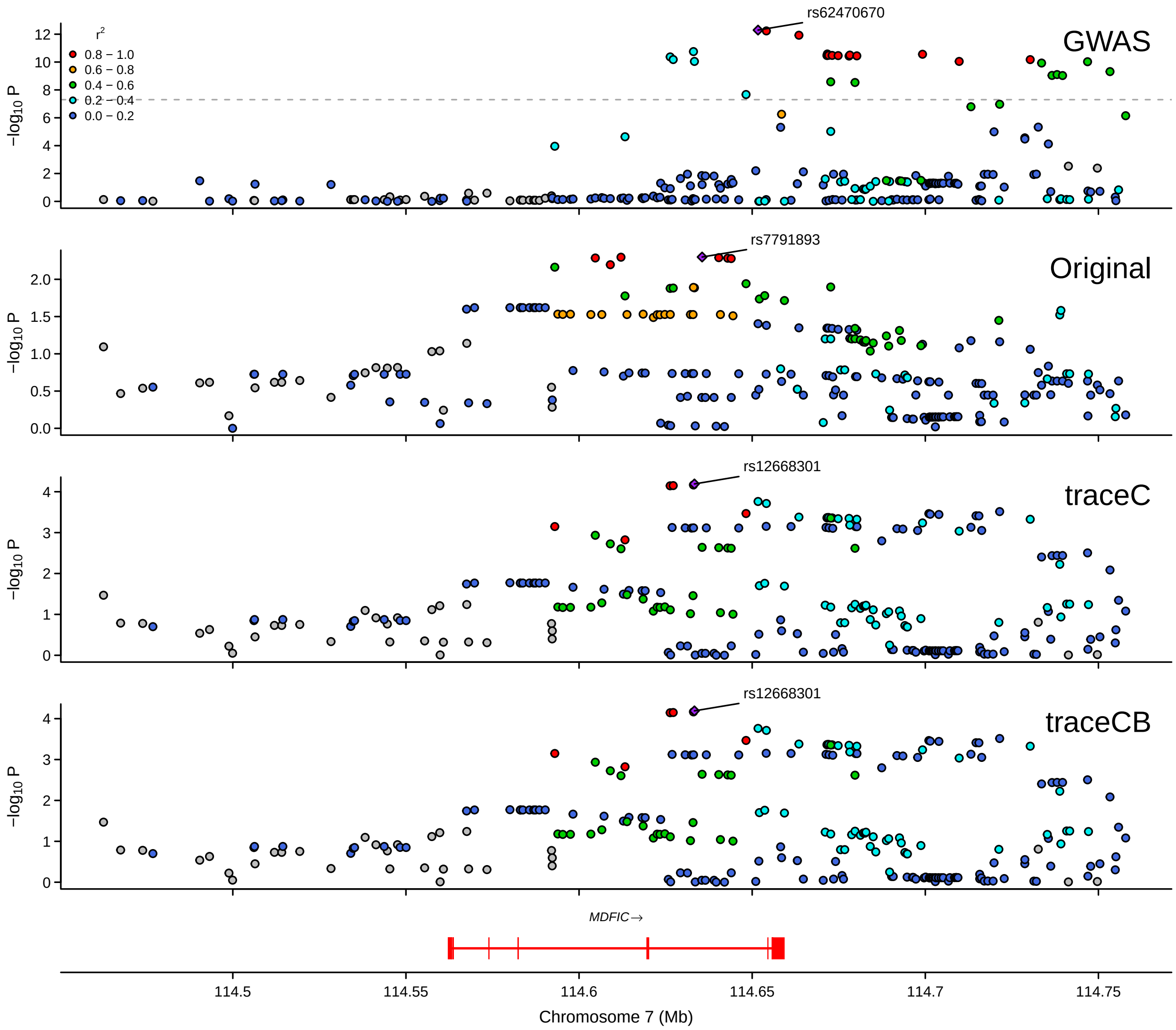

**Supplementary Figure 26:** LocusZoom plot for *MDFIC* in prioritizing BBJ CD4$^+$ T cell eQTL with BLUEPRINT (167) and eQTLGen. GWAS summary statistics are from BCX monocyte count.

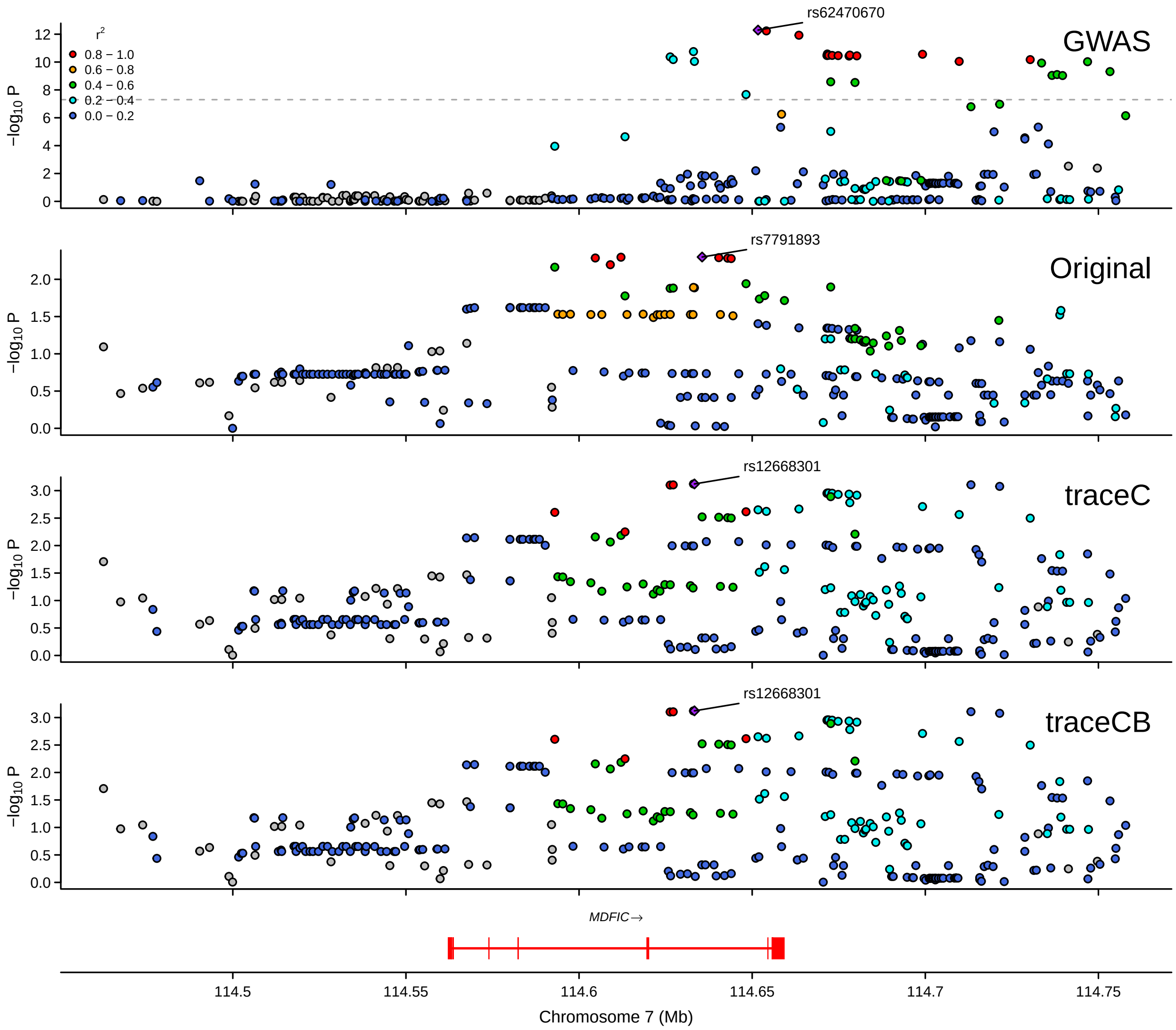

**Supplementary Figure 27:** LocusZoom plot for *MDFIC* in prioritizing BBJ CD4$^+$ T cell eQTL with CEDAR (290) and eQTLGen. GWAS summary statistics are from BCX monocyte count.

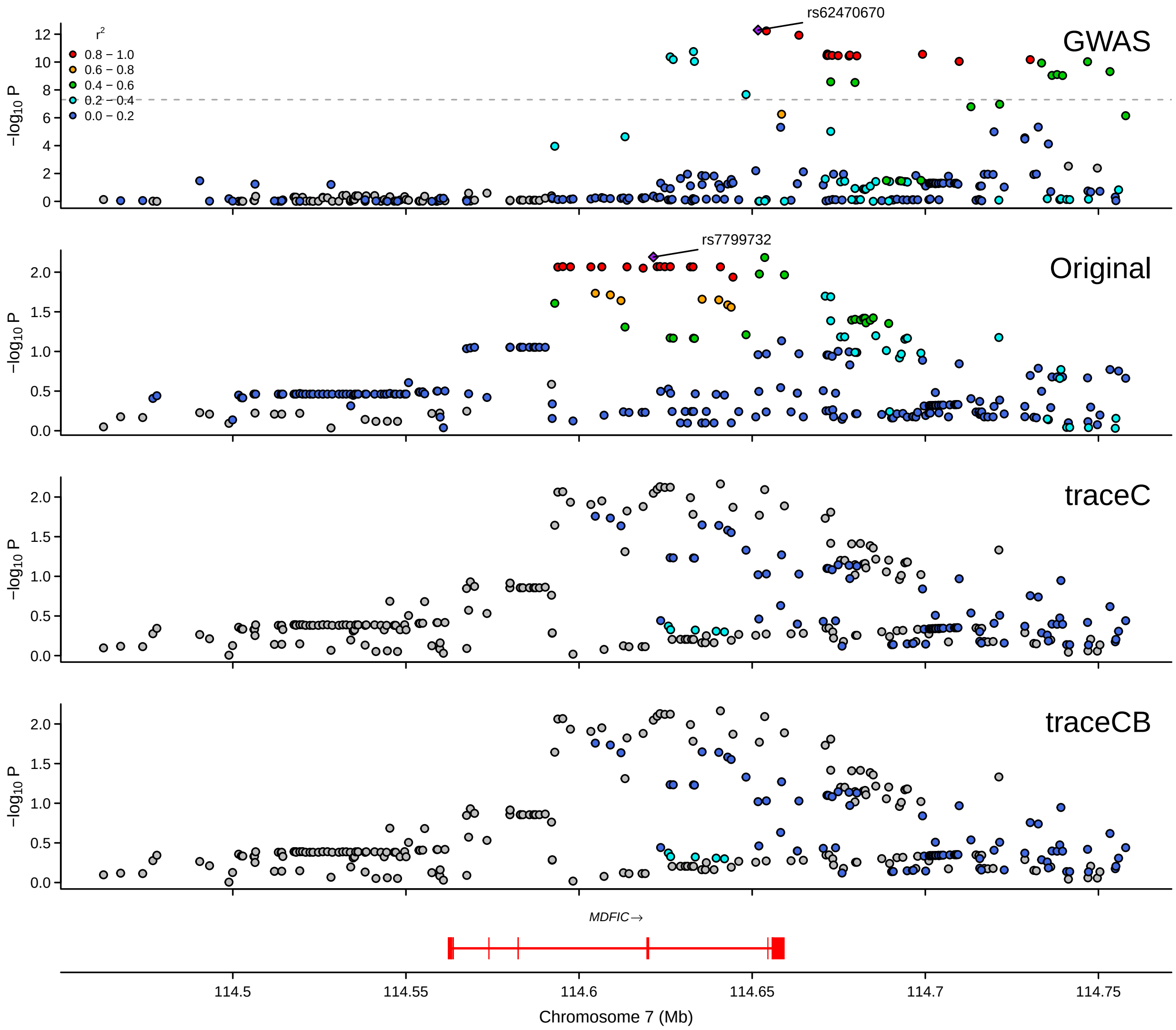

**Supplementary Figure 28:** LocusZoom plot for *MDFIC* in prioritizing BBJ CD8$^{+}$ T cell eQTL with CEDAR (277) and eQTLGen. GWAS summary statistics are from BCX monocyte count.

#### 1.4 Additional results of trans-ancestry eQTL mapping in BBJ cohort

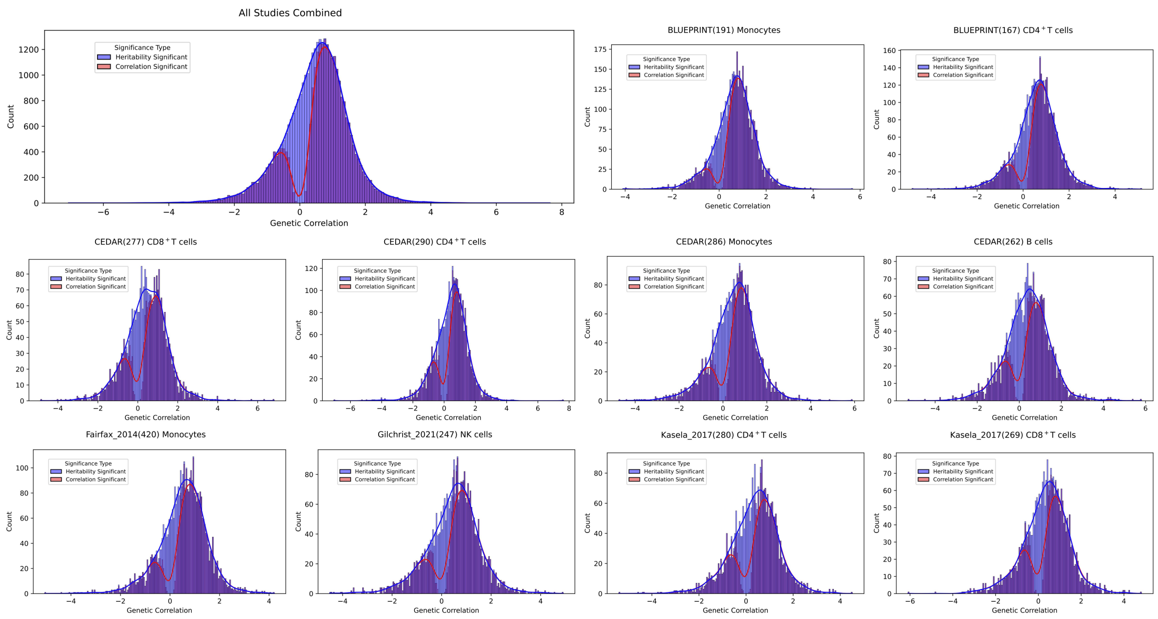

**Supplementary Figure 29:** The distribution of genetic correlation estimates between BBJ ct-eQTL and 10 European ancestry ct-eQTL studies of matched cell types. The density plots are colored by whether the gene has significant correlation between ancestries.

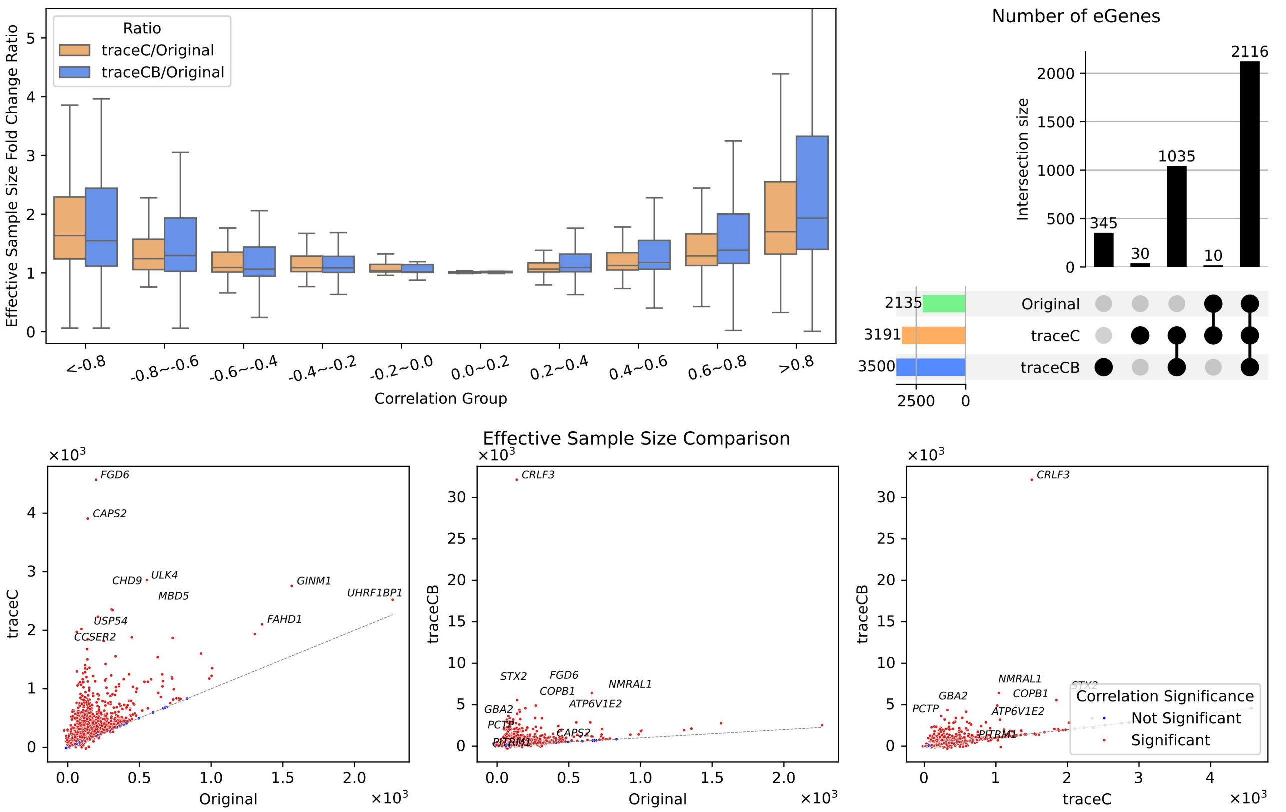

**Supplementary Figure 30:** Case study of BBJ monocyte eQTL prioritization using BLUEPRINT (191) and eQTLGen. Upper left: Effective sample size ratio compared across genetic correlation quartiles; Upper right: Upset plots illustrate the number of eGenes identified by each analysis, and the novel eGenes discovered by traceC and traceCB. Lower: Effective sample size comparison restricted to genes with significant heritability across both ancestries. Colors represent genes with (red) and without (blue) significant trans-ancestry correlation.

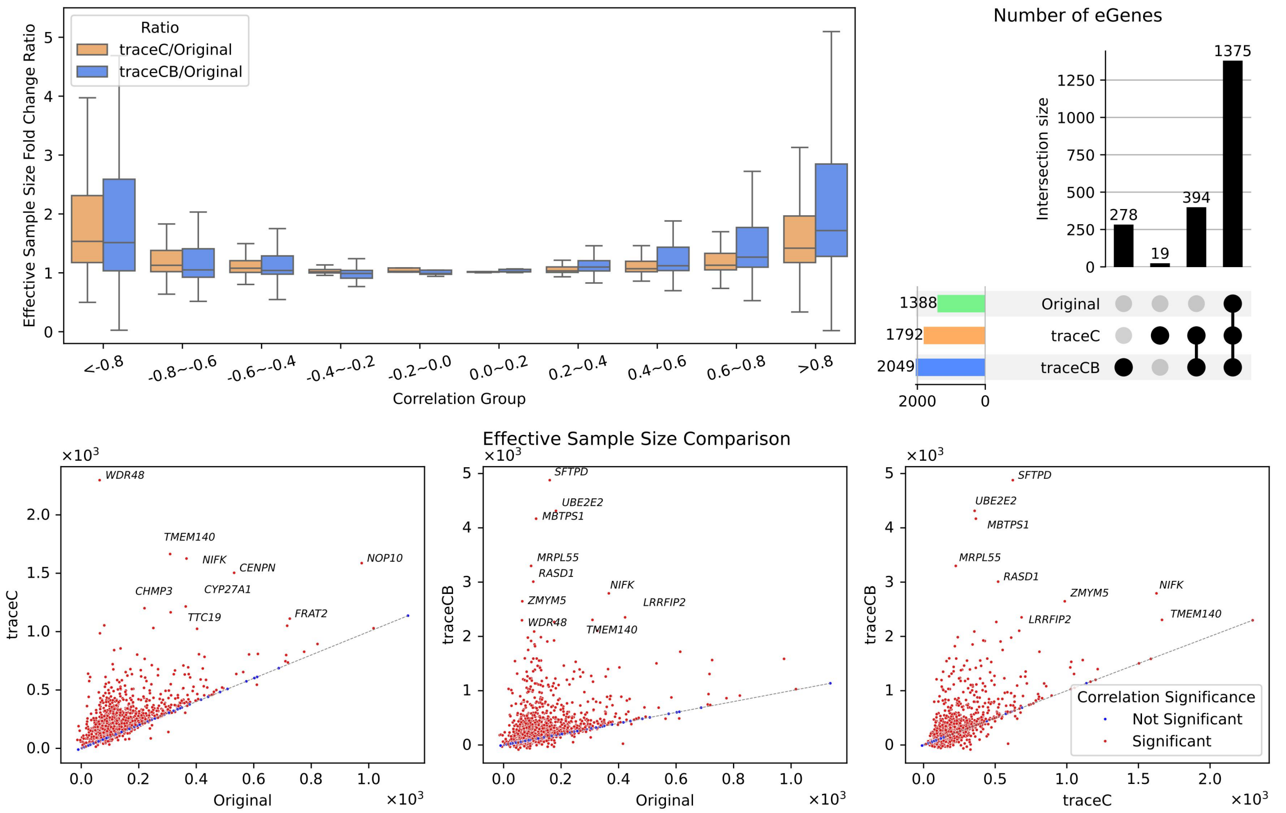

**Supplementary Figure 31:** Case study of BBJ monocyte eQTL prioritization using CEDAR (286) and eQTLGen. Upper left: Effective sample size ratio comparison across genetic correlation quartiles; Upper right: Upset plots illustrate the number of eGenes identified by each analysis, and the novel eGenes discovered by traceC and traceCB. Lower: Effective sample size comparison restricted to genes with significant heritability across both ancestries. Colors represent genes with (red) and without (blue) significant trans-ancestry correlation.

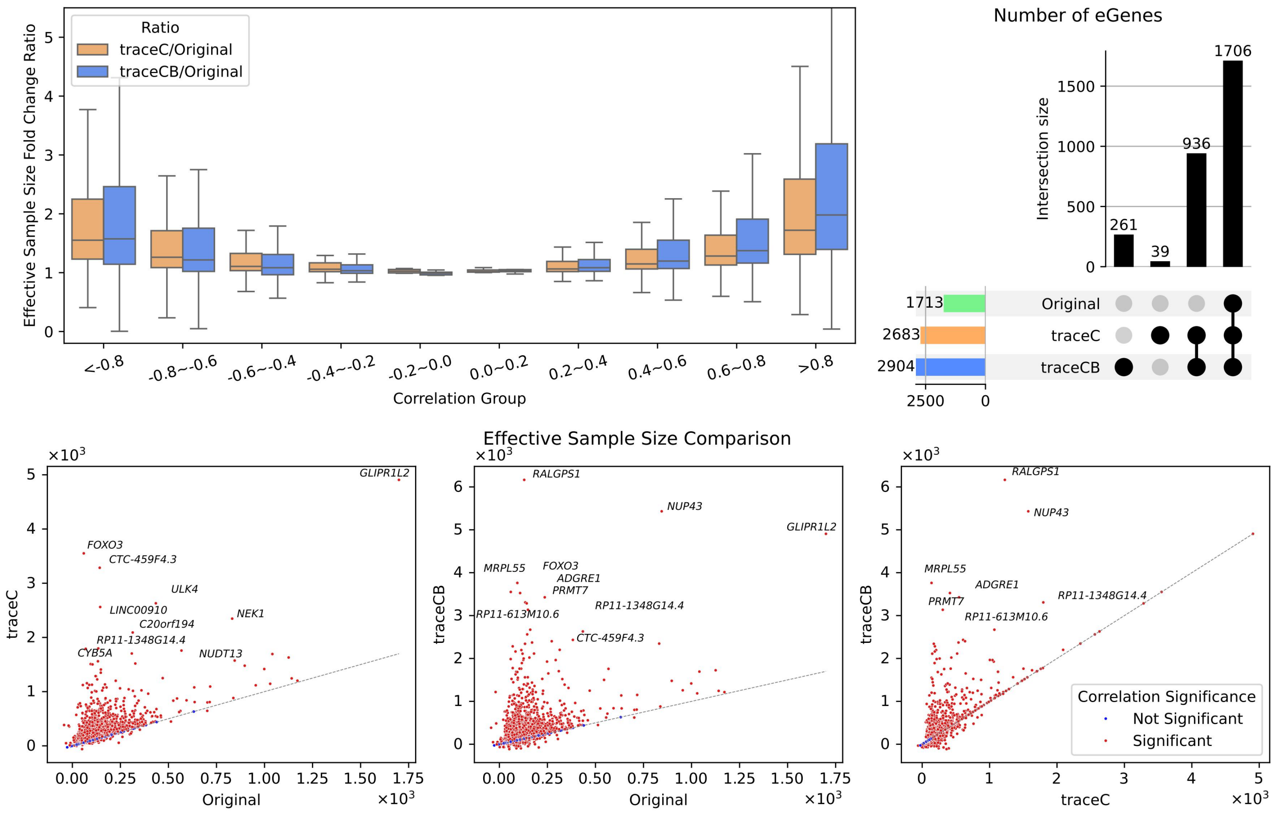

**Supplementary Figure 32:** Case study of BBJ CD4$^+$ T eQTL prioritization using BLUEPRINT (167) and eQTLGen. Upper left: Effective sample size ratio comparison across genetic correlation quartiles; Upper right: Upset plots illustrate the number of eGenes identified by each analysis, and the novel eGenes discovered by traceC and traceCB. Lower: Effective sample size comparison restricted to genes with significant heritability across both ancestries. Colors represent genes with (red) and without (blue) significant trans-ancestry correlation.

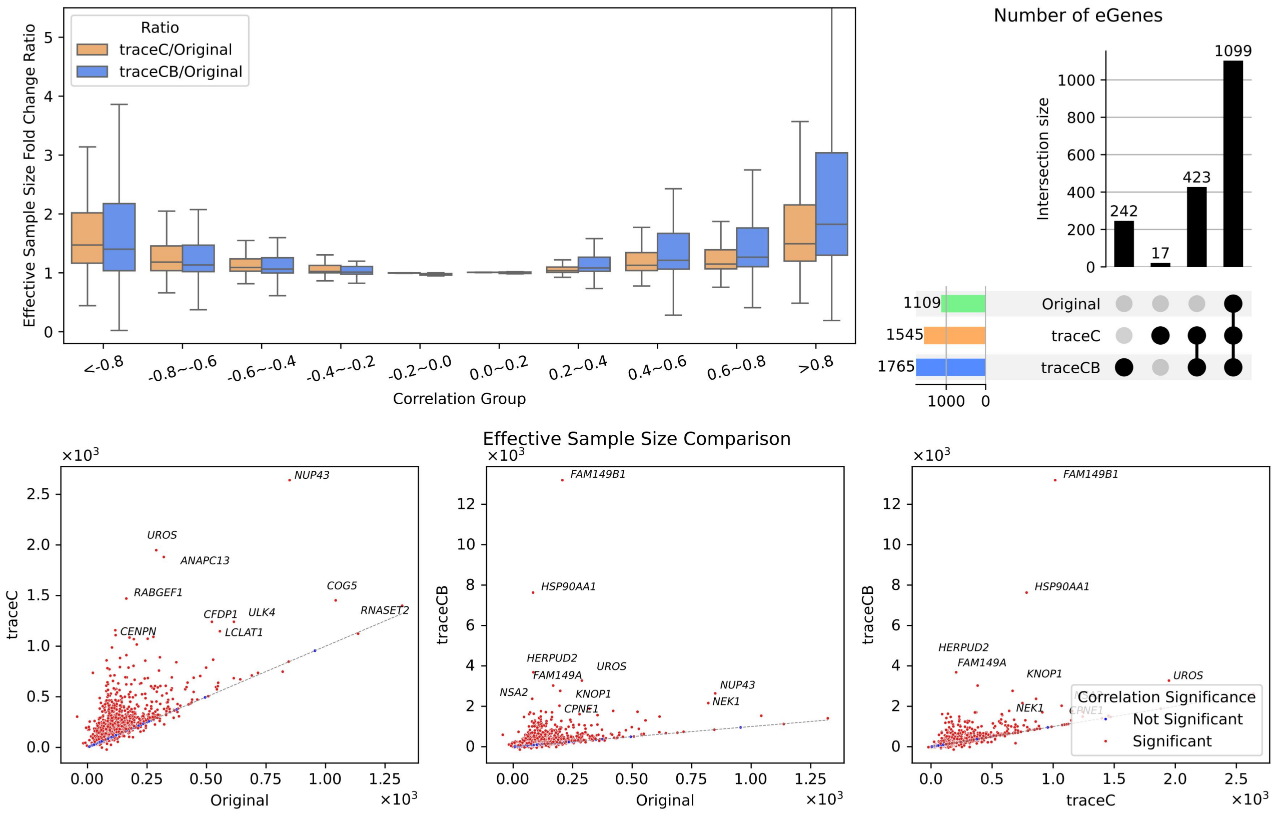

**Supplementary Figure 33:** Case study of BBJ CD4$^{+}$ T eQTL prioritization using CEDAR (290) and eQTLGen. Upper left: Effective sample size ratio comparison across genetic correlation quartiles; Upper right: Upset plots illustrate the number of eGenes identified by each analysis, and the novel eGenes discovered by traceC and traceCB. Lower: Effective sample size comparison restricted to genes with significant heritability across both ancestries. Colors represent genes with (red) and without (blue) significant trans-ancestry correlation.

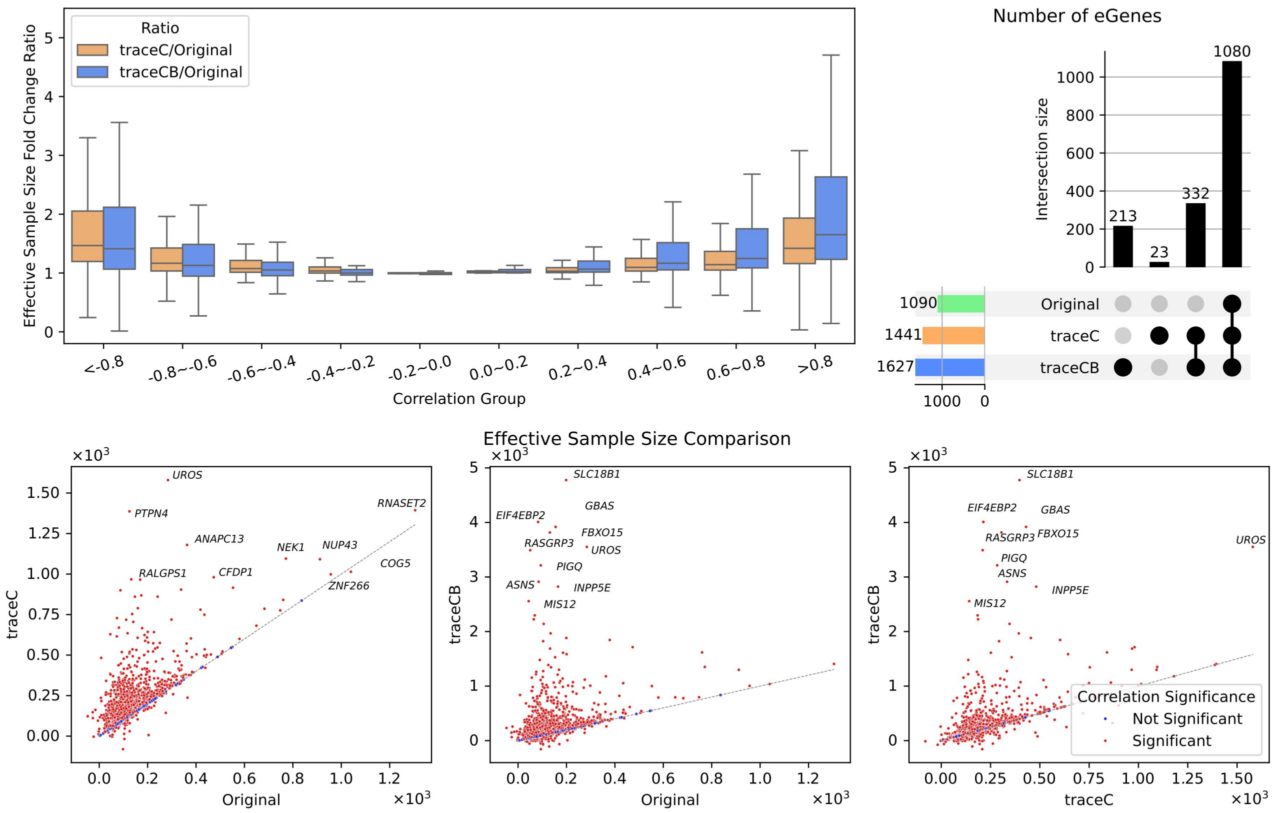

**Supplementary Figure 34:** Case study of BBJ CD4$^+$ T eQTL prioritization using Kasela_2017 (280) and eQTLGen. Upper left: Effective sample size ratio comparison across genetic correlation quartiles; Upper right: Upset plots illustrate the number of eGenes identified by each analysis, and the novel eGenes discovered by traceC and traceCB. Lower: Effective sample size comparison restricted to genes with significant heritability across both ancestries. Colors represent genes with (red) and without (blue) significant trans-ancestry correlation.

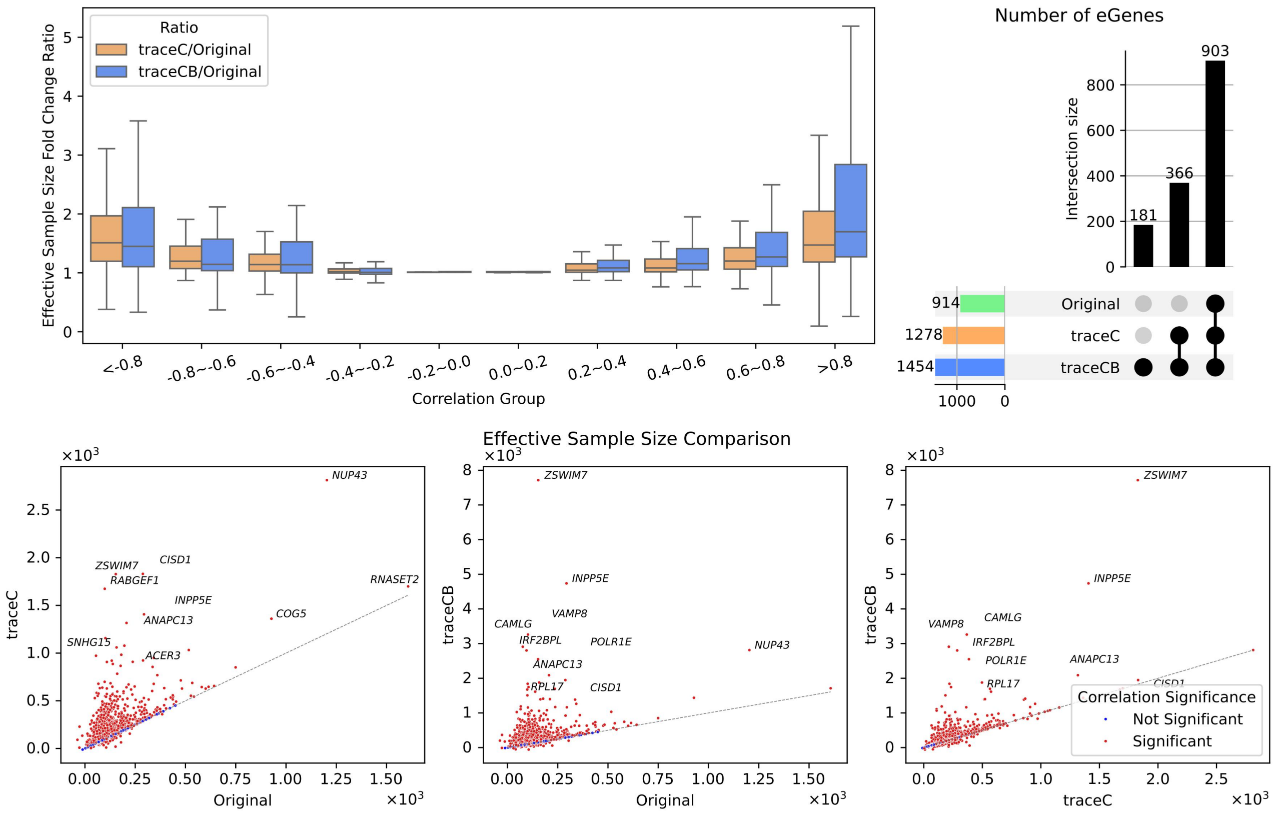

**Supplementary Figure 35:** Case study of BBJ CD8$^{+}$ T eQTL prioritization using CEDAR (277) and eQTLGen. Upper left: Effective sample size ratio comparison across genetic correlation quartiles; Upper right: Upset plots illustrate the number of eGenes identified by each analysis, and the novel eGenes discovered by traceC and traceCB. Lower: Effective sample size comparison restricted to genes with significant heritability across both ancestries. Colors represent genes with (red) and without (blue) significant trans-ancestry correlation.

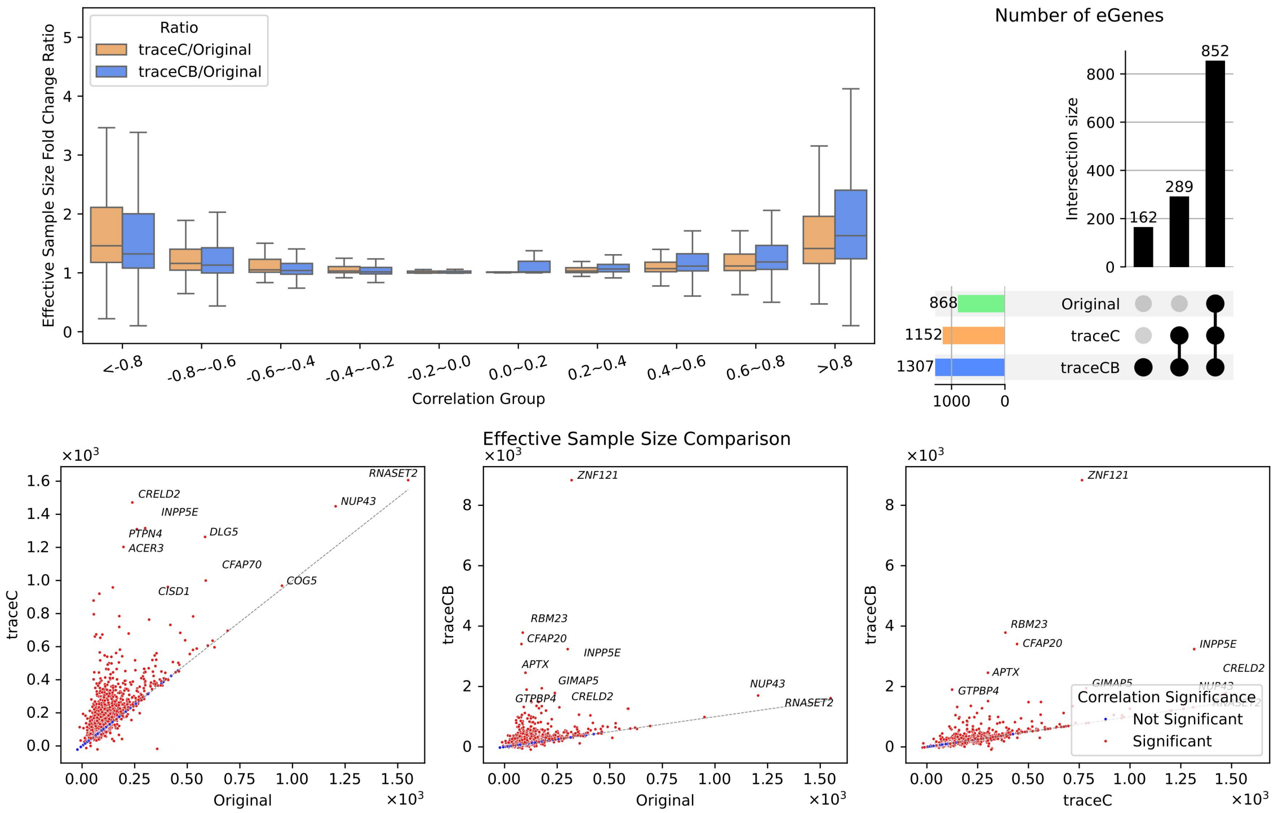

**Supplementary Figure 36:** Case study of BBJ CD8$^+$ T eQTL prioritization using Kasela_2017 (269) and eQTLGen. Upper left: Effective sample size ratio comparison across genetic correlation quartiles; Upper right: Upset plots illustrate the number of eGenes identified by each analysis, and the novel eGenes discovered by traceC and traceCB. Lower: Effective sample size comparison restricted to genes with significant heritability across both ancestries. Colors represent genes with (red) and without (blue) significant trans-ancestry correlation.

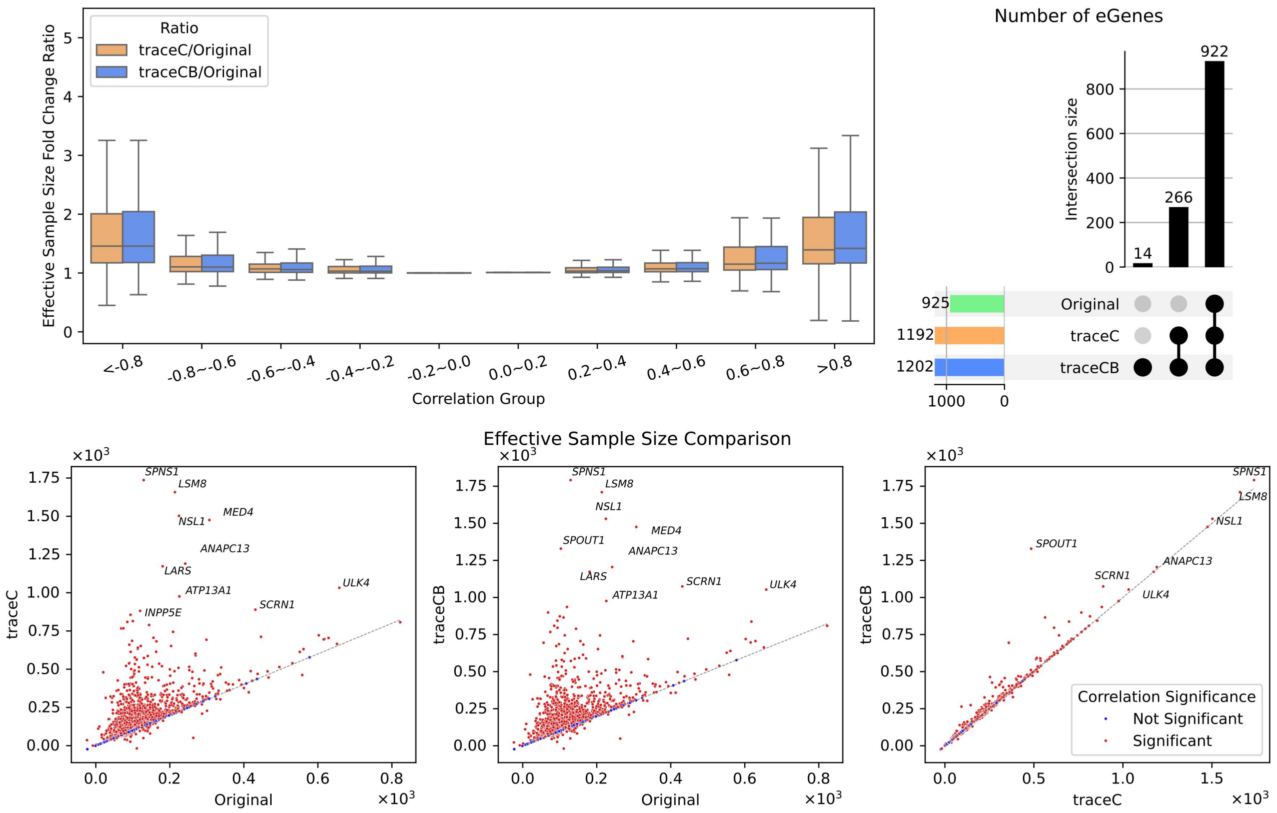

**Supplementary Figure 37:** Case study of BBJ B cell eQTL prioritization using CEDAR (262) and eQTLGen. Upper left: Effective sample size ratio comparison across genetic correlation quartiles; Upper right: Upset plots illustrate the number of eGenes identified by each analysis, and the novel eGenes discovered by traceC and traceCB. Lower: Effective sample size comparison restricted to genes with significant heritability across both ancestries. Colors represent genes with (red) and without (blue) significant trans-ancestry correlation.

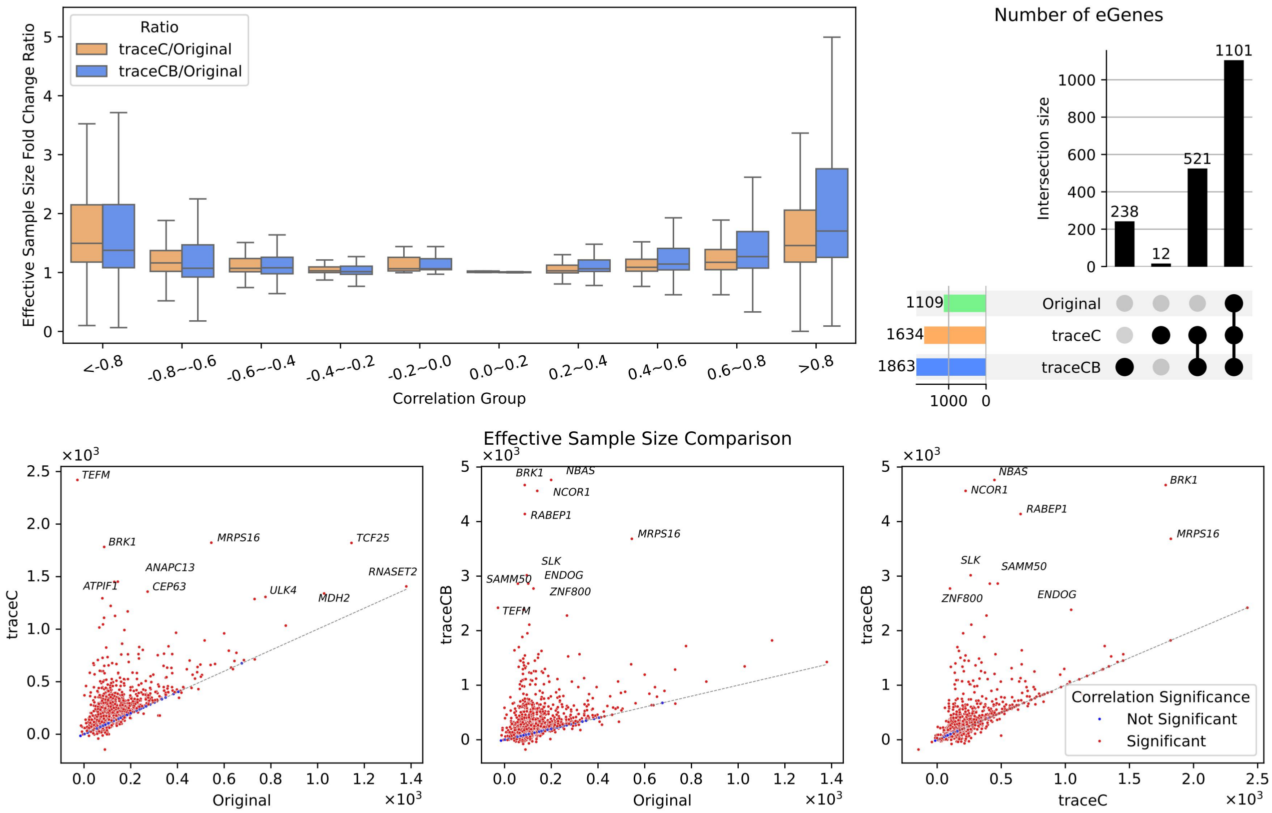

**Supplementary Figure 38:** Case study of BBJ NK cell eQTL prioritization using Gilchrist_2021 (247) and eQTLGen. Upper left: Effective sample size ratio comparison across genetic correlation quartiles; Upper right: Upset plots illustrate the number of eGenes identified by each analysis, and the novel eGenes discovered by traceC and traceCB. Lower: Effective sample size comparison restricted to genes with significant heritability across both ancestries. Colors represent genes with (red) and without (blue) significant trans-ancestry correlation.

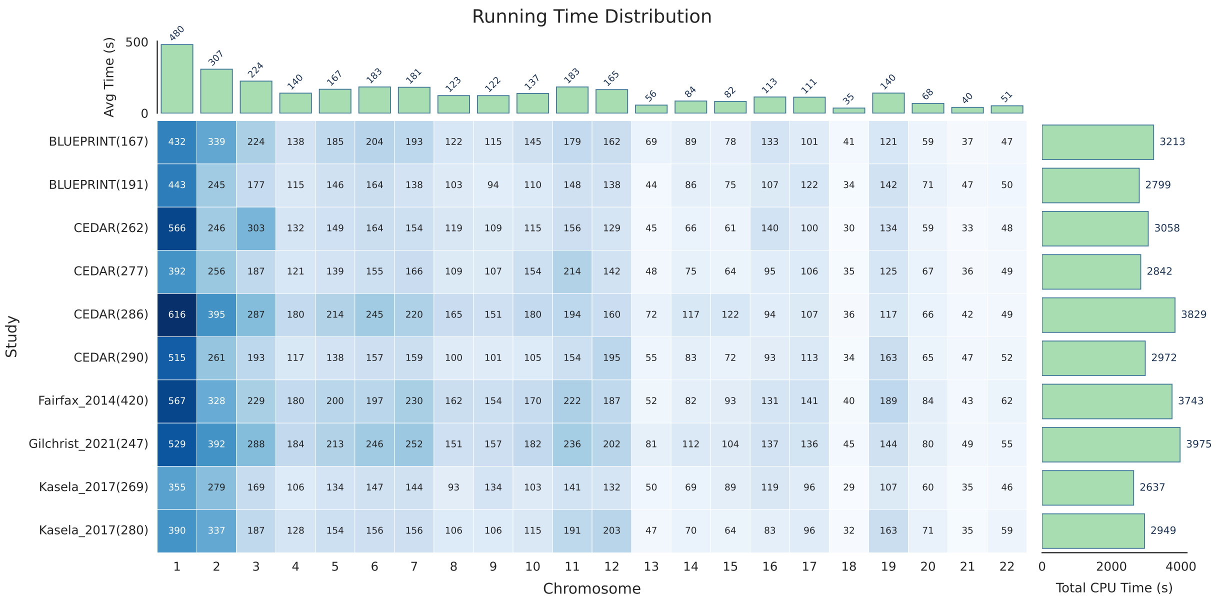

**Supplementary Figure 39:** Computational efficiency of traceCB across chromosomes and studies. The heatmap displays the runtime (in seconds) for traceCB applied to BBJ data integrated with eQTL Catalogue and eQTLGen datasets. The top marginal bar plot shows the mean runtime per chromosome averaged across all studies, while the right marginal bar plot indicates the total cpu time per study summed across all chromosomes.

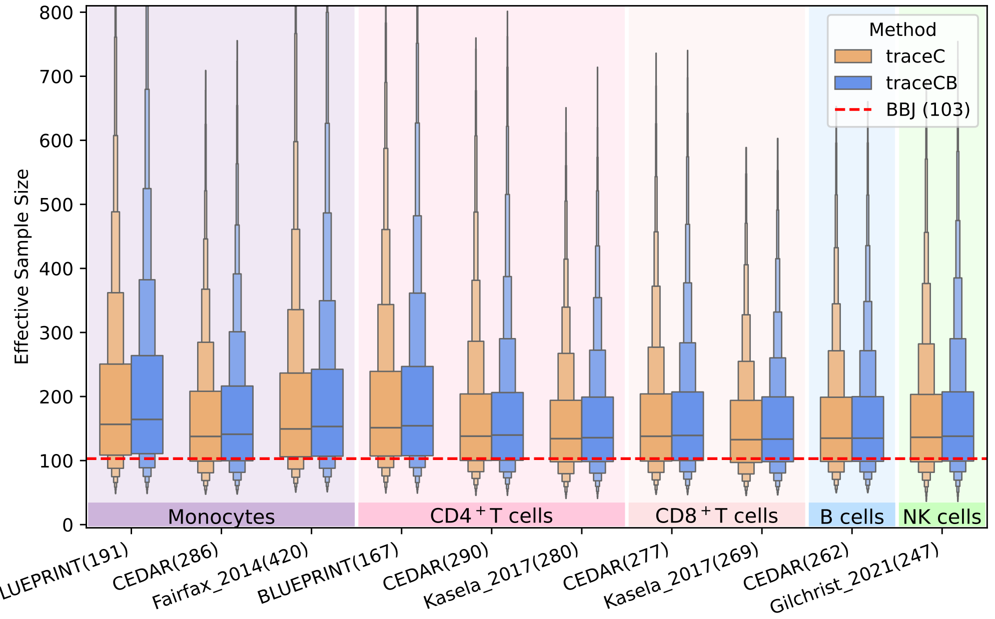

**Supplementary Figure 40:** Distribution of effective sample size for ct-eQTLs prioritized by traceC (orange) and traceCB (blue) relative to the original BBJ eQTL study sample size (red dashed line, sample size = 103), where the tissue-level eQTLs comes from GTEx.

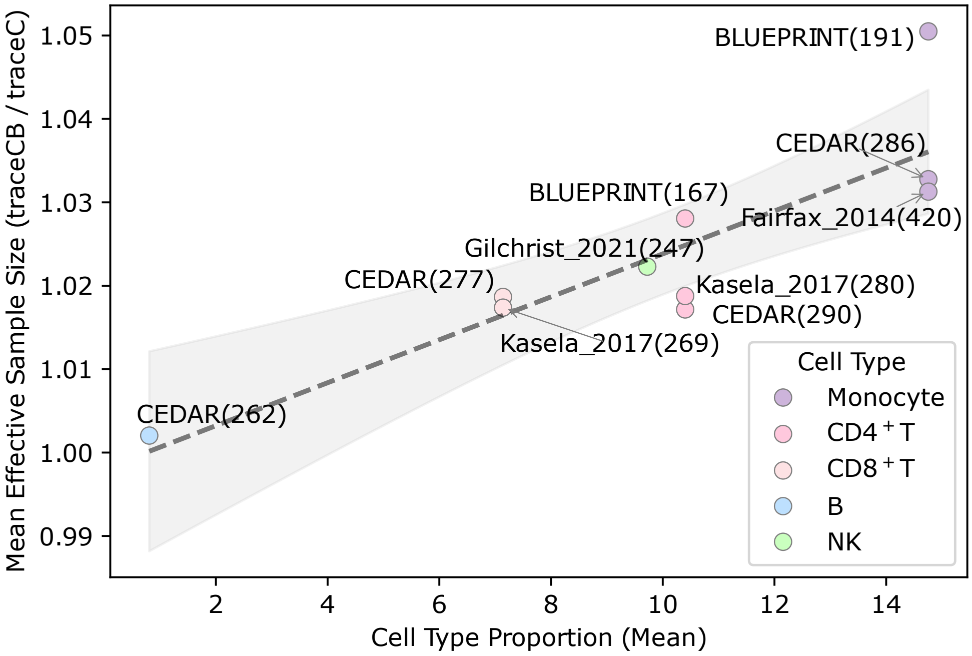

**Supplementary Figure 41:** Relationship between the cell-type proportions in GTEx whole blood and the effective sample size ratio (traceCB vs. traceC) across different cell types and studies. Regression lines with 95% confidence intervals are shown in gray.

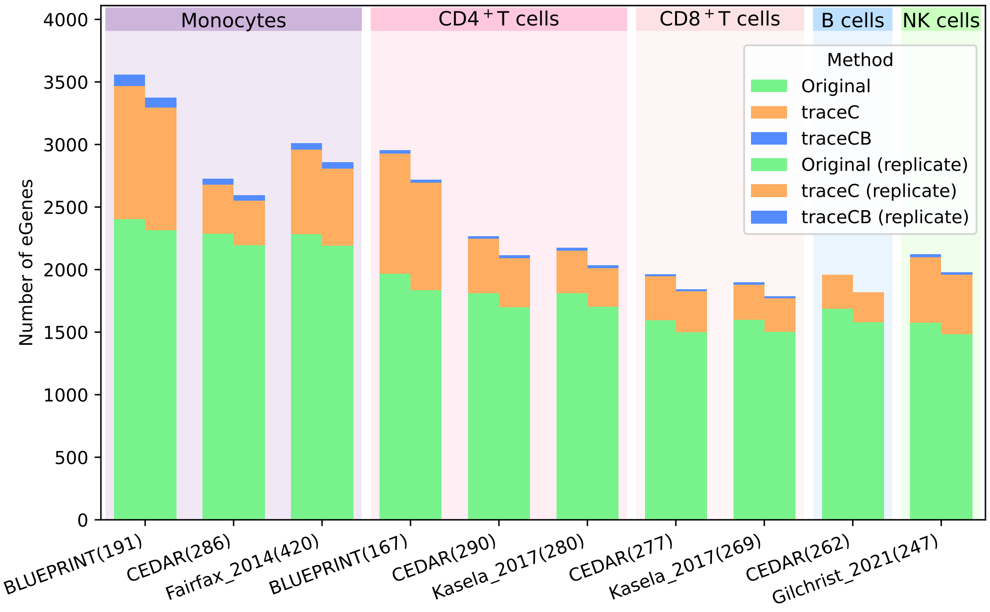

**Supplementary Figure 42:** Number of eGenes identified by the original BBJ analysis, and the novel eGenes discovered by traceC and traceCB when integrating with GTEx bulk eQTLs across different cell types and studies.

#### 1.5 Additional results of trans-ancestry eQTL mapping in African cohort

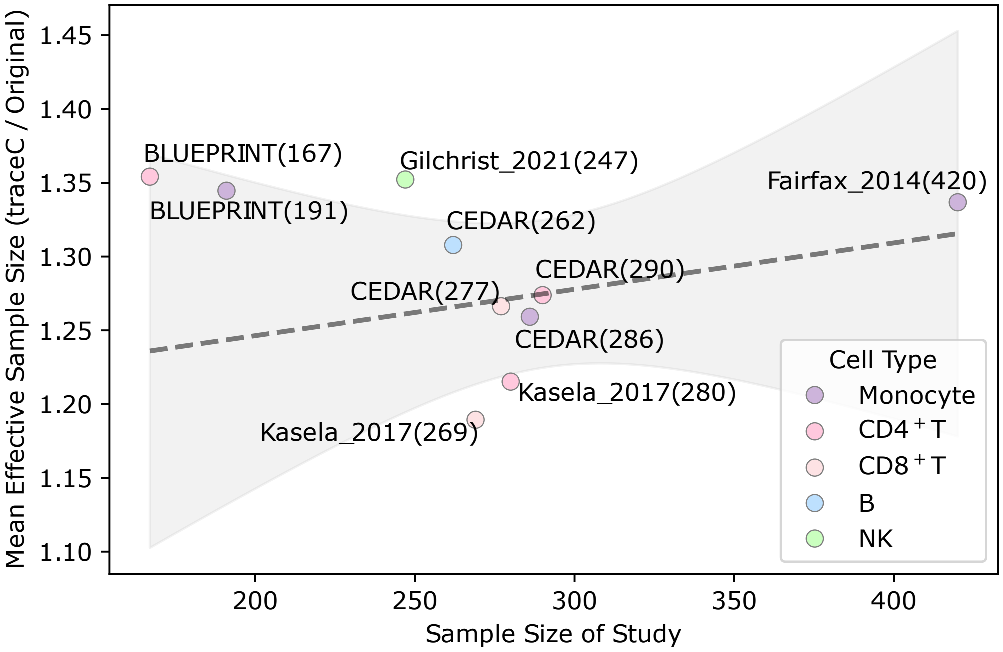

**Supplementary Figure 43:** Relationship between the auxiliary European ancestry ct-eQTL study sample size and the effective sample size ratio (traceC vs. African eQTL study) across different cell types and studies. Regression lines with 95% confidence intervals are shown in gray.

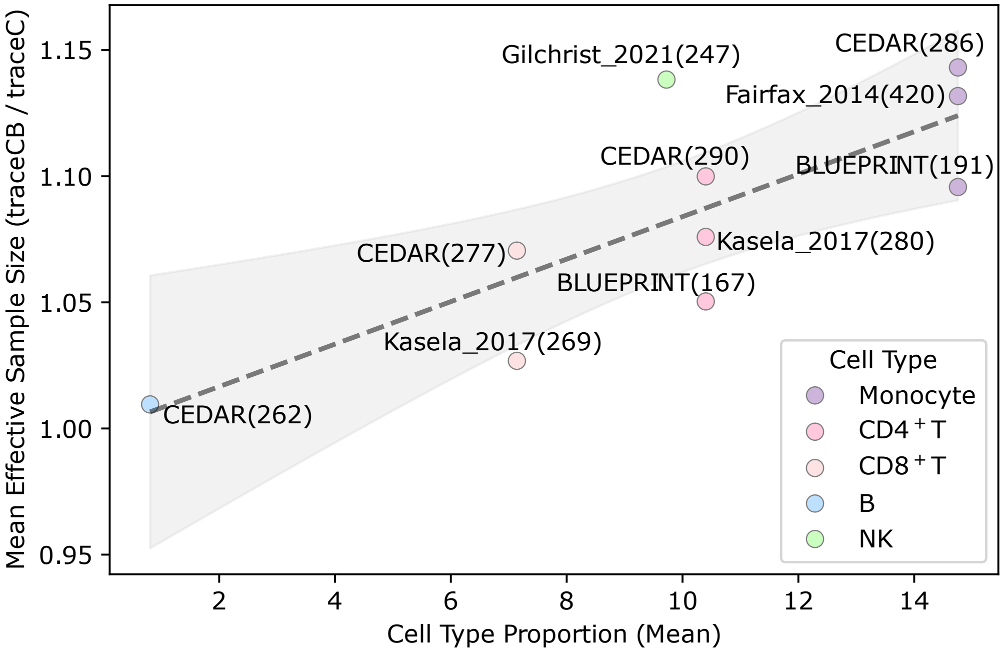

**Supplementary Figure 44:** Relationship between the cell type proportion in GTEx whole blood and the effective sample size ratio (traceCB vs. traceC) across different cell types and studies. Regression lines with 95% confidence intervals are shown in gray.

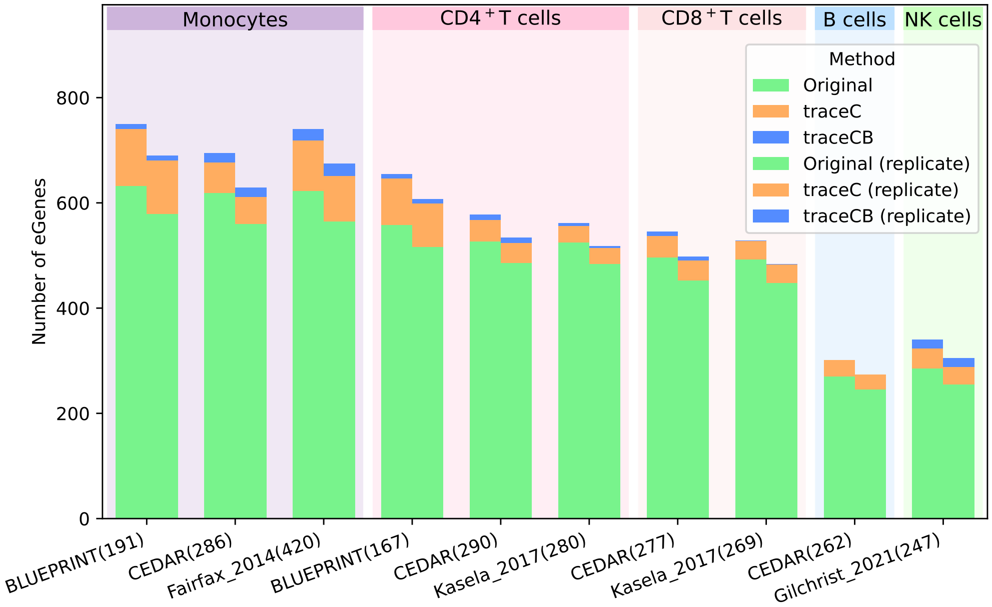

**Supplementary Figure 45:** Number of eGenes identified by the original African analysis, and the novel eGenes discovered by traceC and traceCB when integrating with eQTLGen bulk eQTLs across different cell types and studies.

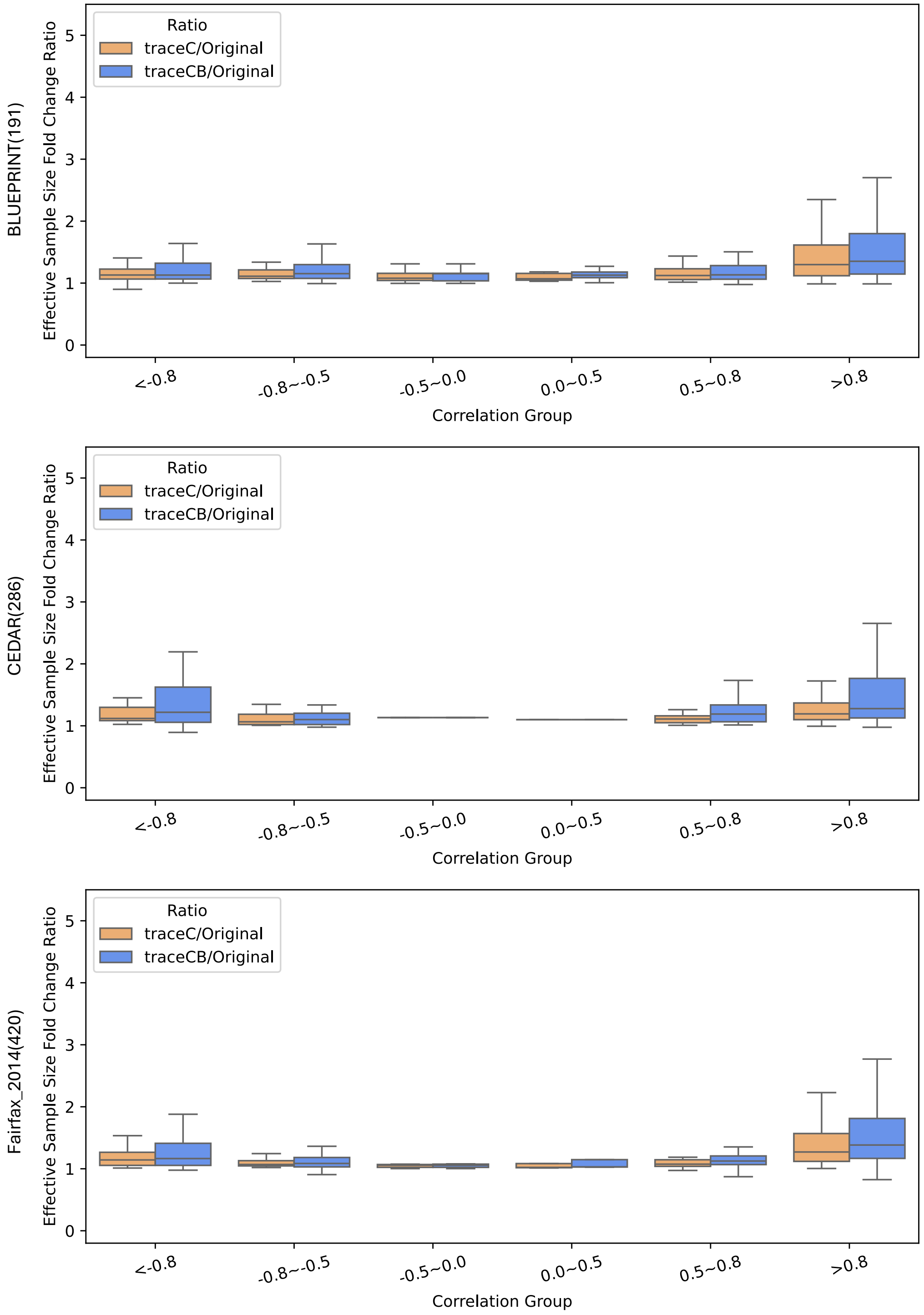

**Supplementary Figure 46:** Effective sample size ratio comparison across genetic correlation quartiles in three African monocyte eQTL studies when integrating with eQTLGen bulk eQTLs.

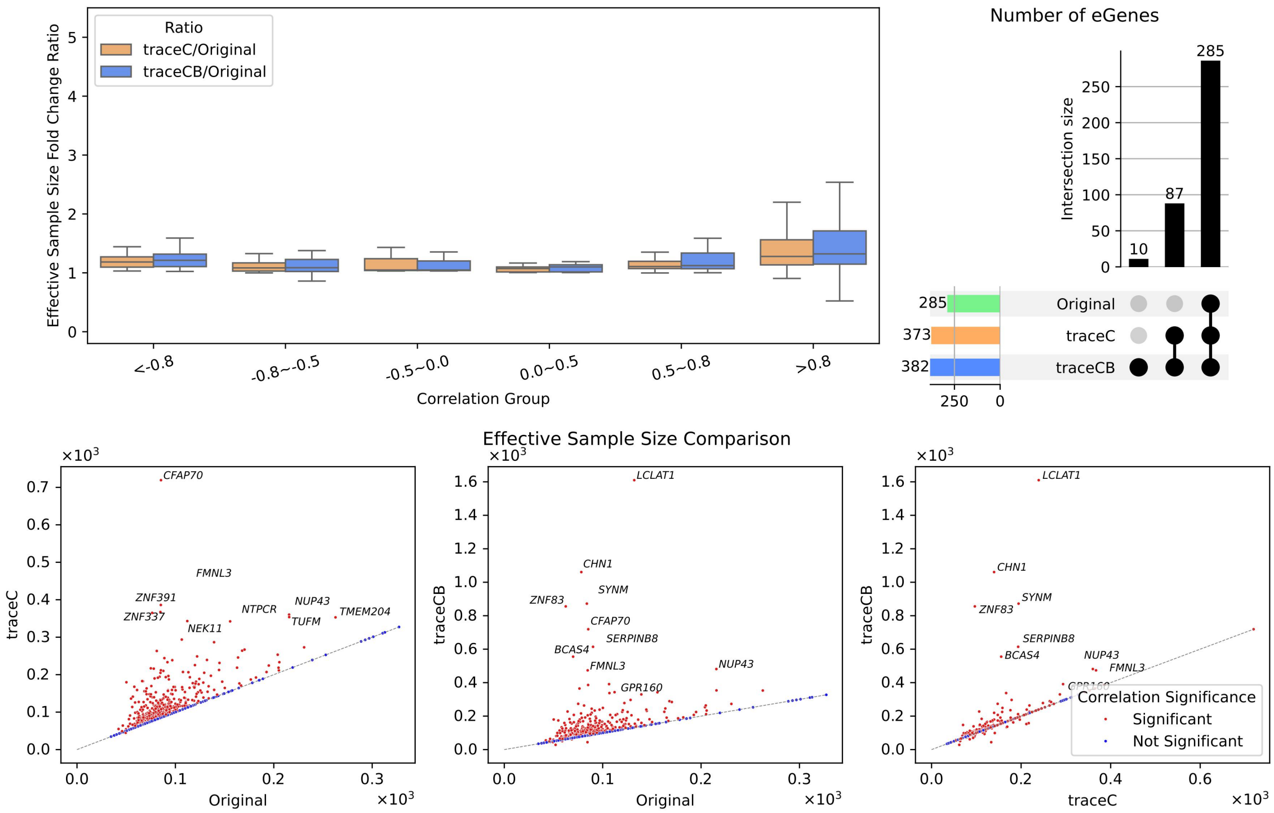

**Supplementary Figure 47:** Case study of African CD4$^+$ T eQTL prioritization using BLUEPRINT (167) and eQTLGen. Upper left: Effective sample size ratio comparison across genetic correlation quartiles; Upper right: Upset plots illustrate the number of eGenes identified by each analysis, and the novel eGenes discovered by traceC and traceCB. Lower: Effective sample size comparison restricted to genes with significant heritability across both ancestries. Colors represent genes with (red) and without (blue) significant trans-ancestry correlation.

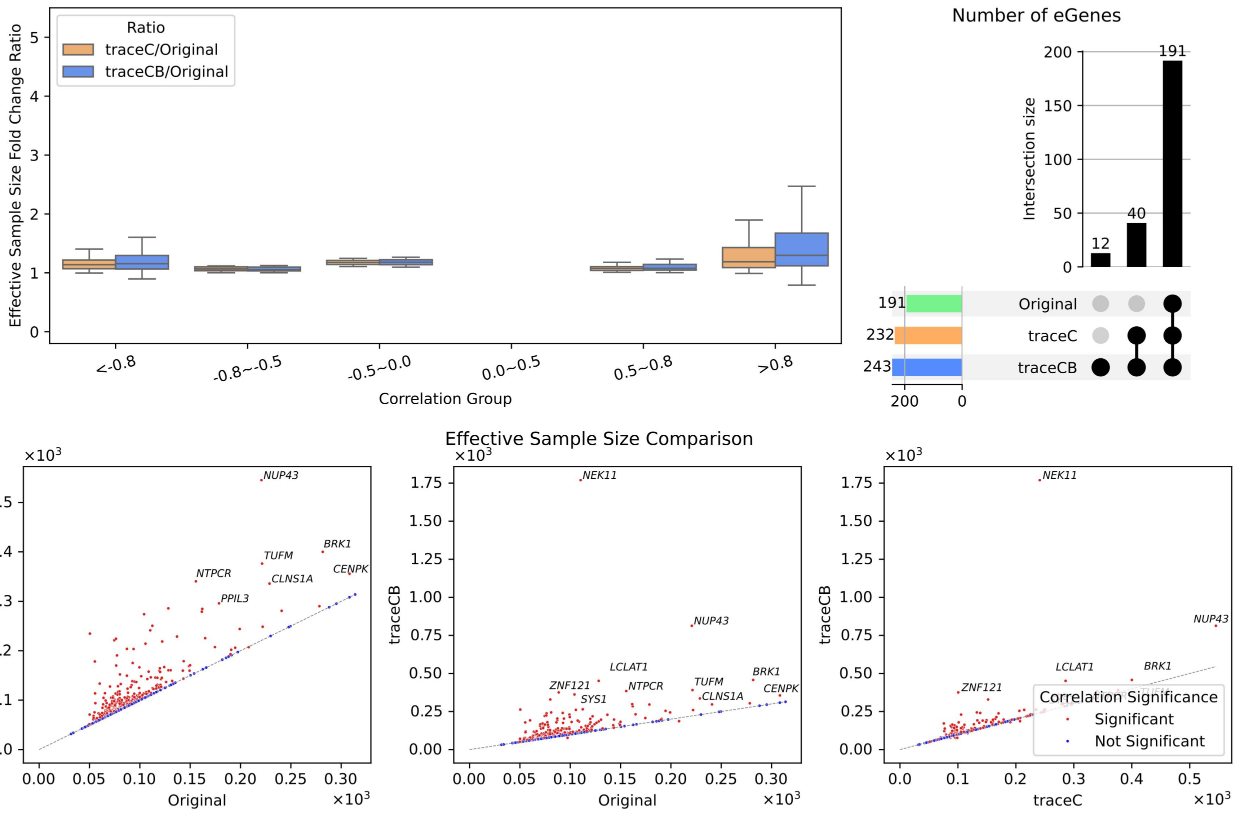

**Supplementary Figure 48:** Case study of African CD4$^+$ T eQTL prioritization using CEDAR (290) and eQTLGen. Upper left: Effective sample size ratio comparison across genetic correlation quartiles; Upper right: Upset plots illustrate the number of eGenes identified by each analysis, and the novel eGenes discovered by traceC and traceCB. Lower: Effective sample size comparison restricted to genes with significant heritability across both ancestries. Colors represent genes with (red) and without (blue) significant trans-ancestry correlation.

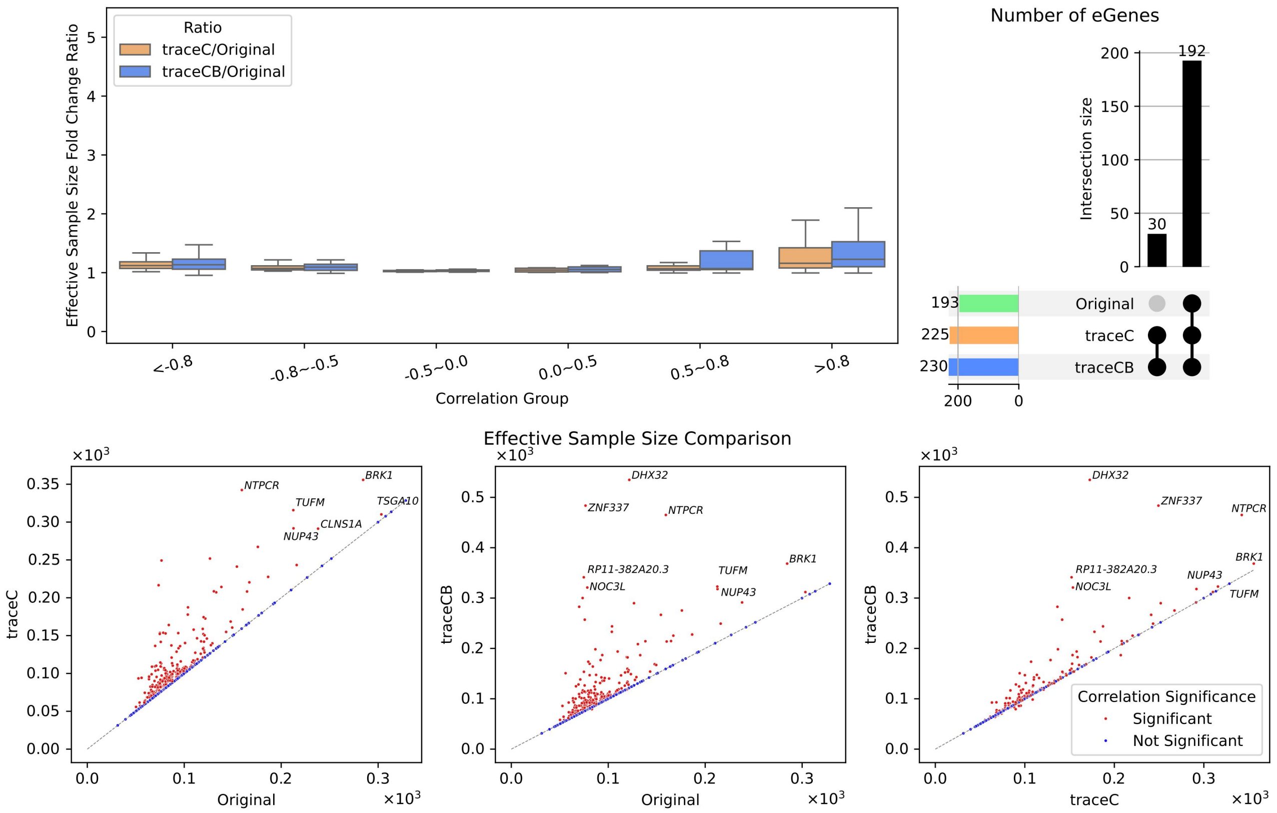

**Supplementary Figure 49:** Case study of African CD4$^+$ T eQTL prioritization using Kasela_2017 (280) and eQTLGen. Upper left: Effective sample size ratio comparison across genetic correlation quartiles; Upper right: Upset plots illustrate the number of eGenes identified by each analysis, and the novel eGenes discovered by traceC and traceCB. Lower: Effective sample size comparison restricted to genes with significant heritability across both ancestries. Colors represent genes with (red) and without (blue) significant trans-ancestry correlation.

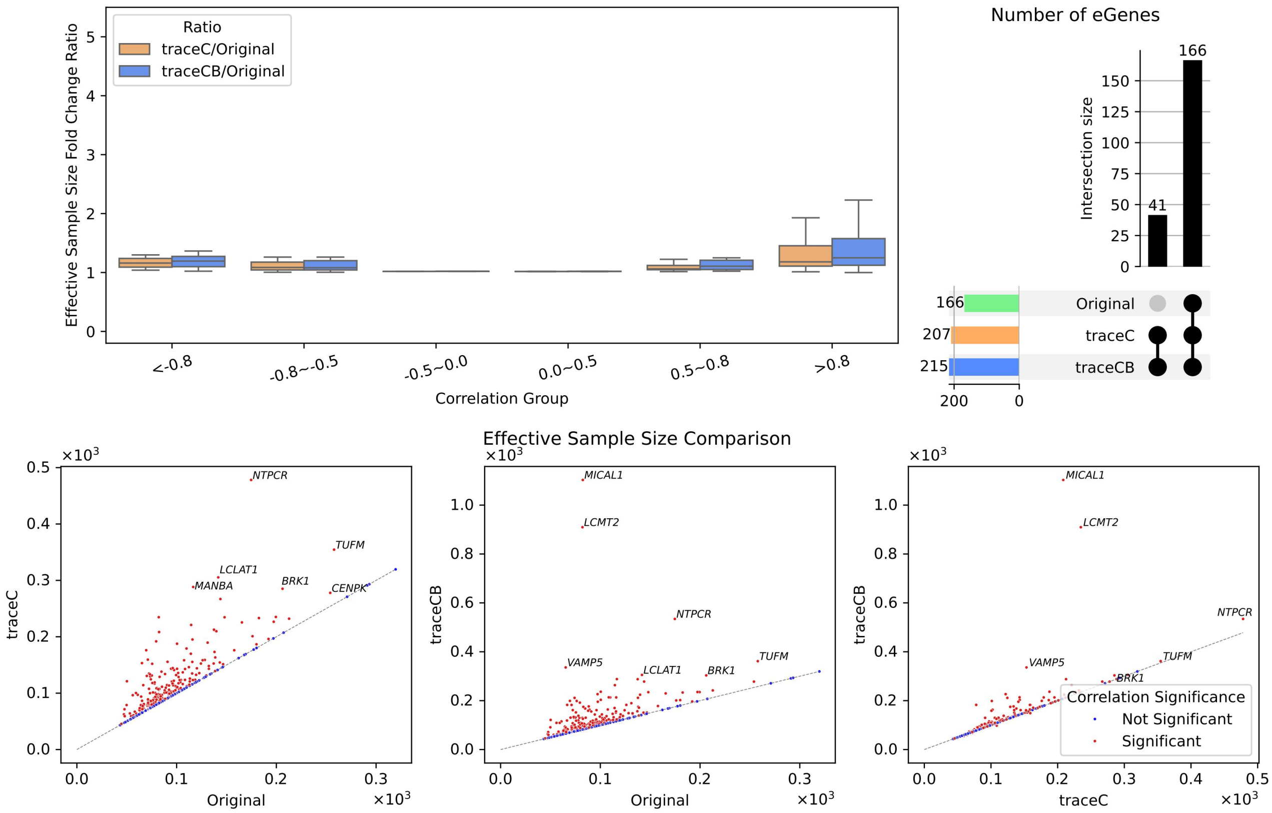

**Supplementary Figure 50:** Case study of African CD8$^{+}$ T eQTL prioritization using CEDAR (277) and eQTLGen. Upper left: Effective sample size ratio comparison across genetic correlation quartiles; Upper right: Upset plots illustrate the number of eGenes identified by each analysis, and the novel eGenes discovered by traceC and traceCB. Lower: Effective sample size comparison restricted to genes with significant heritability across both ancestries. Colors represent genes with (red) and without (blue) significant trans-ancestry correlation.

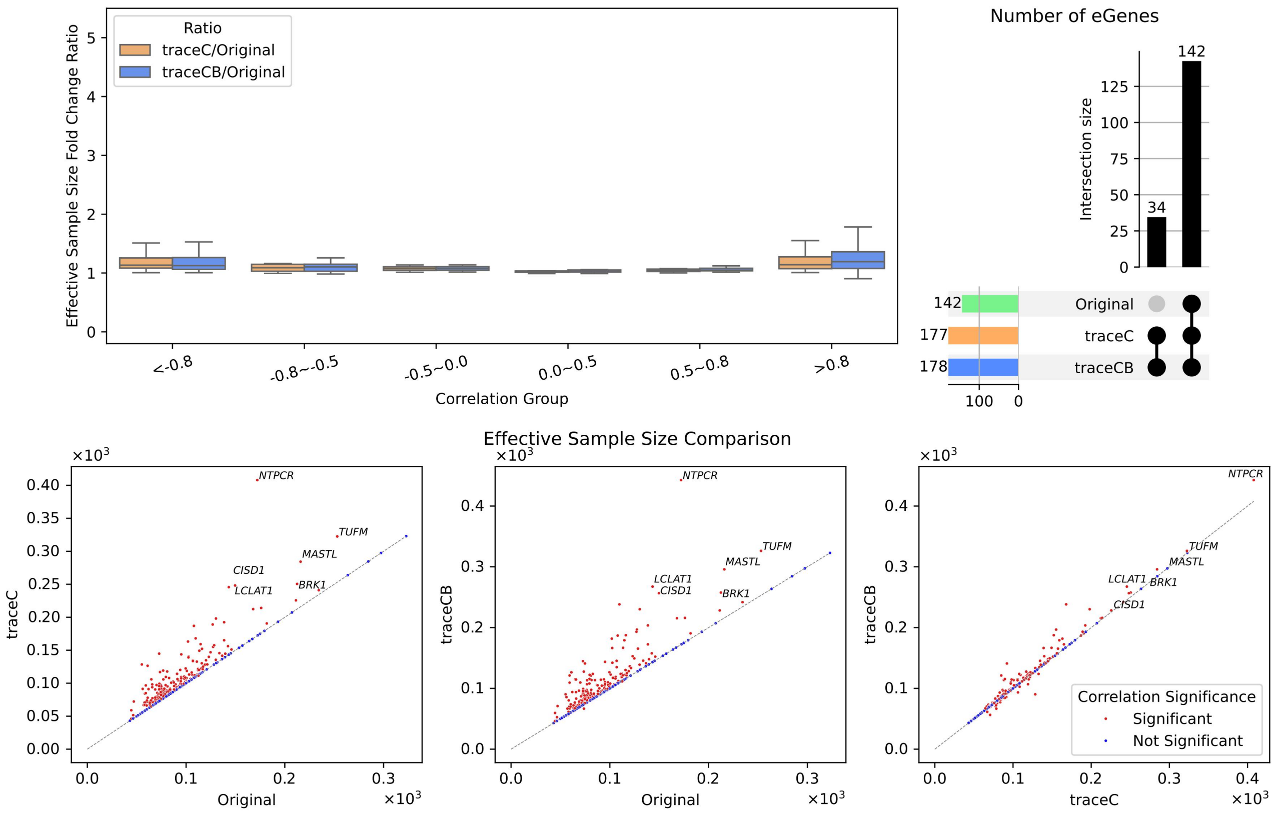

**Supplementary Figure 51:** Case study of African CD8$^{+}$ T eQTL prioritization using Kasela_2017 (269) and eQTLGen. Upper left: Effective sample size ratio comparison across genetic correlation quartiles; Upper right: Upset plots illustrate the number of eGenes identified by each analysis, and the novel eGenes discovered by traceC and traceCB. Lower: Effective sample size comparison restricted to genes with significant heritability across both ancestries. Colors represent genes with (red) and without (blue) significant trans-ancestry correlation.

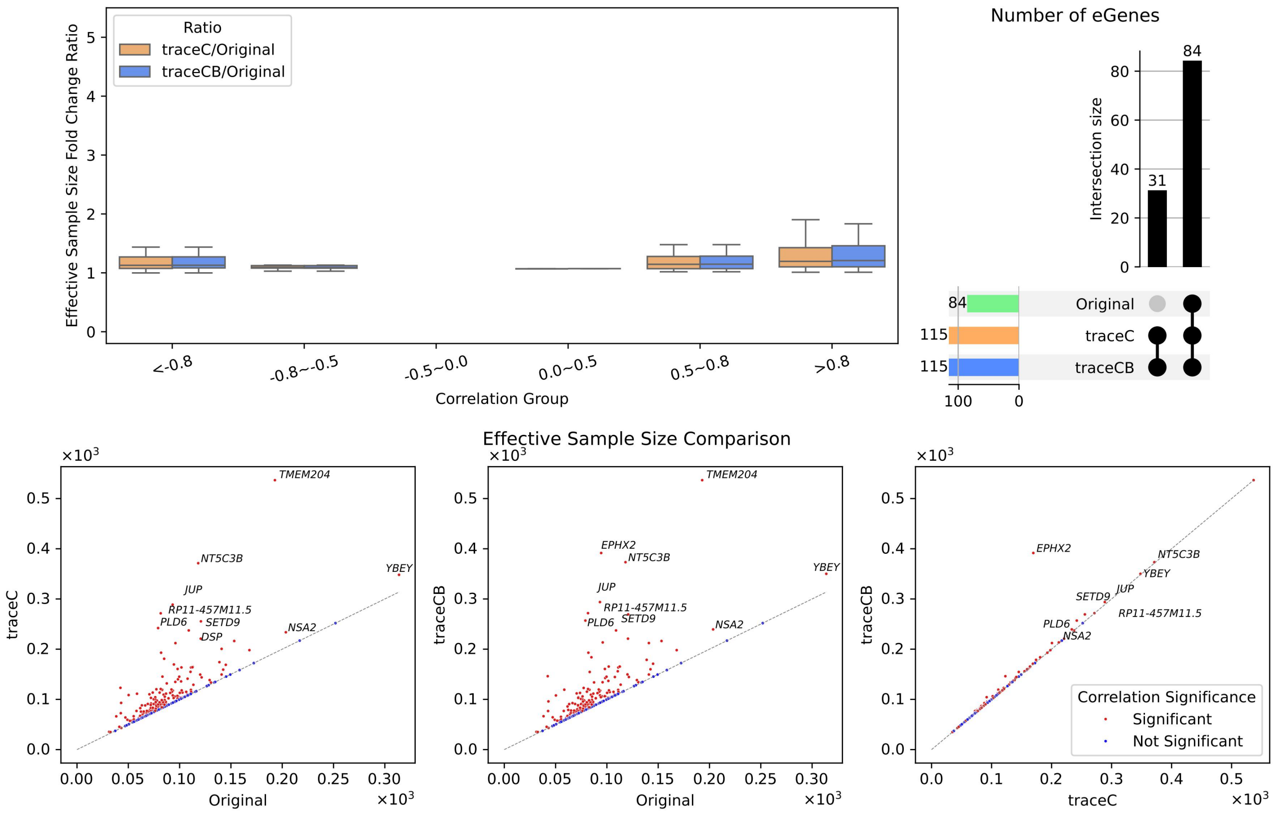

**Supplementary Figure 52:** Case study of African B cell eQTL prioritization using CEDAR (262) and eQTLGen. Upper left: Effective sample size ratio comparison across genetic correlation quartiles; Upper right: Upset plots illustrate the number of eGenes identified by each analysis, and the novel eGenes discovered by traceC and traceCB. Lower: Effective sample size comparison restricted to genes with significant heritability across both ancestries. Colors represent genes with (red) and without (blue) significant trans-ancestry correlation.

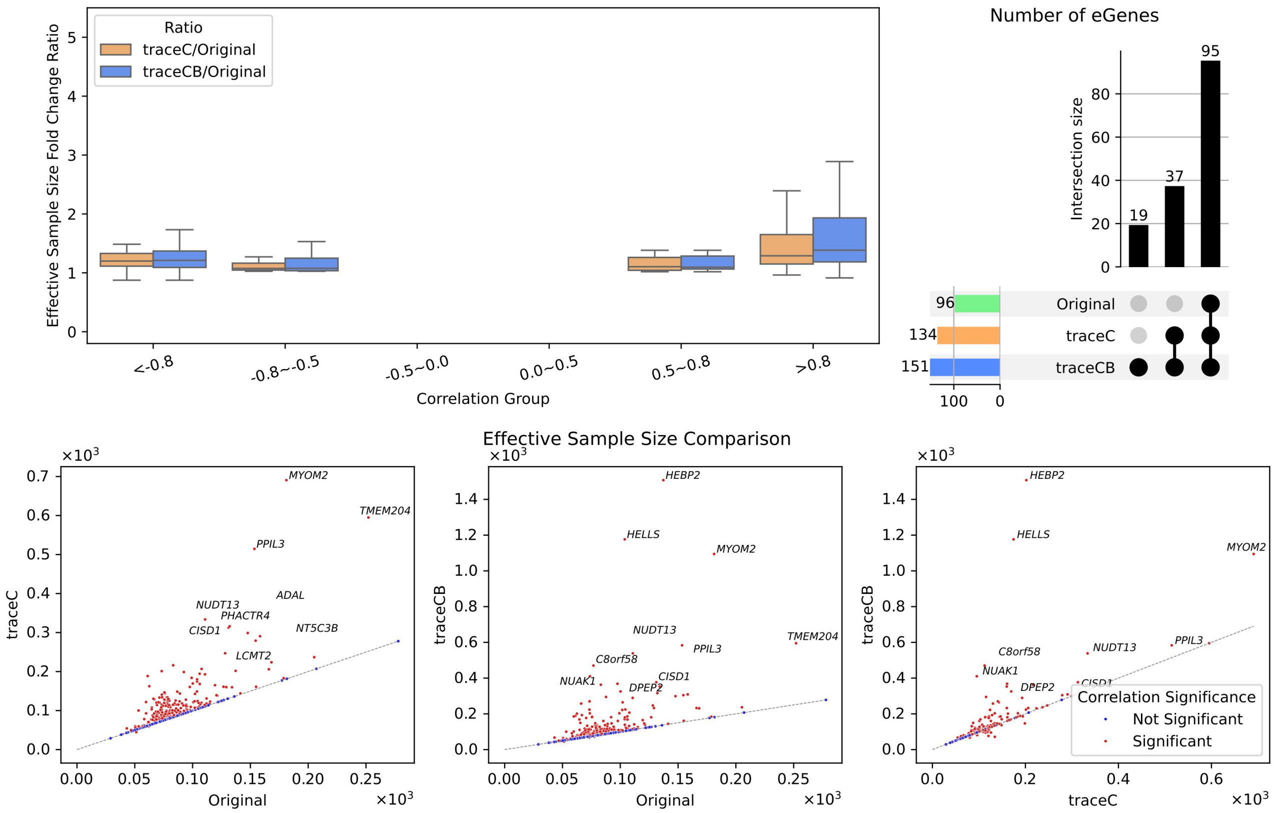

**Supplementary Figure 53:** Case study of African NK cell eQTL prioritization using Gilchrist_2021 (247) and eQTLGen. Upper left: Effective sample size ratio comparison across genetic correlation quartiles; Upper right: Upset plots illustrate the number of eGenes identified by each analysis, and the novel eGenes discovered by traceC and traceCB. Lower: Effective sample size comparison restricted to genes with significant heritability across both ancestries. Colors represent genes with (red) and without (blue) significant trans-ancestry correlation.

### 2 Supplementary Tables

**Supplementary Table 1:** Summary of data sources utilized in this study, along with their corresponding ancestries, sources, and access links. AFR: African ancestry.

| Category | Ancestry | Data Name | ID/Description | Link |
|---|---|---|---|---|
| ct-eQTL summary statistics | EAS | BBJ eQTL | | http://jenger.riken.jp/result |
| | | OASIS | $CD4^+$ T cells; $CD8^+$ T cells; B cells; NK cells; Monocytes | https://humandbs.biosciencedbc.jp/en/hum0197-latest |
| | EUR | Onek1k | | https://onek1k.org/ |
| | | eQTL Catalogue | QTD000021, QTD000031, QTD000066, QTD000067, QTD000069, QTD000073, QTD000081, QTD000115, QTD000371, QTD000372 | https://www.ebi.ac.uk/eqtl/ |
| Tissue eQTL summary statistics | AFR | Popcell | 80 Africans+non-stimulated condition | https://dataset.owey.io/doi/10.48802/owey.e4qn-9190 |
| | EUR | GTEx eQTL | GTEx_Analysis_v8_QTLs-GTEx_Analysis_ v8_eQTL_all_associations-Whole_Blood.allpairs | https://www.gtexportal.org |
| | | eQTLGen | Full cis-eQTL summary statistics | https://eqtlgen.org/cis-eqtls.html |
| | EAS | Wang et al. | hum0343.v3.eqtl.v1 | https://humandbs.dbcls.jp/en/hum0343-v4 |
| GWAS summary statistics | EAS | BBJ GWAS | Asthma, Atopic dermatitis, Rheumatoid arthritis | http://jenger.riken.jp/result |
| | | Blood Cell Consortium | East-Asian: Red blood cells, White blood cells, Platelets | http://www.mhi-humangenetics.org/en/resources/ |
| Individual-level bulk gene expression | EUR | GTEx RNA-Seq | GTEx_Analysis_v8_QTLs-GTEx_Analysis_ v8_eQTL_all_associations-Whole_Blood.allpairs | https://www.gtexportal.org |
| CIBERSORTx Reference | | GEPIA2021 | LM22 | https://github.com/zwj-tina/GEPIA2021/blob/main/reference/LM22.txt |
| LD Reference Panel | | 1000 Genomes | | https://www.cog-genomics.org/plink/1.9/resources |
| H3K27ac ChIP-seq signal | EUR | ENCODE | $CD4^+$ T cells (ENCFF357NOB), $CD8^+$ T cells (ENCFF455UVQ), B cells (ENCFF701BIL), NK cells (ENCFF473CXT), and monocytes (ENCFF840HBF) | https://www.encodeproject.org/ |

**Supplementary Table 2:** Summary of the BBJ cell-type-specific eQTL datasets. For each immune cell type, the table details the sample size, the number of genes analyzed, the total count of SNPs with available RSIDs, and the average number of SNPs per gene.

| Cell Type | Sample Size | Gene Count | SNP Count | SNP per Gene |
|---|---|---|---|---|
| $CD4^+$ T cells | 103 | 20,108 | 4,670,839 | 3,742.81 |
| $CD8^+$ T cells | 103 | 19,814 | 4,654,813 | 3,746.10 |
| B cells | 104 | 20,203 | 4,695,082 | 3,737.40 |
| NK cells | 104 | 19,919 | 4,702,533 | 3,749.02 |
| Monocytes | 105 | 19,139 | 4,661,682 | 3,750.35 |
| Average | 103.8 | 19,836.6 | 4,676,989.8 | 3,745.14 |

**Supplementary Table 3:** Summary of the 10 auxiliary studies from the eQTL Catalogue utilized in the primary analyses. This table provides detailed information for each study, including the cell type, dataset ID, quantification method, and the number of genes and SNPs. The sample size for each study is indicated in parentheses.

| Auxiliary Study Label | Cell Type | Dataset ID | Quant Method | Gene Count | SNP Count | SNP per Gene |
|---|---|---|---|---|---|---|
| BLUEPRINT (191) | Monocytes | QTD000021 | bulk RNA-seq | 16,060 | 7,133,403 | 5,823.83 |
| BLUEPRINT (167) | $CD4^+$ T | QTD000031 | bulk RNA-seq | 17,071 | 7,242,793 | 5,909.54 |
| CEDAR (277) | $CD8^+$ T | QTD000066 | microarray | 19,541 | 7,879,923 | 5,793.08 |
| CEDAR (290) | $CD4^+$ T | QTD000067 | microarray | 19,541 | 7,915,119 | 5,818.81 |
| CEDAR (286) | Monocytes | QTD000069 | microarray | 19,541 | 7,904,587 | 5,811.36 |
| CEDAR (262) | B | QTD000073 | microarray | 19,538 | 7,831,977 | 5,758.30 |
| Fairfax_2014 (420) | Monocytes | QTD000081 | microarray | 19,536 | 7,865,841 | 5,768.65 |
| Gilchrist_2021 (247) | NK | QTD000115 | microarray | 19,533 | 7,807,949 | 5,728.62 |
| Kasela_2017 (280) | $CD4^+$ T | QTD000371 | microarray | 19,558 | 7,893,389 | 5,814.62 |
| Kasela_2017 (269) | $CD8^+$ T | QTD000372 | microarray | 19,558 | 7,862,519 | 5,791.46 |
| Average | — | — | — | 18,948 | 7,733,750 | 5,801.83 |

**Supplementary Table 4:** Summary of the African ct-eQTL, eQTLGen, and GTEx whole blood eQTL datasets. For each immune cell type, the table details the sample size, the number of genes analyzed, the total count of SNPs with available RSIDs, and the average number of SNPs per gene.

| Data Type | Sample Size | Gene Count | SNP Count | SNP per Gene |
|---|---|---|---|---|
| eQTLGen | 31,684 | 19,250 | 8,932,843 | 6,615.16 |
| GTEx | 670 | 19,696 | 8,780,381 | 6,923.72 |
| African | 80 | 12,114 | 1,541,283 | 1,279.43 |

**Supplementary Table 5:** Estimated averaged proportions of immune cell types in GTEx whole blood, derived from individual-level gene expression data using the CIBERSORTx deconvolution algorithm.

| Cell type | B cells | $CD4^+$ T | $CD8^+$ T | NK cells | Monocytes |
|---|---|---|---|---|---|
| Proportion (%) | 0.806900174 | 10.4011658 | 7.140400706 | 9.722658552 | 14.75536109 |

**Supplementary Table 6:** Summary of the intersecting datasets for the three primary analytical configurations. This table details the number of common genes, SNPs, and the average number of SNPs per gene after combining the target population data (EAS or AFR) with the respective auxiliary tissue data (GTEx or eQTLGen) and for each of the 10 cell-type-specific auxiliary studies.

| Target Study_Auxiliary Tissue | Auxiliary Study | Gene Count | SNP Count | SNP per Gene |
|---|---|---|---|---|
| AFR_eQTLGen | BLUEPRINT (191) | 10,476 | 1,104,929 | 860.59 |
| | CEDAR (277) | 10,455 | 1,134,315 | 878.47 |
| | CEDAR (286) | 10,455 | 1,134,951 | 878.97 |
| | CEDAR (262) | 10,455 | 1,132,829 | 877.27 |
| | Gilchrist_2021 (247) | 10,453 | 1,129,982 | 874.73 |
| | Kasela_2017 (269) | 10,455 | 1,121,639 | 869.19 |
| | BLUEPRINT (167) | 10,667 | 1,107,309 | 862.38 |
| | CEDAR (290) | 10,455 | 1,135,209 | 879.15 |
| | Fairfax_2014 (420) | 10,455 | 1,132,004 | 876.18 |
| | Kasela_2017 (280) | 10,455 | 1,122,725 | 870.02 |
| EAS_GTEx | BLUEPRINT (191) | 12,538 | 3,052,080 | 2,453.29 |
| | CEDAR (277) | 12,505 | 3,096,389 | 2,488.17 |
| | CEDAR (286) | 12,471 | 3,108,792 | 2,488.56 |
| | CEDAR (262) | 12,506 | 3,101,876 | 2,483.79 |
| | Gilchrist_2021 (247) | 12,684 | 3,111,779 | 2,486.14 |
| | Kasela_2017 (269) | 12,505 | 3,093,671 | 2,486.45 |
| | BLUEPRINT (167) | 12,840 | 3,047,088 | 2,455.64 |
| | CEDAR (290) | 12,549 | 3,099,107 | 2,489.57 |
| | Fairfax_2014 (420) | 12,471 | 3,107,643 | 2,486.65 |
| | Kasela_2017 (280) | 12,549 | 3,097,006 | 2,488.63 |
| EAS_eQTLGen | BLUEPRINT (191) | 12,270 | 3,126,948 | 2,483.09 |
| | CEDAR (277) | 12,701 | 3,188,964 | 2,519.00 |
| | CEDAR (286) | 12,547 | 3,193,578 | 2,518.23 |
| | CEDAR (262) | 12,675 | 3,191,808 | 2,512.62 |
| | Gilchrist_2021 (247) | 12,834 | 3,200,447 | 2,514.95 |
| | Kasela_2017 (269) | 12,701 | 3,192,922 | 2,521.65 |
| | BLUEPRINT (167) | 12,774 | 3,123,222 | 2,486.62 |
| | CEDAR (290) | 12,770 | 3,191,622 | 2,520.46 |
| | Fairfax_2014 (420) | 12,547 | 3,192,502 | 2,516.34 |
| | Kasela_2017 (280) | 12,770 | 3,196,148 | 2,523.91 |

**Supplementary Table 7:** Comparison of effective sample sizes for BBJ eQTL prioritization, contrasting the use of eQTLGen and GTEx as auxiliary tissue data. The table presents the mean effective sample sizes for both traceC and traceCB across 10 cell-type-specific auxiliary studies. The sample size for each auxiliary study is indicated in parentheses.

| Auxiliary Study | eQTLGen | | GTEx | |
|---|---|---|---|---|
| Method | TraceC | TraceCB | TraceC | TraceCB |
| BLUEPRINT (167) | 212.92 | 258.13 | 213.00 | 220.45 |
| BLUEPRINT (191) | 220.52 | 286.69 | 222.27 | 235.34 |
| CEDAR (262) | 172.92 | 176.8 | 174.30 | 174.74 |
| CEDAR (277) | 179.53 | 225.28 | 180.78 | 184.72 |
| CEDAR (286) | 180.12 | 247.85 | 179.41 | 186.96 |
| CEDAR (290) | 183.13 | 242.58 | 184.52 | 188.14 |
| Fairfax_2014 (420) | 213.11 | 298.07 | 214.47 | 222.34 |
| Gilchrist_2021 (247) | 179.12 | 230.67 | 178.35 | 183.13 |
| Kasela_2017 (269) | 167.49 | 203.27 | 168.21 | 171.71 |
| Kasela_2017 (280) | 169.5 | 223.08 | 170.39 | 173.90 |
| Average | 187.84 | 239.24 | 188.57 | 194.14 |

**Supplementary Table 8:** Evaluation of eGene replication in an external East Asian bulk tissue dataset (OASIS). The table presents the number of identified eGenes, the number of replicated eGenes, and the corresponding replication rates for the original BBJ eQTL analysis, traceC, and traceCB. The analysis was conducted using eQTLGen as the auxiliary tissue data.

| | Number of eGenes identified | | | Number of eGenes replicated | | | Replication rate | | |
|---|---|---|---|---|---|---|---|---|---|
| Auxiliary study | Original | traceC | traceCB | Original | traceC | traceCB | Original | traceC | traceCB |
| BLUEPRINT (167) | 1985 | 2955 | 3176 | 1825 | 2678 | 2861 | 0.92 | 0.91 | 0.90 |
| BLUEPRINT (191) | 2385 | 3441 | 3750 | 2282 | 3251 | 3503 | 0.96 | 0.94 | 0.93 |
| CEDAR (262) | 1721 | 1988 | 1998 | 1589 | 1825 | 1831 | 0.92 | 0.92 | 0.92 |
| CEDAR (277) | 1623 | 1987 | 2163 | 1508 | 1845 | 1996 | 0.93 | 0.93 | 0.92 |
| CEDAR (286) | 2313 | 2717 | 2974 | 2198 | 2561 | 2778 | 0.95 | 0.94 | 0.93 |
| CEDAR (290) | 1860 | 2296 | 2516 | 1718 | 2106 | 2291 | 0.92 | 0.92 | 0.91 |
| Fairfax_2014 (420) | 2308 | 2996 | 3286 | 2193 | 2820 | 3062 | 0.95 | 0.94 | 0.93 |
| Gilchrist_2021 (247) | 1606 | 2131 | 2360 | 1490 | 1966 | 2166 | 0.93 | 0.92 | 0.92 |
| Kasela_2017 (269) | 1628 | 1912 | 2067 | 1512 | 1777 | 1913 | 0.93 | 0.93 | 0.93 |
| Kasela_2017 (280) | 1860 | 2211 | 2397 | 1719 | 2037 | 2201 | 0.92 | 0.92 | 0.92 |
| Average | 1928.9 | 2463.4 | 2668.7 | 1803.4 | 2286.6 | 2460.2 | 0.93 | 0.93 | 0.92 |

**Supplementary Table 9:** Evaluation of eGene replication in an external East Asian bulk tissue dataset (OASIS). The table presents the number of identified eGenes, the number of replicated eGenes, and the corresponding replication rates for the original BBJ eQTL analysis, traceC, and traceCB. The analysis was conducted using GTEx whole blood as the auxiliary tissue.

| | Number of eGenes Identified | | | Number of eGenes Replicated | | | Replication Rate | | |
|---|---|---|---|---|---|---|---|---|---|
| Auxiliary study | Original | traceC | traceCB | Original | traceC | traceCB | Original | traceC | traceCB |
| BLUEPRINT (167) | 1965 | 2925 | 2952 | 1834 | 2692 | 2717 | 0.93 | 0.92 | 0.92 |
| BLUEPRINT (191) | 2401 | 3466 | 3559 | 2312 | 3294 | 3374 | 0.96 | 0.95 | 0.95 |
| CEDAR (262) | 1685 | 1956 | 1956 | 1577 | 1819 | 1819 | 0.94 | 0.93 | 0.93 |
| CEDAR (277) | 1593 | 1946 | 1960 | 1499 | 1825 | 1840 | 0.94 | 0.94 | 0.94 |
| CEDAR (286) | 2287 | 2677 | 2724 | 2194 | 2550 | 2594 | 0.96 | 0.95 | 0.95 |
| CEDAR (290) | 1809 | 2244 | 2267 | 1699 | 2091 | 2113 | 0.94 | 0.93 | 0.93 |
| Fairfax_2014 (420) | 2282 | 2956 | 3011 | 2189 | 2806 | 2857 | 0.96 | 0.95 | 0.95 |
| Gilchrist_2021 (247) | 1573 | 2097 | 2120 | 1482 | 1958 | 1979 | 0.94 | 0.93 | 0.93 |
| Kasela_2017 (269) | 1597 | 1879 | 1898 | 1502 | 1768 | 1787 | 0.94 | 0.94 | 0.94 |
| Kasela_2017 (280) | 1810 | 2149 | 2174 | 1700 | 2009 | 2032 | 0.94 | 0.93 | 0.93 |
| Average | 1900.2 | 2429.5 | 2462.1 | 1798.8 | 2281.2 | 2311.2 | 0.95 | 0.94 | 0.94 |

**Supplementary Table 10:** Effective sample size and number of identified eGenes in the African ancestry eQTL analysis. Results are shown for prioritization using 10 different European auxiliary studies, with eQTLGen as the auxiliary tissue data. The number in parentheses indicates the sample size of each auxiliary study.

| | Effective Sample Size | | Number of eGene Identified | | |
|---|---|---|---|---|---|
| Auxiliary study | traceC | traceCB | Original | traceC | traceCB |
| BLUEPRINT (167) | 123.88 | 139.65 | 558 | 646 | 655 |
| BLUEPRINT (191) | 127.79 | 144.70 | 632 | 740 | 750 |
| CEDAR (262) | 116.77 | 119.40 | 270 | 301 | 301 |
| CEDAR (277) | 121.52 | 137.11 | 496 | 537 | 545 |
| CEDAR (286) | 116.92 | 142.32 | 618 | 676 | 694 |
| CEDAR (290) | 120.85 | 139.32 | 526 | 567 | 578 |
| Fairfax_2014 (420) | 123.14 | 143.40 | 622 | 718 | 740 |
| Gilchrist_2021 (247) | 123.72 | 153.95 | 285 | 323 | 340 |
| Kasela_2017 (269) | 110.31 | 113.85 | 492 | 527 | 528 |
| Kasela_2017 (280) | 113.32 | 125.63 | 524 | 556 | 561 |
| Average | 119.82 | 135.93 | 502.3 | 559.1 | 569.2 |

### 3 Supplementary Notes

#### 3.1 Approximation of tissue-level marginal effects using mean cell-type proportions

In this section, we provide a detailed derivation to justify the approximation of the true marginal effect size in bulk tissue $b_{tj}$, using the mean cell-type proportions $\bar{\pi}_c$ and $\bar{\pi}_o$ instead of individual-level proportions $\boldsymbol{\pi}_c$ and $\boldsymbol{\pi}_o$. The estimated marginal effect size for SNP $j$ in the bulk tissue is given by:

$$\begin{aligned}
\hat{b}_{tj} &= \frac{\mathbf{x}_{tj}^T \mathbf{y}_t}{\mathbf{x}_{tj}^T \mathbf{x}_{tj}} \\
&= \frac{1}{N_t} \mathbf{x}_{tj}^T \mathbf{y}_t \\
&= \frac{1}{N_t} \mathbf{x}_{tj}^T (\boldsymbol{\pi}_c \odot \mathbf{X}_t \boldsymbol{\beta}_{c2} + \boldsymbol{\pi}_o \odot \mathbf{X}_t \boldsymbol{\beta}_{o2} + \boldsymbol{\epsilon}_t) \\
&= \frac{1}{N_t} \sum_{i=1}^{N_t} x_{tj,i} (\pi_{ci} \mathbf{x}_{t,i}^\top \boldsymbol{\beta}_{c2} + \pi_{oi} \mathbf{x}_{t,i}^\top \boldsymbol{\beta}_{o2} + \epsilon_{t,i}),
\end{aligned}$$

where $x_{tj,i}$ is the $i$-th individual's genotype at SNP $j$, $\mathbf{x}_{t,i}$ is the genotype vector of individual $i$ across all $M$ SNPs, and $\epsilon_{t,i}$ is the residual error for individual $i$ in tissue. The notation $\odot$ denotes element-wise multiplication.

A key challenge in relating the tissue-level effect $b_{tj}$ to cell-type-level effects is that the individual-level cell-type proportions $\pi_{ci}$ and $\pi_{oi}$, vary across samples. However, we can show that for a sufficiently large sample size $N_t$, $\hat{b}_{tj}$ can be approximated using the mean proportions

$\bar{\pi}_c$ and $\bar{\pi}_o$ across the cohort. The derivation proceeds as follows:

$$\begin{aligned}
\hat{b}_{tj} &= \frac{1}{N_t}\sum_{i=1}^{N_t} x_{tj,i}\left[\pi_{ci}\left(x_{tj,i}\beta_{c2j} + \sum_{k\neq j} x_{tk,i}\beta_{c2k}\right) + \pi_{oi}\left(x_{tj,i}\beta_{o2j} + \sum_{k\neq j} x_{tk,i}\beta_{o2k}\right) + \epsilon_{t,i}\right] \\
&\overset{(1)}{\approx} \frac{1}{N_t}\sum_{i=1}^{N_t} x_{tj,i}(\pi_{ci}x_{tj,i}\beta_{c2j} + \pi_{oi}x_{tj,i}\beta_{o2j}) + \text{constant} \\
&= \frac{1}{N_t}\sum_{i=1}^{N_t}(\pi_{ci}x_{tj,i}^2\beta_{c2j} + \pi_{oi}x_{tj,i}^2\beta_{o2j}) + \text{constant} \\
&= \frac{1}{N_t}\sum_{g\in\{0,1,2\}}\sum_{i:g_{tj,i}=g}(\pi_{ci}x_{tj,i}^2\beta_{c2j} + \pi_{oi}x_{tj,i}^2\beta_{o2j}) + \text{constant} \\
&\overset{(2)}{\approx} \frac{1}{N_t}\sum_{g\in\{0,1,2\}}\sum_{i:g_{tj,i}=g}(\bar{\pi}_c x_{tj,i}^2\beta_{c2j} + \bar{\pi}_o x_{tj,i}^2\beta_{o2j}) + \text{constant} \\
&= \frac{1}{N_t}\sum_{i=1}^{N_t}(\bar{\pi}_c x_{tj,i}^2\beta_{c2j} + \bar{\pi}_o x_{tj,i}^2\beta_{o2j}) + \text{constant} \\
&\overset{(3)}{\approx} \frac{1}{N_t}\sum_{i=1}^{N_t} x_{tj,i}(\bar{\pi}_c \mathbf{x}_{t,i}^\top \boldsymbol{\beta}_{c2} + \bar{\pi}_o \mathbf{x}_{t,i}^\top \boldsymbol{\beta}_{o2} + \epsilon_{t,i}) \\
&= \frac{1}{N_t}\mathbf{x}_{tj}^\top(\bar{\pi}_c \odot \mathbf{X}_t\boldsymbol{\beta}_{c2} + \bar{\pi}_o \odot \mathbf{X}_t\boldsymbol{\beta}_{o2} + \boldsymbol{\epsilon}_t),
\end{aligned}$$

where $g_{tj,i}$ is the raw genotype corresponding to the standardized genotype $x_{tj,i}$. In approximation (1), we isolate the contribution of SNP $j$'s causal effect, treating the aggregate effects from other SNPs in LD as a constant term. In approximation (2), we apply the law of large numbers: for a sufficiently large sample size $N_t$, the number of individuals $N_g$ in each genotype stratum $g$ will also be large. Consequently, the average of individual cell-type proportions with the same genotype value, $\frac{1}{N_g}\sum_{i:g_{tj,i}=g}\pi_{ci}$, converges to the mean proportion $\bar{\pi}_c$. The approximation (3) repeats the procedure of approximation (1). This derivation demonstrates that $\hat{b}_{tj}$ can be reasonably approximated using mean cell-type proportions.

#### 3.2 Derivation of traceCB estimator

We apply the generalized method of moments (GMM) to obtain the traceCB estimator. To derive the conditional mean and variance of the observed marginal effects $\hat{\boldsymbol{b}}_j = [\hat{b}_{1j}, \hat{b}_{2j}, \hat{b}_{tj}]^\top$ given the true effect size $b_{1j}$, we start by establishing the relationship between the observed effects and the true effects using a linear projection. The projection of $\hat{b}_{2j}$ onto $b_{1j}$ is given as:

$$\mathbb{E}\left[\hat{b}_{2j} - \lambda b_{1j}\right] = 0,$$

where $\lambda$ is a scalar parameter representing the linear projection coefficient. We solve for $\lambda$ by minimizing the quadratic objective function:

$$\min_{\lambda} \mathbb{E}\left[\left(\hat{b}_{2j} - \lambda b_{1j}\right)^2\right].$$

Taking the derivative with respect to $\lambda$ and setting it to zero yields:

$$\begin{aligned}
0 &= \frac{\partial}{\partial \lambda}\mathbb{E}\left[\left(\hat{b}_{2j} - \lambda b_{1j}\right)^2\right] \\
&= 2\mathbb{E}\left[\left(\hat{b}_{2j} - \lambda b_{1j}\right)(-b_{1j})\right] \\
&= -2\mathbb{E}\left[b_{1,j}\left(b_{2,j} + e_{2,j} - \lambda b_{1,j}\right)\right] \\
&= -2\left(\mathbb{E}\left[b_{1,j} b_{2,j}\right] - \lambda \mathbb{E}\left[b_{1,j}^2\right]\right) \\
&= -2\left(\omega_x - \lambda \omega_1\right).
\end{aligned}$$

Then, we obtain the estimator $\hat{\lambda} = \omega_x/\omega_1$. Similarly, for the moment condition $\mathbb{E}[(\hat{b}_{t,j} - \lambda_t b_{1,j})] = 0$, we can derive $\hat{\lambda}_t = \bar{\pi}_c \omega_x/\omega_1$.

We can extend these scalar relationships to a vector form. The moment equation for the vector of observed effects $\hat{\boldsymbol{b}}_j$ is defined as $\mathbf{m}(b) := \hat{\boldsymbol{b}}_j - \boldsymbol{\lambda}_1 b$, leading to the first-order moment condition presented in the main text:

$$\mathbb{E}[\mathbf{m}(b)|_{b=b_{1j}}] = \mathbb{E}\left(\hat{\boldsymbol{b}}_j - \boldsymbol{\lambda}_1 b_{1j}\right) = \mathbf{0},$$

where $\boldsymbol{\lambda}_1 = [\omega_1, \omega_x, \bar{\pi}_c \omega_x]^\top/\omega_1$.

From this moment condition, the law of total expectation implies $\mathbb{E}[\mathbb{E}(\hat{\boldsymbol{b}}_j - \boldsymbol{\lambda}_1 b_{1,j} | b_{1,j})] = \mathbf{0}$. This allows us to derive the conditional mean of $\hat{\boldsymbol{b}}_j$ given $b_{1,j}$:

$$\mathbb{E}(\hat{\boldsymbol{b}}_j | b_{1,j}) = \mathbb{E}(\boldsymbol{\lambda}_1 b_{1,j} | b_{1,j}) = \boldsymbol{\lambda}_1 b_{1,j}.$$

Using the law of total variance, the conditional variance is derived as:

$$\begin{aligned}
\operatorname{Var}(\hat{\boldsymbol{b}}_j | b_{1,j}) &= \operatorname{Var}(\hat{\boldsymbol{b}}_j) - \operatorname{Var}(\mathbb{E}(\hat{\boldsymbol{b}}_j | b_{1,j})) \\
&= \left(\mathbf{A}\boldsymbol{\Omega}_j \mathbf{A}^T + \boldsymbol{\Sigma}_{oj} + \hat{\mathbf{S}}_j \mathbf{C} \hat{\mathbf{S}}_j\right) - \operatorname{Var}(\boldsymbol{\lambda}_1 b_{1,j}) \\
&= \mathbf{A}\boldsymbol{\Omega}_j \mathbf{A}^T + \boldsymbol{\Sigma}_{oj} + \hat{\mathbf{S}}_j \mathbf{C} \hat{\mathbf{S}}_j - \omega_1 \boldsymbol{\lambda}_1 \boldsymbol{\lambda}_1^T \\
&:= \boldsymbol{\Lambda}_{1j}^{-1}.
\end{aligned}$$

#### 3.3 Parameter estimation with two correlated cell types in bulk tissue

To estimate the unknown parameters $\boldsymbol{\Omega}$, $\omega_o$ and $\mathbf{C}$ from Equation (15), we perform a series of regressions based on the Z scores. The slopes of these regressions yield estimates for the per-SNP heritabilities ($\omega_1, \omega_2, \omega_o$) and co-heritability ($\omega_x$), while the intercepts provide estimates for the inflation parameters $(c_1, c_2, c_t, c_x)$. The regression models are specified as follows:

$$
\begin{aligned}
\frac{\hat{b}_{1j}^2}{\hat{s}_{1j}^2} &\approx \omega_1 \frac{l_{1j}}{\hat{s}_{1j}^2} + c_1, \\
\frac{\hat{b}_{2j}^2}{\hat{s}_{2j}^2} &\approx \omega_2 \frac{l_{2j}}{\hat{s}_{2j}^2} + c_2, \\
\frac{\hat{b}_{1j}\hat{b}_{2j}}{\hat{s}_{1j}\hat{s}_{2j}} &\approx \omega_x \frac{l_{xj}}{\hat{s}_{1j}\hat{s}_{2j}} + c_x, \\
\frac{\hat{b}_{tj}^2}{\hat{s}_{tj}^2} &\approx \left(\bar{\pi}_c^2\omega_2 + \bar{\pi}_o^2\omega_o\right) \frac{l_{2j}}{\hat{s}_{tj}^2} + c_t,
\end{aligned}
$$

We first obtain $\hat{\omega}_1, \hat{\omega}_2$, and $\hat{\omega}_x$ from the corresponding coefficients of the first three regression equations. Given $\bar{\pi}_c, \bar{\pi}_o$ and $\hat{\omega}_2$, $\hat{\omega}_o$ can be calculated based on the slope of the last regression equation. The LD scores $(l_{1j}, l_{2j}, l_{xj})$ can be pre-computed from external reference panels, such as the 1000 Genomes Project, matched to the ancestry of the respective populations.

Following the standard LDSC [1], we employ a weighted least squares estimator to account for the dependency among Z scores. To mitigate the influence of outlier SNPs with large effects, we adopt a robust two-step estimation procedure. In the first step, we estimate the intercepts $(c_1, c_2, c_x, c_t)$ by excluding SNPs with high association signals (e.g., $\chi^2 > 30$). In the second step, holding the intercepts fixed at their estimated values, we estimate the heritability and co-heritability parameters $(\omega_1, \omega_2, \omega_x, \omega_o)$ using the full set of SNPs. In the absence of population structure and sample overlap, these confounding factors can be set to their theoretical values, e.g. $\hat{\mathbf{C}} = \mathbf{I}$. By plugging in the estimated parameters, we can construct the estimated matrices $\hat{\mathbf{\Omega}}_j$ and $\hat{\mathbf{C}}$.

In the simulation and real data analyses conducted in our study, we set the confounding factors matrix $\mathbf{C}$ to the identity matrix $\mathbf{I}$, as no significant confounding effects were observed in the datasets we analyzed. However, for scenarios where such confounding is present, we recommend estimating $\mathbf{C}$ using the LDSC-based approach described above.

##### 3.4 Parameter estimation with multiple correlated cell types in bulk tissue

When there are multiple highly correlated cell types in the tissue sample, ignoring the genetic correlation may lead to enlarged variance for traceCB's GMM estimator, restricting the power of eQTL mapping. Here, we extend the above parameter estimation procedure to accommodate multiple correlated cell types within a tissue to account for genetic correlations within non-target cell types. We consider a scenario where the bulk tissue comprises $K + 1$ distinct cell types: one target cell type of interest and $K$ other cell types.

Assume the proportions of non-target cell types are $[\boldsymbol{\pi}_1, \boldsymbol{\pi}_2, \ldots, \boldsymbol{\pi}_K]^\top$. The relationship between gene expression levels in bulk tissue and cell-type-specific expressions can be updated to:

$$\mathbf{y}_t = \boldsymbol{\pi}_c \odot \mathbf{X}_t \boldsymbol{\beta}_{c2} + \sum_{k=1}^{K} \boldsymbol{\pi}_k \odot \mathbf{X}_t \boldsymbol{\beta}_k + \boldsymbol{\epsilon}_t,$$

where $\boldsymbol{\beta}_k$ represents the eQTL effect sizes specific to cell type $k$. Following a similar derivation as in the two-cell-type scenario, the true marginal effect size in bulk tissue can be expressed as:

$$b_{tj} = \mathbb{E}\left[\hat{b}_{tj} \middle| \boldsymbol{\beta}_{c2}, \{\boldsymbol{\beta}_k\}_{k=1}^{K}\right] \approx \bar{\pi}_c \beta_{c2j} + \sum_{k=1}^{K} \bar{\pi}_k \beta_{kj}.$$

where $\beta_{kj}$ denotes the eQTL effect size for SNP $j$ in cell type $k$ and $\bar{\pi}_k$ is the mean proportion of cell type $k$ in the bulk tissue.

The variance of the observed marginal effects $\hat{b_{tj}}$ can be modeled similarly, incorporating the contributions from all $K$ non-target cell types. The $\boldsymbol{\Sigma}_{oj}$ in Equation (8) is modified to:

$$\boldsymbol{\Sigma}_{oj} = \begin{pmatrix} 0 & 0 & 0 \\ 0 & 0 & 0 \\ 0 & 0 & \sum_{k=1}^{K} \bar{\pi}_k^2 \omega_k l_{2j} \end{pmatrix},$$

where $\omega_k$ is the per-SNP heritability for cell type $k$.

Similarly, we use LDSC regression equations to estimate the unknown $\boldsymbol{\Sigma}_{oj}$. Then, we update the expectations in Equation (15) as follows:

$$\begin{aligned} \mathbb{E}\left[z_{2j} z_{tj}\right] &\approx \left(\bar{\pi}_c \omega_2 + \sum_{k=1}^{K} \bar{\pi}_k \omega_{2k}\right) l_{2j} / \left(\hat{s}_{2j} \hat{s}_{tj}\right) + c_{2t}, \\ \mathbb{E}\left[z_{tj}^2\right] &\approx \left(\bar{\pi}_c^2 \omega_2 + \sum_{k=1}^{K} \bar{\pi}_k^2 \omega_k + 2 \sum_{k=1}^{K} \bar{\pi}_c \bar{\pi}_k \omega_{2k} + \sum_{k=1}^{K} \sum_{k' \neq k} \bar{\pi}_k \bar{\pi}_{k'} \omega_{kk'}\right) l_{2j} / \hat{s}_{tj}^2 + c_t, \end{aligned}$$

where $\omega_{2k}$ is the co-heritability between the target cell type and cell type $k$, and $\omega_{kk'}$ is the co-heritability between cell types $k$ and $k'$.

To simplify the notation, let $S(.,.)$ denote the slope obtained from regressing the left-hand side on the right-hand side of each equation. We can then express the heritability and co-heritability estimates as:

$$\begin{aligned} S\left(z_{2j}, z_{tj}\right) &= \bar{\pi}_c \omega_2 + \sum_{k=1}^{K} \bar{\pi}_k \omega_{2k}, \\ S\left(z_{tj}, z_{tj}\right) &= \bar{\pi}_c^2 \omega_2 + \sum_{k=1}^{K} \bar{\pi}_k^2 \omega_k + 2 \sum_{k=1}^{K} \bar{\pi}_c \bar{\pi}_k \omega_{2k} + \sum_{k=1}^{K} \sum_{k' \neq k} \bar{\pi}_k \bar{\pi}_{k'} \omega_{kk'}. \end{aligned}$$

We assume the co-heritability between cell types in tissue are approximately equal, i.e., $\omega_{kk'} \approx \bar{\omega}_{2k}$ for all $k \neq k'$. Therefore, the above equations can be simplified to:

$$\begin{aligned} & S\left(z_{2j}, z_{tj}\right) \approx \bar{\pi}_c \omega_2 + \left(1 - \bar{\pi}_c\right) \bar{\omega}_{2k}, \\ & S\left(z_{tj}, z_{tj}\right) \approx \bar{\pi}_c^2 \omega_2 + \sum_{k=1}^{K} \bar{\pi}_k^2 \omega_k + 2 \bar{\pi}_c \left(S\left(z_{2j}, z_{tj}\right) - \bar{\pi}_c \omega_2\right) + \sum_{k=1}^{K} \sum_{k' \neq k} \bar{\pi}_k \bar{\pi}_{k'} \bar{\omega}_{2k}. \end{aligned}$$

Finally, we can estimate $\mathbf{\Sigma}_{oj}$ by:

$$\begin{aligned}\mathbf{\Sigma}_{oj}/l_{2j} &= \sum_{k=1}^{K} \bar{\pi}_k^2 \omega_k \\ &= S\left(z_{tj}, z_{tj}\right) - \bar{\pi}_c^2 \omega_2 - \left(2\bar{\pi}_c + \frac{\sum_{k=1}^{K} \sum_{k' \neq k} \bar{\pi}_k \bar{\pi}_{k'}}{1 - \bar{\pi}_c}\right) \left(S\left(z_{2j}, z_{tj}\right) - \bar{\pi}_c \omega_2\right).\end{aligned}$$
